# Replicate avian hybrid zones reveal the progression of genetic and trait introgression through time

**DOI:** 10.1101/2025.08.07.669146

**Authors:** María Isabel Castaño, Elizabeth Croyle, Carlos Daniel Cadena, J. Albert C. Uy

## Abstract

Replicate hybrid zones between the same taxa provide a unique opportunity to assess the repeatability of interspecific matings by uncovering recurrent genomic and phenotypic introgression patterns. Replicates also facilitate exploration of the causes of temporal shifts in hybrid zone structure. Here we sampled transects along three geographically separate hybrid zones between two avian taxa –the Lemon-rumped (*Ramphocelus flammigerus icteronotus*) and Flame-rumped (*R. f. flammigerus*) Tanagers – which hybridize in low passes across the Western Andes of Colombia. We examined environmental, phenotypic and genetic variation using reflectance spectrophotometry and genotype-by-sequencing data mapped to a high quality *de novo* genome assembly, aiming to assess the repeatability and progression of introgression after hybridization. We found that all hybrid zones formed independently, showed parallel phenotypic divergence along ecological gradients, low population structure across parental ranges and similar demographic histories. Replicates also exhibited asymmetric introgression of neutral markers from the yellow *icteronotus* into the hybrid zone. However, the age of the hybrid zones differed, resulting in differences in the extent of geographic and genomic cline displacement from environmental transitions into the red *flammigerus* range. Despite heterogeneity in locus-specific introgression, the only shared introgression outliers across all hybrid zones were in a genomic region linked to plumage color. Clines for these loci were consistently narrow, suggesting a role in long-term reproductive isolation. Altogether, we showed that locus-specific introgression is largely stochastic, but the magnitude and directionality of neutral introgression can be predictable when demographic conditions are similar and for traits involved in reproductive isolation.

## INTRODUCTION

Hybrid zones offer a unique opportunity to characterize interactions between incipient species, providing insights on the processes underlying species origin and maintenance (Barton & Hewitt, 1985; Harrison, 1990). Decades of research across diverse taxa have revealed that divergence between lineages can be maintained in the face of gene flow, as long as the genomic regions contributing to assortative mating, local adaptation or hybrid disadvantage remain differentiated by selection (Gompert et al., 2017; Harrison & Larson, 2016). In addition, hybrid zones can act as conduits for the spread of adaptive traits across populations (e.g., Baiz et al., 2021; Dasmahapatra et al., 2012; Lim et al., 2024). However, most hybrid zone studies involve a single contact zone between two taxa; hence, the predictability of the outcomes of hybridization and the progression of introgression remain little understood. To this end, study systems with replicate hybrid zones allow for tests of the repeatability and progression of hybridization, revealing that selection often leads to predictable patterns of local ancestry and genomic differentiation around loci involved in reproductive isolation or adaptive divergence (e.g., Blain et al., 2025; Langdon et al., 2024; Nouhaud et al., 2022; Scordato et al., 2017; Westram et al., 2021). However, introgression patterns at neutral loci can be highly variable (McFarlane et al., 2024) and the predictability of the progression of introgression over time remains elusive.

The outcomes of hybridization can be influenced by a range of factors that vary across hybrid zones, even when the interacting taxa are the same. For instance, differences in population structure across transects due to historical divergence, drift, or local adaptation can result in distinct genetic interactions, as different lineages may carry unique incompatibilities or adaptive alleles. These differences can then lead to lineage-specific introgression patterns (Barton & De Cara, 2009; Orr & Turelli, 2001).

Adaptation to unique ecological conditions may also result in variable hybridization outcomes, as local adaptation could favor different genotypes or alter the epistatic effects of alleles in hybrids (Cutter, 2012). Given the challenges of fully controlling for these ecological and genetic variables in natural populations, studies often attribute variation in introgression patterns across replicate hybrid zones to differences in environmental context (Bolte et al., 2024; Duvernell et al., 2023; Van Riemsdijk et al., 2023) or the extent of genetic divergence between parental populations (Conte et al., 2012; Dufresnes et al., 2015; Nolte et al., 2009). Finding similarities despite this variation remains a powerful way to identify genomic regions involved in reproductive isolation (Kingston et al., 2017; Nadeau et al., 2014; Scordato et al., 2020; Simon et al., 2021). However, such scenarios are rare (Gompert et al., 2017), and heterogeneous or asymmetric introgression shaped by local conditions and stochastic processes remains common (Kingston et al., 2017; McFarlane et al., 2024; Moran et al., 2021; Vijay et al., 2016).

Examples of replicate hybrid zones typically offer snapshots of short temporal windows following secondary contact, limiting our ability to assess how hybridization unfolds over evolutionary time (Buggs, 2007; Long et al., 2024; Wang et al., 2019). Yet, timing is crucial because despite repeatability in local ancestry patterns shortly after hybridization (Langdon et al., 2024; Nouhaud et al., 2022), recombination and fluctuating selection can erode these signals and make introgression increasingly unpredictable over time. As a result, variation in hybrid zone age can also contribute to differences in observed outcomes of hybridization (Groh & Coop, 2024). In turn, this temporal variability can document different stages of the speciation continuum, examine whether genetic incompatibilities or adaptive traits are maintained, and reconstruct the progression of trait introgression through time (Beysard & Heckel, 2014; Stankowski & Ravinet, 2021; Zieliński et al., 2019).

Here we take advantage of a species complex that has established multiple hybrid zones to test the repeatability of hybridization outcomes and explore the progression of introgression through evolutionary time. The Flame-rumped and Lemon-rumped Tanagers (*Ramphocelus flammigerus*) subspecies complex is found along the Pacific Coast, and slopes of the Cauca River Valley in the Western Andes of Colombia (Fig 1). The two subspecies exhibit dramatic variation in carotenoid-based plumage color of the rump, where red plumage is the ancestral state and yellow is derived (McCoy et al., 2021).

**Figure 1.**
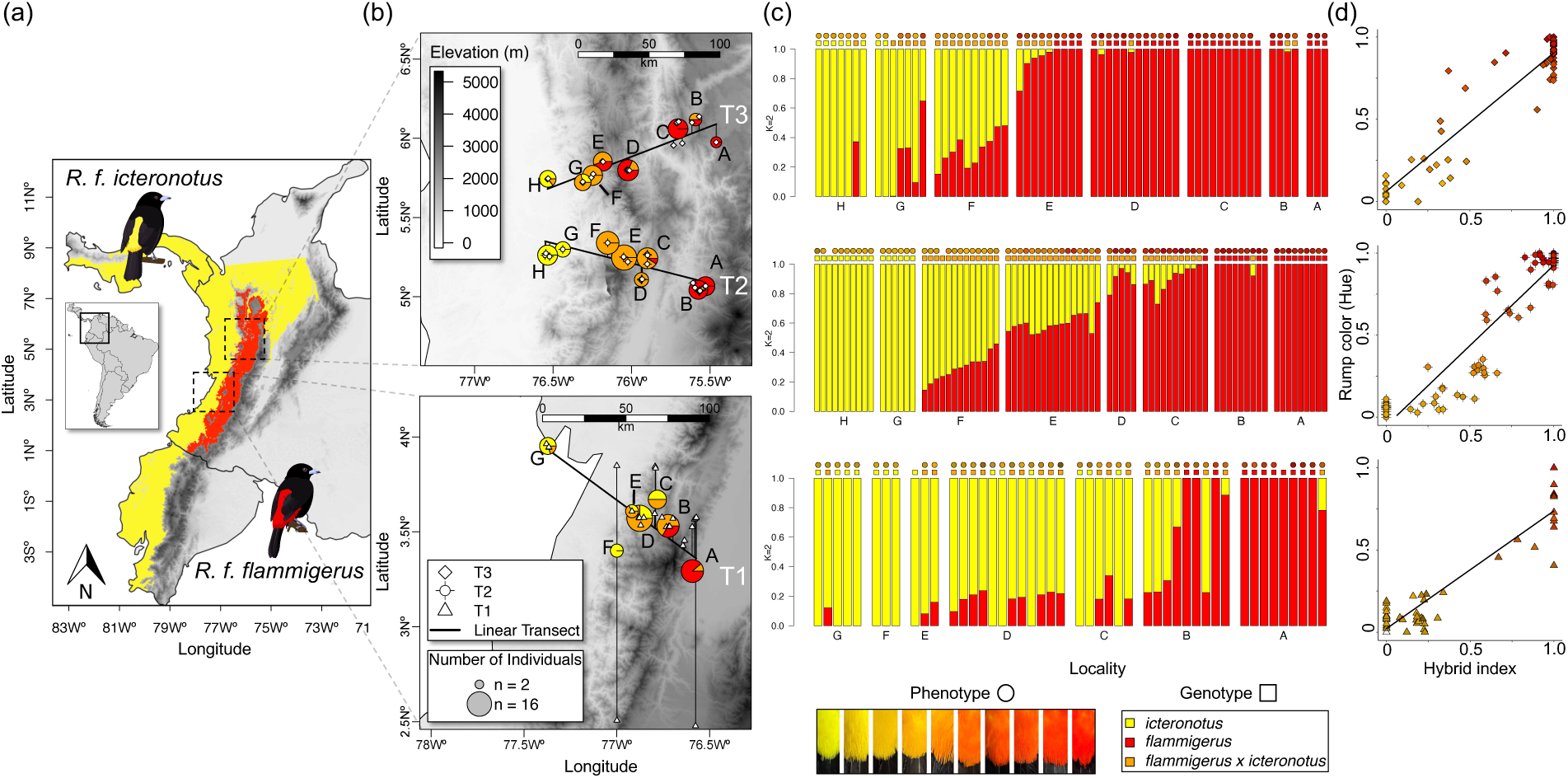
Genetic and phenotypic sampling across replicate transects. A) Geographic ranges of *R. f. flammigerus* (red) and *R. f. icteronotus* (yellow). B) Sampling localities along three geographic transects in the Western Andes of Colombia. Transects coincide with low mountain passes that connect the Chocó biogeographic region in the Pacific to the Cauca Valley. Pie charts show the proportion of pure *R. f. flammigerus* (red), pure *R. f. icteronotus* (yellow) or hybrids (orange) in each locality based on admixture proportions. Circle size represents number of individuals in each locality. C) Admixture proportions (hybrid index – Q) for *k* = 2 from 14,123 SNPs, with individual samples ordered geographically from locality A in the Cauca Valley to locality G (T1) or H (T2 & T3) in the Pacific. Each bar represents a bird, circles above each bar are colored according to the rump color of each bird and squares above each bar represent the genomic ancestry assignment: yellow = pure *R. f. icteronotus* (Q < 0.01), orange = hybrids (0.01 < Q < 0.99), red = pure *R. f. flammigerus* (Q > 0.99). D) Hybrid index is strongly correlated with rump color (hue) in all transects (T1 *r* = 0.93, *p* < 5.44 × 10^−8^; T2 *r* = 0.93, *p* < 1.78 × 10^−35^; T3 *r* = 0.94, *p* < 6.21 × 10^−28^).

The smaller, Lemon-rumped Tanager (*R. f. icteronotus*), hereafter *icteronotus,* has a yellow rump and is found at low through mid-elevations (0–1400m) along the Pacific Coast from Peru to Costa Rica (Hilty, 2020). In Colombia, the Pacific Coast (i.e., Chocó biogeographic region) is mostly dense primary forest, with extremely high precipitation year round (exceeding 250cm annually in some areas) (Urrea et al., 2019). In contrast, the larger Flame-rumped Tanager (*R. f. flammigerus*), hereafter *flammigerus*, has a red rump and is endemic to Colombia, found at mid-elevations along the slopes of the Cauca River Valley (800–2200m; Fig. 1; Burns et al., 2014; Del Hoyo et al., 2018). The Cauca Valley is characterized by monocultures and pastures for cattle ranching with sparse patches of secondary growth, very limited primary forest and a bimodal precipitation regime (mean annual precipitation of 1200–4000 mm at the lowest and highest parts of the mountain respectively; Romero-Hernández et al., 2024). Low mountain passes at multiple locations in the Western Andes serve as bridges where the two subspecies overlap and hybridize extensively, producing replicate hybrid zones with a gradient in coloration and body mass along altitudinal gradients (Bedoya & Murillo, 2012; Morales-Rozo et al., 2017).

We sampled transects along three such low-elevation passes that transverse the Western Andes, and conducted analyses of environmental, phenotypic and genetic variation using reflectance spectrophotometry and genotype-by-sequencing data mapped to a high quality *de novo* genome assembly for *R. f. icteronotus*. We first sought to identify the factors that may influence the likelihood of detecting similar genomic landscapes of differentiation and shared patterns of introgression across the three replicate hybrid zones. Then, we quantified the extent of shared introgression patterns derived from geographic and genomic cline analyses. The replicate hybrid zones provided an opportunity to test for the roles of population structure, ecology, demography and time in producing repeatable outcomes and to observe the progression of introgression through evolutionary time.

## MATERIALS AND METHODS

### Genetic and phenotypic sample collection

We collected morphometric data, blood and feather samples of males and females along three transects in low mountain passes that connect the western and eastern flanks of the Cordillera Occidental in Valle del Cauca (T1), Risaralda (T2), and Antioquia (T3), Colombia (Fig. 1B). We obtained T1 samples (n = 58) from specimens deposited at Museo de Historia Natural de la Universidad de Los Andes (ANDES) collected between 2007–2010, and sampled T2 (n = 80) and T3 (n = 78) in 2022–2023 (see supplementary methods for details). Sample collection and exportation permits in Colombia were issued by the Agencia Nacional de Licencias Ambientales (ANLA) under the Permiso Marco de Recolección No. IDB 0359. Additional *Ramphocelus* tissues were obtained from LSU Museum of Natural Science to span the whole distribution of the species complex (i.e, samples from Ecuador and Panama).

### DNA and RNA sequencing

To produce a high-quality reference genome, a single female *R. f. icteronotus* was sequenced with PacBio HiFi sequencing by the Delaware Biotechnology Institute, yielding 93.9 Gb of data and ∼6.5 million high-fidelity reads. In addition, we extracted RNA from 5 emerging feather follicles of the same female and generated 18 Gb of raw RNAseq data for genome annotation (raw RNA-seq reads are available on Dryad - DOI:10.5061/dryad.zkh1893n9). For population genomic analyses, we extracted genomic DNA from a total of 252 birds (S1 Table) and sent the samples for genotype-by-sequencing (GBS; Elshire et al., 2011) to the University of Wisconsin Biotechnology Center (see supplementary methods for details).

### *De novo* genome assembly and annotation for *R. f. icteronotus*

Methods for reference genome assembly for the female *R. f. icteronotus* are detailed in the supplementary methods. Briefly, we generated a phased assembly with hifiasm v0.16.1 (-primary; Cheng et al., 2021), screened for contaminants with BlobTools2 (Challis et al., 2020) and removed 11 contigs not matching Passeriformes. We assembled the mitochondrial genome with MitoHiFi (16,784 bp, 37 genes) and added it back to the nuclear assembly. We evaluated quality and completeness with BUSCO v5.8 (passeriformes_odb10; Manni et al., 2021) and scaffolded the genome using the chromosome-level assemblies of the Black-throated Flower piercer (*Diglossa brunneiventris*; NCBI assembly ID: GCA_019023105.1) and Zebra finch (*Taeniopygia guttata;* NCBI assembly ID: GCA_003957565.4).

Before annotation, we used RepeatModeler v.2.0.1 (Flynn et al., 2020) to identify and classify repetitive elements *de novo*. We concatenated our *de novo* repeat library with the *T. guttata* repeat database, and used it to soft mask the genome with RepeatMasker v.4.0.7 (Smit et al., 2015). Then, we used STAR v2.7.1 (Dobin et al., 2013) to map the RNA-seq reads to the soft-masked genome. We used the mapped reads (-bam) and the annotated proteins from the published genome of *T. guttata* (NCBI annotation ID: GCF_003957565.2) as evidence input for GeMoMa v.1.9 (Keilwagen et al., 2019), to generate splice site aware homology-based gene predictions.

### Read processing, variant calling and filtering

We processed raw reads from all individuals using the TASSEL GBSv2 pipeline v5.2.65 (Glaubitz et al., 2014) and aligned them to our *R. f. icteronotus* genome with Bowtie2 (Langmead & Salzberg, 2013). Variant calling yielded ∼2.6 million genome-wide SNPs across 254 individuals (TASSEL script is available at https://github.com/mi-castano10/ReplicateHybridZones and the resulting raw unfiltered vcf file is available on Dryad DOI:10.5061/dryad.zkh1893n9). We used VCFtools v0.1.16 (Danecek et al., 2011) to remove samples with mean depth < 1 and retain only high-quality SNPs (biallelic, no indels, site depth ≥ 4 and less than 25% missing data). Then we performed successive rounds of filtering to generate datasets optimized for population genetic analysis with different requirements (Datasets A-D; supplementary methods and Table S3).

### Phenotypic and genomic ancestry assignment

To characterize hybrid zones along a one-dimensional geographic axis, we fitted a linear regression between latitude and longitude, projected sampling sites onto their position along this line and calculated distances from the westernmost point using the R package *sf* (Pebesma et al., 2024). Individuals within ∼5km were assigned to the same locality. Next, to determine whether our sampling transects had parallel patterns of genomic and phenotypic divergence, we assessed the extent to which a given variable predicted genomic ancestry. First, we used 14,123 unlinked SNPs (dataset C; Table S3) as input for ADMIXTURE v.1.3.0 (Alexander et al., 2009) to identify the most likely number of distinct population clusters (k = 1–5) and assign genomic ancestry. We assigned a genomic ancestry score (Q) to each individual, with thresholds of Q < 0.01 as pure yellow *icteronotus*, 0.01 < Q < 0.99 as hybrids and Q > 0.99 as pure red *flammigerus.* Throughout we refer as allopatric ranges to localities with higher average proportions of pure parental individuals (Q < 0.1 & Q > 0.9).

To quantify phenotypic variation, we collected rump reflectance data and ran a principal components analysis (PCA) for morphometric data (see supplementary methods for details). We used hue (wavelength of maximum reflectance; H4) as the main color metric (Montgomerie, 2006) and PC1 as an index for body and bill size. Values for phenotypic traits were scaled between 0 and 1 within each transect. Finally, we calculated the correlation coefficient between genomic ancestry (Q), rump plumage color (hue), body size (PC1) and distance along transect.

### Genomic differentiation and population structure

We tested whether our transects were independent, and the degree of population structure within and between the subspecies ranges. For this we used two complementary approaches based on 14,123 unlinked SNPs (dataset C; Table S3). First, we used EEMS (Estimated Effective Migration Surfaces) to infer and visualize patterns of gene flow and genetic isolation across the geographic landscape (Petkova et al., 2015). We explored nDemes values of 100, 200, and 300 with three randomized replicates each, and since results were consistent across values, we present the intermediate value (nDemes = 200) for optimal resolution. Results were plotted using *rEEMSplots* (Petkova et al., 2015). Second, we conducted a principal components analysis (PCA) with plink v1.9 (Chang et al., 2015), which enabled visualization of genotype clustering and degree of population structure within and between subspecies across transects.

Finally, we used VCFtools v0.1.16 (Danecek et al., 2011) to calculate Weir and Cockerham weighted *F*_ST_ (Weir & Cockerham, 1984) and estimate genome-wide differentiation between pure allopatric populations (excluding all admixed individuals based on ADMIXTURE results). We used 50,868 SNPs (dataset B; Table S3) to calculated pairwise *F*_ST_ between: 1) subspecies using pure red *flammigerus* and yellow *icteronotus* individuals from any available locality (Panama, Ecuador and Colombia), 2) subspecies but for each transect independently, 3) populations of the same subspecies across transects and 4) sampling localities in each transect. Then we used the R packages *StAMPP* (Pembleton et al., 2013) and *ggplot2* to plot the pairwise *F*_ST_ on a heatmap.

### Ecological gradients

To assess whether our sampling spanned similar ecological gradients, we compared environmental variation between localities across transects. We used *geodata* (Hijmans et al., 2024) to obtain bioclimatic data from the 19 WorldClim layers (WorldClim Global Climate Data, 2020) and tree landcover data from the ESA WorldCover data set (Zanaga et al., 2022) at 30-seconds spatial resolution for each GPS coordinate in our sampling transects. Tree landcover represents the fraction (0-100%) of a specified area (e.g., a pixel) covered by tree canopy, which typically reflects the density of trees. In addition, we used *elevatr* (Hollister et al., 2023) to obtain elevation data at zoom = 10 spatial resolution from the Amazon Services Terrain Tiles and the Open Topography global datasets API. We conducted a one-way ANOVA to test for differences in ecological conditions across sampling localities, followed by Tukey’s HSD post-hoc test to identify pairwise differences between groups. Finally, we used *corrplot* (Wei et al., 2024) to test if elevation and tree landcover (%) predicted individual genomic ancestry score (Q) ancestry across transects.

### Demographic history

We reconstructed the demographic history of the three transects using complementary approaches. First, we estimated fluctuations in effective population size (*N*_e_) using PopSizeABC (Boitard et al., 2016), which provides better resolution at more recent time scales, and Stairwayplot2 (Liu & Fu, 2020) for a general view of historical fluctuations.

For PopSizeABC we used 12,103 SNPs (dataset D) from either 5 allopatric (n=10) or 3 sympatric (n=6) individuals per subspecies per transect. We explored *N*_e_ fluctuations in a time window of ∼1.5 Mya to the present (Tmax=550,000 generations in 30 time bins), because dispersal of *R. flammigerus* into South America is estimated at approximately 1.3 Mya (Burns & Racicot, 2009). Following scripts from Bemmels et al. (2021), for each population we calculated observed summary statistics in 2 Mbp windows and simulated 450,000 datasets of 100 2 Mbp segments, using log-uniform *N*_e_ priors (10-100,000) and a limit of 10-fold change between adjacent bins. We used a recombination rate of 3.1 × 10^−8^, reported for *Ficedula* flycatchers (Kawakami et al., 2014), and a per generation mutation rate of 4.47 × 10^−9^ derived from recalibrating the yearly mutation rate from *Geospiza fortis* (Nadachowska-Brzyska et al., 2015), with an estimated generation time of 2.6 years for *R. flammigerus* (Bird et al., 2020). ABC inference was conducted separately for each transect and population type, retaining the top 0.1% of best-fitting simulations to generate posterior *N*_e_ distributions (time reported in years and *N*_e_ in number of haploid individuals).

Next, we used 5006 SNPs (a thinned version of dataset D) as input for easySFS (Gutenkunst et al., 2009; Overcast, 2016/2025) to construct a folded site-frequency spectrum for 6 pure individuals from allopatric populations of each subspecies and 6 hybrid individuals from sympatric populations in each transect. We calculated the total number of observed nucleic sites (including polymorphic and monomorphic) and used the SFS for allopatric populations to run Stairwayplot2 (Liu & Fu, 2020) with nseq=12 (haploid sample size), smallest size of SFS bin = 1, largest size of SFS bin = 6, ninput=200 and 2,5,7,10 as number of random break points for each try.

For each transect, we tested 15 demographic models in fastSIMCOAL2 (v27.09; Excoffier et al., 2021) using the previously calculated SFS from three population groups (AB - allopatric red *flammigerus*, CDEF - hybrids, GH allopatric yellow *icteronotus*), including null models (1–3), allopatric divergence (4–6, 13–15), and divergence with geneflow (7–9, 10–12), with or without population expansion (S9 Fig). In allopatric models Tm1 marks subspecies divergence, while in divergence with geneflow models Tm1 indicates hybrid zone formation and Tm2 reflects an increase in gene flow. For all models we used the same mutation rates and generation times as for PopSizeABC, and parameter estimates for current *N*_e_ were bounded to the posterior distributions estimated with this method (1 – 100,000). We ran each model 100 times, with 40 EMC loops and 250,000 simulated SFS per iteration, selected the run with the highest likelihood for each model and defined the best fit model using Akaike’s information criterion (AIC). Finally, we calculated confidence intervals via bootstrap (Table 1; see supplementary methods for details). Example input files for models 4-9 are available in (https://github.com/mi-castano10/ReplicateHybridZones).

**Table 1.** Parameter estimates of the best fit demographic model inferred with FastSIMCOAL2 for each transect.

| Parameter | T1 - Model 7 | T2 - Model 8 | T3 - Model 8 |
| --- | --- | --- | --- |
| Ne ICT | 45131 (42805-45635) | 40710 (38758-40866) | 33990 (32014-37897) |
| Ne HYB | 49315 (49058-49283) | 49358 (49106-49485) | 49483 (49023-49593) |
| Ne FLA | 8491.5 (8417-8644) | 8919 (8790-9027) | 6851.5 (6815-7082) |
| **TIME 1** | **6123 (6075-6198)** | **5044 (5030-5268)** | **3341 (3289-3366)** |
| ANCSIZE | 493216.5 (493196-493403) | 499427.5 (499128-499528) | 494837 (494815-494835) |
| Nm HYB → ICT | 0.039 (0.0384-0.0414) | 0.031 (0.0307-0.0341) | 0.0347 (0.0347-0.0423) |
| Nm ICT → HYB | 24.672 (22.5967-24.8132) | 18.9588 (18.4709-22.954) | 18.9468 (17.4755-19.6061) |
| Nm FLA → HYB | 0.0338 (0.0332-0.0382) | 0.0504 (0.047-0.062) | 0.033 (0.0324-0.0353) |
| Nm HYB → FLA | 0.0264 (0.0263-0.0265) | 0.0279 (0.0279-0.0279) | 0.0273 (0.0273-0.0274) |
| Mig. rate HYB → ICT | 0.0000004 (0.0000004-0.0000005) | 0.0000004 (0.0000004-0.0000004) | 0.0000004 (0.0000004-0.0000005) |
| Mig. rate ICT → HYB | 0.0002734 (0.0002578-0.0002813) | 0.0002329 (0.0002248-0.0002932) | 0.0002788 (0.0002378-0.000286) |
| Mig. rate FLA → HYB | 0.000002 (0.000002-0.0000023) | 0.0000029 (0.0000027-0.0000035) | 0.0000025 (0.0000024-0.0000026) |
| Mig. rate HYB → FLA | 0.0000003 (0.0000003-0.0000003) | 0.0000003 (0.0000003-0.0000003) | 0.0000003 (0.0000003-0.0000003) |
| **TIME 2** |  | 2498.6 (1898-2534) | 1112.8 (1081-1183) |
| Recent Nm HYB → ICT |  | 0.3926 (0.3455-2.1985) | 0.9998 (0.8809-1.4738) |
| Recent Nm ICT → HYB |  | 41.5471 (40.4412-42.4435) | 35.6937 (32.8995-35.7186) |
| Recent Nm FLA → HYB |  | 0.4255 (0.4038-0.6601) | 37.6062 (36.1468-37.9432) |
| Recent Nm HYB → FLA |  | 0.0833 (0.0815-0.5224) | 0.2736 (0.2021-2.4603) |
| Recent Mig. rate HYB → ICT |  | 0.000004 (0.0000036-0.0000224) | 0.0000102 (0.000009-0.000015) |
| Recent Mig. rate ICT → HYB |  | 0.0005103 (0.0005063-0.0005411) | 0.0005251 (0.0004453-0.0005893) |
| Recent Mig. rate FLA → HYB |  | 0.0000239 (0.0000228-0.0000367) | 0.0027443 (0.0025843-0.0027576) |
| Recent Mig. rate HYB → FLA |  | 0.0000009 (0.0000009-0.0000054) | 0.0000028 (0.0000021-0.0000251) |

Age of formation of a stable hybrid zone (Time 1 in bold) and time of increase in gene flow (Time 2) are in years and effective population size
(Ne) is in number of diploid individuals. Nm indicates number of migrants, ICT = *icteronotus*, FLA = *flammigerus* and HYB = hybrids. Numbers
in parenthesis are confidence intervals.

### Admixture surface and environmental transitions

To test whether the hybrid zones are constrained to an ecotone or if they have shifted through time, we estimated the location of the genetic center of each transect and assessed its position relative to the most relevant ecological transitions between the ranges of both subspecies. First we used the coordinates and genomic ancestry scores (Q) to predict a spatial admixture surface with the *krig* function from the R package *fields* (Nychka et al., 2024). Then we used the *surface* function to estimate the contour line (hereafter isocline) at which the genomic ancestry score (Q) = 0.5 across the geographic landscape, which was set as the genetic center of the hybrid zone. To obtain comparable distances, we centered all transects at the point where they intersected the Q = 0.5 isocline and calculated the distance of each site along the linear transect relative to this interception point (*i.e.*, the genetic center; Fig 4). To estimate the locations of relevant environmental transitions (elevation and tree landcover %), we calculated the center of the range of overlap in these two variables between allopatric subspecies. We extracted elevation and tree landcover data for 158 georeferenced localities for the pure red *flammigerus* and 385 for the pure yellow *icteronotus* subspecies from museum specimens (GBIF, 2025a, 2025b). Locations outside the 95% confidence interval for each subspecies were filtered out to remove outlier observations. Then, we calculated the center of the overlap between the elevational and tree landcover ranges (Fig S7A). Lastly, we used the raster layers to plot the elevation and tree landcover contour lines that corresponded to the center of the overlap using the *rasterToContour* function of the R package *raster* (Hijmans et al., 2025). We defined the contour lines as our environmental isoclines and used the points at which the linear transects intersect the environmental isoclines as the location at which environmental transitions occur in each transect (Fig 4A; S7B Fig; S8-10 Tables).

### Classification of hybrid classes

To classify F1s, later-generation hybrids, backcrosses, and parental individuals in each transect we used triangle plots of interspecific heterozygosity vs. hybrid index constructed with *Introgress* (Gompert & Alex Buerkle, 2010). We used our estimations of overall Weir and Cockerham’s *F*_ST_ to identify informative markers. Because only two SNPs were fixed between subspecies, we used 15 highly differentiated SNPs (*F*_ST_ > 0.9; S4 Fig) as semi-diagnostic markers to perform the hybrid index and heterozygosity calculations. However, constructing triangle plots with SNPs that are not fixed can result in F1 hybrids with lower heterozygosity values than expected. Thus, we used a relaxed threshold for heterozygosity (> 0.8) to identify potential F1 hybrids (as in Ocampo et al., 2023; Scordato et al., 2017).

### Geographic clines and neutral diffusion

We used *HZAR* (Derryberry et al., 2014) to fit genomic and phenotypic data to geographic cline models to estimate the extent of introgression between yellow *icteronotus* and red *flammigerus*, and gauge the position of clines relative to the elevation and tree landcover environmental transitions. Because *R. flammigerus ssp.* are sexually dimorphic, we fitted geographic clines for males and females separately.

Also, to evaluate how clines for specific diagnostic markers compare to the overall genomic cline, we used VCFtools (Danecek et al., 2011) to estimate the allele frequencies for the two fixed SNPs between subspecies (Table 2; S11 Table). To obtain comparable displacement data, we used distances from the one-dimensional geographic axis, where linear transects are centered at the Q = 0.5 isocline (S8-10 Tables). For genomic data (Q and two diagnostic SNPs) we evaluated 16 models, and for phenotypic data we evaluated 6 models (S12 Table). We ran the MCMC model-fitting method in *HZAR* with three fit requests for each model, default chain length (1 × 10^6^), burn in (1 × 10^4^) and thinning (100), but made each fit request for each model run off a separate seed (Derryberry et al., 2014). We selected the best fit model using AIC and estimated cline centers and widths with their respective 95% confidence intervals using the *hzar.getLLCutParam* function. We determined whether any two given clines were coincident (overlap in their center) or concordant (overlap in their width) by visual observation of nonoverlapping confidence intervals.

**Table 2.** Geographic cline analysis.

| Transect | Sex | Trait | Best model | Center (95% CI) | Width (95% CI) |
| --- | --- | --- | --- | --- | --- |
| 1 | Male | Rump color (Hue) | Model 4 | 2 (0.62-3.56) | 5.56 (3.04-8.36) |
|  |  | Q | Model 6 | -0.53 (-10.08-4.99) | 33.55 (12.65-37.66) |
|  |  | SNP 1 | Model 1 | -3.5 (-9.83-5.3) | 30.4 (16.98-64.9) |
|  |  | SNP 2 | Model 1 | 1.94 (-1.82-2.98) | 1.23 (0.03-14.2) |
|  | Female | Rump color (Hue) | Model 1 | 11.13 (5.25-19.05) | 12.02 (10.84-80.54) |
|  |  | Q | Model 1 | 0.64 (-10.45-12.69) | 31.73 (14.86-87.99) |
|  |  | SNP 1 | Model 1 | -4.52 (-13.82-5.64) | 38.2 (21.69-83.02) |
|  |  | SNP 2 | Model 1 | -5.21 (-13.74-3.8) | 29.92 (16.23-66) |
| 2 | Male | Rump color (Hue) | Model 1 | 6.75 (5.05-8.77) | 18.06 (12.68-22.94) |
|  |  | Q | Model 6 | 2.52 (-7.83-10.11) | 33.73 (17.08-76.47) |
|  |  | SNP 1 | Model 1 | 2.35 (-6.99-8.91) | 44.84 (26.81-83.86) |
|  |  | SNP 2 | Model 1 | 6.14 (0.55-11.32) | 32.08 (19.99-57.28) |
|  | Female | Rump color (Hue) | Model 1 | 14.58 (11.2-17.18) | 21.55 (14.61-29.16) |
|  |  | Q | Model 1 | 3.45 (-10.53-15.89) | 37.04 (16.81-88.82) |
|  |  | SNP 1 | Model 1 | 2.12 (-6.13-8.93) | 27.53 (14.88-57.45) |
|  |  | SNP 2 | Model 1 | 14.22 (6-24.06) | 32.9 (17.58-68.99) |
| 3 | Male | Rump color (Hue) | Model 1 | -2.42 (-5.1-0.05) | 17.07 (7.28-24.36) |
|  |  | Q | Model 6 | -1.36 (-9.02-5.12) | 15.61 (6.32-47.5) |
|  |  | SNP 1 | Model 1 | -10.5 (-31.52-2.51) | 60.52 (31.7-87.63) |
|  |  | SNP 2 | Model 2 | -6.24 (-8.58-3.24) | 5.96 (1.69-43.6) |
|  | Female | Rump color (Hue) | Model 3 | 4.73 (0.25-5.65) | 1.93 (0-23.36) |
|  |  | Q | Model 6 | 0.57 (-5.85-8.05) | 8.28 (0.11-53.23) |
|  |  | SNP 1 | Model 1 | -24.05 (-87-1.84) | 87.47 (50.69-100) |
|  |  | SNP 2 | Model 1 | -4.5 (-22.53-9.66) | 40.36 (17.29-103.3) |

Best-supported model and estimates for cline center (displacement in km from the Q isocline), width (km)
of genetic and phenotypic clines.

To infer whether genomic or phenotypic traits are involved in reproductive isolation, we compared our cline width estimations with the expected width of geographic clines under neutral diffusion (formula in Barton & Gale, 1993). We assumed a generation time of 2.6 years (Bird et al., 2020), dispersal distances (σ) of 1-20 km (Morales-Rozo et al., 2017) and explored range of time of onset of hybridization (t) from 1000 – 6000 years equivalent to 384 – 1923 generations based on the estimated age of the hybrid zones obtained from demographic modeling.

### Genomic Clines

In contrast to geographic clines, genomic clines describe patterns of allele frequency change for individual markers as a function of the hybrid index (Fitzpatrick, 2013; Gompert & Buerkle, 2011). To ameliorate the effects of uneven sampling across transects, we used the Bayesian MCMC approach implemented in *gghybrid* (Bailey, 2024) to estimate genomic clines and identify any shared patterns in locus-specific introgression. We used PGDspider (Lischer & Excoffier, 2012) and custom R scripts to convert a vcf with 50,868 SNPs (dataset B; Table S3) for each transect into the *Structure* input file format required by *gghybrid*. Parental reference individuals were defined as those with Q > 0.99 or Q < 0.01 from admixture. Then we estimated genomic clines for all loci that had an allele frequency difference of > 0.5 between pure parental individuals in each transect. Loci were further filtered so that they would have a minor parental allele frequency < 0.1 and no overlap between parental populations in the Bayesian posterior 95% credible intervals of allele frequency (AF.CIoverlap = FALSE). We ran the MCMC chain for 7000 iterations with a 3000 iteration burnin for both hybrid index and genomic cline estimations. To confirm significance of outlier loci, we ran the genomic cline estimation two more times but with parameters fixed to null values (center = 0.5 or steepness = 1). We then compared the AIC values from the different runs. Loci with a negative AIC difference were deemed significant, indicating that a steeper-than-expected or displaced cline provided a better fit than the null model (Bailey, 2024). Last, we assessed the degree of outlier sharing between transects by calculating the number of shared 100kb windows along the genome with at least one outlier locus (S13-15 Tables).

To estimate whether clines for outlier loci were coincident with plumage color, we fitted a sigmoidal cline for rump hue as a function of hybrid index. Following the methodology in Morales-Rozo et al. (2017), we used Hill functions of the *drc* software to fit a log-logistic two-parameter model, with hybrid index as the dose and rump color as the response. The non-linear function (LL.2) has two parameters: 1) slope, which determines the steepness and 2) the effective dose at which 50 % of the response is achieved (center). Then, we fit 1000 clines in a bootstrap with replacement to obtain confidence intervals (S13 Fig).

### Genome Wide Association Study (GWAS)

We performed a genome-wide association study (GWAS) with individuals from all transects combined to identify regions of the genome associated with rump hue (H4). First, we used BEAGLE v5.5 (Browning et al., 2018) to impute missing data from dataset B (50,868 SNPs). Then we used GEMMA v0.98.5 (Zhou & Stephens, 2012) to fit Bayesian sparse linear mixed models to the data accounting for population structure and relatedness. We implemented the Wald test to evaluate association significance and applied a Bonferroni correction for multiple comparisons to calculate the significance threshold (alpha < 1.115 × 10^-6^).

## RESULTS

### Chromosome scale reference genome for *R. f. icteronotus*

We generated the first near chromosome-scale *de novo* genome assembly for *Ramphocelus.* The genome was 1.3 Gb long, contained in 109 scaffolds and 336 contigs. The scaffold and contig N50 were 75.4 and 71.1 Mb respectively, with > 95% of all the sequence contained in the largest 42 scaffolds, corresponding to 39 autosomes, the Z and W chromosomes and the mitochondria (S2Table; S2 Fig). Our genome was very complete, with 99% of Passeriformes BUSCOs present in single copy (C:99.4%[S:99.0%,D:0.4%],F:0.4%,M:0.3%,n:10844). Approximately 14.6% of the genome was contained in repeats and the homology-based annotation recovered a total of 17,128 genes (The final assembly and GFF file for the annotation are available on DOI: 10.5061/dryad.zkh1893n9).

### Parallel patterns of phenotypic and genomic divergence

Whether analyzing each transect independently or combining data for all transects, ADMIXTURE consistently showed highest support for *k*=2 as the optimal number of genetic clusters (S4 Table). Across transects, individuals sampled in locality A had a high probability of belonging to the red *flammigerus* subspecies (Q > 0.97, T1 mean = 0.98 ± SD 0.07, T2 mean = 0.99 ± 0.0, T3 mean = 0.99 ± 0.0) and individuals sampled in localities G (T1) or H (T2 & T3) had a high probability of belonging to the yellow *icteronotus* subspecies (Q < 0.05, T1 mean = 0.02 ± SD 0.05, T2 mean = 0.0001 ± 0.0, T3 mean = 0.05 ± 0.14). Additionally, distance along the transect was a strong predictor of genomic ancestry score in all transects (T1 *r* = 0.71, *p* < 3.48 × 10^−8^; T2 *r* = 0.93, *p* < 7.37 × 10^−35^; T3 *r* = 0.76, *p* < 1.78 × 10^−12^; S3 Fig). However, the proportion of genetically admixed individuals (0.01 < Q < 0.99) and the localities in which they occurred, differed among transects. In T1, 46% of individuals were genetically admixed and occurred at both the westernmost (G) and easternmost (A) localities, which were previously considered pure allopatric ranges (Fig 1C). Also, locality B of T1 was the only locality in which pure parental individuals with Q > 0.9 and Q < 0.1 occurred in sympatry. In T2, 56% of individuals were genetically admixed but none of them occurred in localities at the extremes of the transect in the allopatric ranges. In T3, 46% of individuals were genetically admixed and occurred at both the westernmost locality (H) and the easternmost locality (A). From field observations and preliminary admixture analysis from 9 admixed individuals, we detected a fourth hybrid zone spanning from localities A and B of T3 towards the northeast. This fourth hybrid zone is likely to be recent, as the yellow *icteronotus* subspecies is expanding its range into the Magdalena River valley. Because localities A & B were previously considered to be part of the allopatric range of the red *flammigerus* subspecies in T3, we removed the admixed individuals in these localities from all our analysis, reassigned pure individuals in locality A and B and re-ran the admixture analysis accordingly to keep the sampling consistent among replicates.

Rump color (hue) was strongly correlated with genomic ancestry in all transects (T1 *r* = 0.93, *p* < 5.44 × 10^−8^; T2 *r* = 0.93, *p* < 1.78 × 10^−35^; T3 *r* = 0.94, *p* < 6.21 × 10^−28^; Fig 1D). However, contrary to previous findings suggesting that subspecies exhibit variation in body size and that this trait varies gradually along the hybrid zone in T1 (Bedoya & Murillo, 2012; Morales-Rozo et al., 2017), our proxy for body size (morphological PC1) did not show subspecific variation and was not correlated with genomic ancestry in T2 and T3 (T1 *r* = 0.62, *p* < 4.20 × 10^−6^; T2 *r* = 0.2, *p* < 0.08; T3 *r* = −0.08, *p* < 0.56; S3 Fig). Thus, we excluded body size from any further analysis. Overall, our results indicated parallel patterns of phenotypic and genomic divergence across transects, though the presence of admixed individuals in more eastern or western localities in a given transect suggested varying extents of introgression.

### Independent but replicated transects

The estimated effective migration surface (EEMS) showed that high-elevation peaks in the Western Andes (i.e., Cerro Tatamá and Farallones del Citará) act as strong barriers to gene flow between hybrid zones, suggesting that transects are independent (Fig. 2A). In contrast, populations on either side of the mountain exhibited higher connectivity, leading to increased genetic similarity and little population structure within subspecies. The only exception was the allopatric population of the yellow *icteronotus* subspecies in T1, which was separated from T2 and T3 by an area with reduced gene flow.

**Figure 2.**
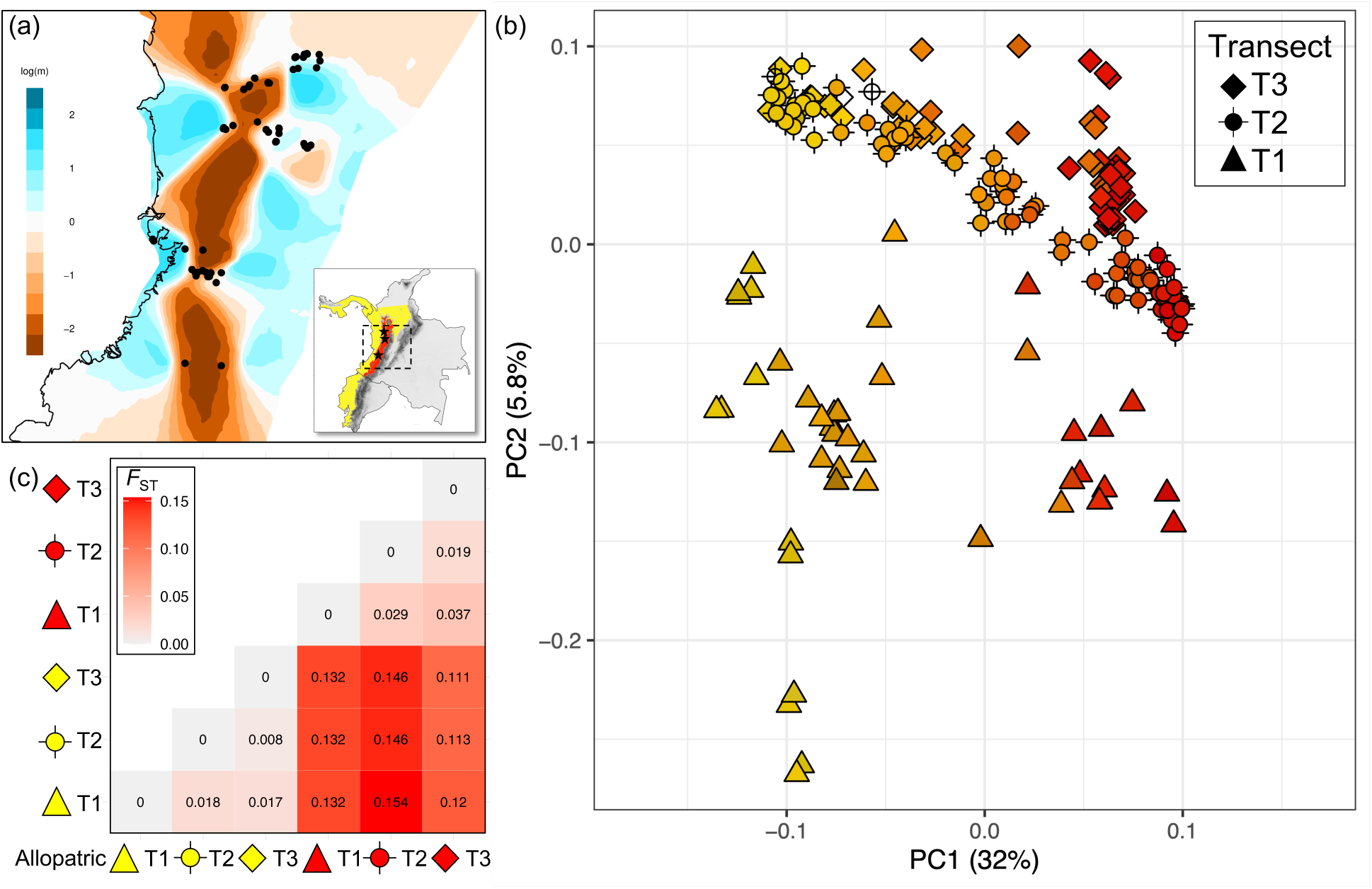
Genomic differentiation and population structure across the landscape indicate replicated and independent hybrid zones. A) Effective migration surface analysis revealed independent transects as gene-flow is reduced (orange) between hybrid zones along the mountain but is unimpeded or increases (blue) between parental populations across their entire geographic range (nDemes =200). B) Genetic PC1 and PC2 of 14,123 genome-wide SNPs with points colored according to the rump color phenotype of each bird. C) Heatmap of pairwise *F*_ST_ between pure allopatric populations of each subspecies across transects suggest low population structure within subspecies (light colors) and higher genomic differentiation between subspecies (warm colors).

Genetic PC1 explained 32% of the variation in the data and separated the two subspecies into clear clusters, with intermediate values for hybrid individuals (Fig 2B). Because the genetic PC1 and the genomic ancestry score (Q) for k = 2 were highly correlated (T1 *r* = 0.97, *p* < 4.88 × 10^−31^; T2 *r* = 0.99, *p* < 2.37 × 10^−91^; T3 *r* = 0.99, *p* < 1.04 × 10^−55^), they readily distinguished parental subspecies and hybrids, and were strong predictors of rump color (Fig 1D, 2B). Subsequent components explained much less of the variation in the data (PC2 = 5.8%, PC3 = 4.6%, PC4 = 4.3%). However, PC2 separated the samples geographically, closely matching the location and distribution of the transects. PC2 showed low within-subspecies population structure and mirrored the results found with the EEMS analyses. Finally, genome-wide differentiation between subspecies was limited, with *F*_ST_ = 0.14 and only 2 fixed SNPs in chromosomes 6 and Z (S4 Fig). Differentiation between subspecies calculated with individuals only at the pure parental ends of each transect was similar, albeit slightly higher in T2 than in the other transects (T1 *F*_ST_ = 0.13, T2 *F*_ST_ = 0.15, T3 *F*_ST_ = 0.11; Fig 3C). In contrast, genomic differentiation within subspecies across transects was consistently low (*F*_ST_ < 0.04) and reflected their geographic distribution, with T1 and T3 being farther apart and exhibiting greater differentiation (Fig 3C). Mean pairwise *F*_ST_ estimates between sampling localities also revealed low overall levels of divergence (*F*_ST_ T1 = 0.05, *F*_ST_ T2 = 0.06, *F*_ST_ T3 = 0.05) and nearly undetectable differentiation between neighboring localities across transects (*F*_ST_ T1 = 0.02, *F*_ST_ T2 = 0.01, *F*_ST_ T3 = 0.02; S5 Fig).

**Figure 3.**
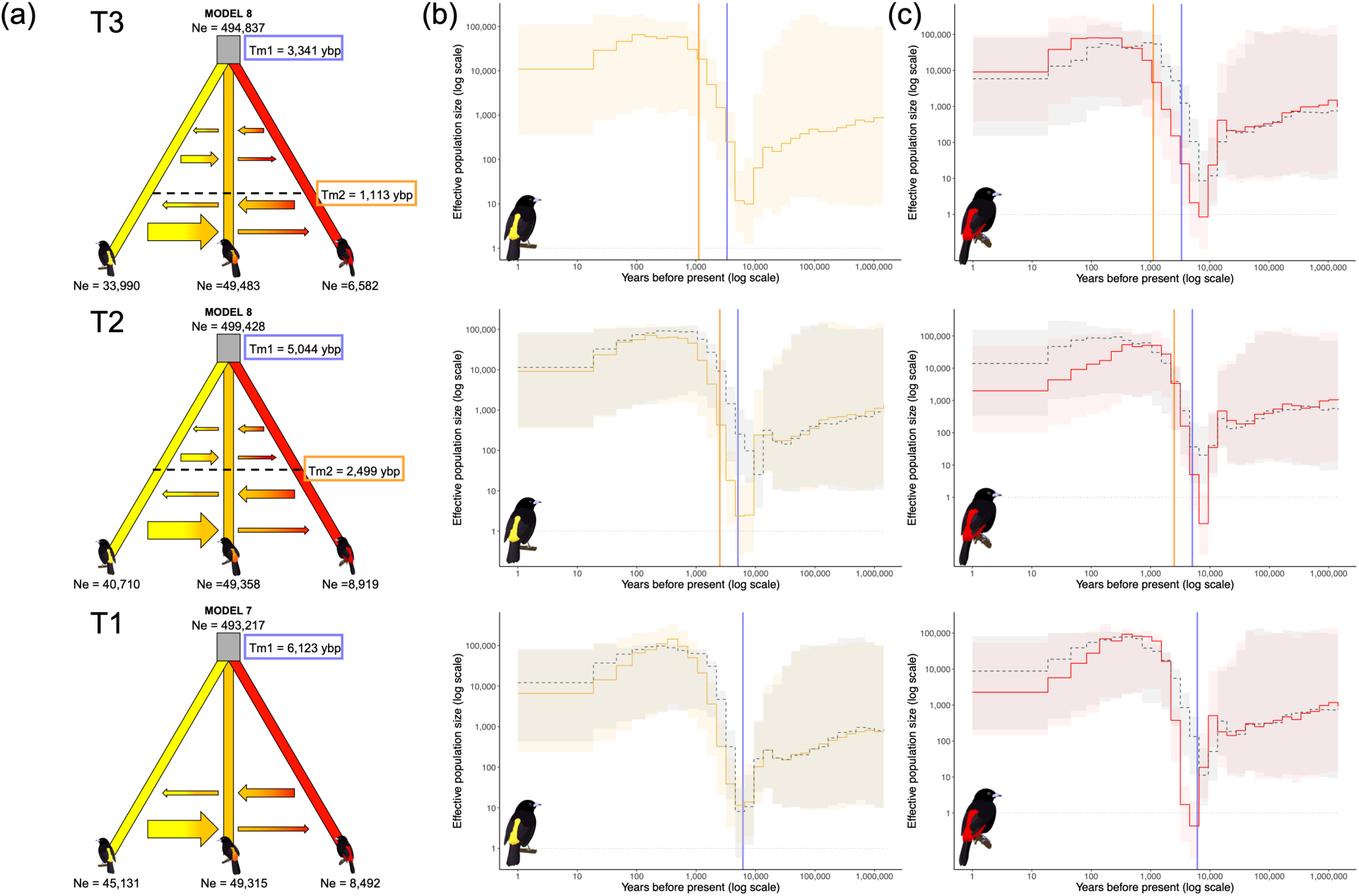
Demographic modeling revealed populations have similar demographic histories, but hybrid zones have different ages. **A)** Across transects a model of divergence with geneflow was the best supported (T1 – bottom = model 7, T2 – middle and T3 – Top = model 8). Red lines represent pure individuals from the allopatric ranges of the *flammigerus* subspecies (localities A, B). Yellow lines represent pure individuals from the allopatric ranges of the *icteronotus* subspecies (localities H, F). Orange lines represent admixed individuals from the hybrid zone (localities C, D, E, F). Grey lines on top of each model represent the ancestral population. In all models time flows from the top to the bottom and arrows represent migration. The size of the arrows is proportional to the amount of gene-flow in each time period. In all models Time 1 (Tm1 - blue) represents the time of the hybrid zone formation and Time 2 (Tm2 - orange) represents the time of increase in gene flow. Parameter estimates for effective population sizes are in diploid number of individuals, and time is in years (assuming 2.6 years per generation). **B & C)** Effective population sizes (Ne) of pure allopatric (solid colored line) and sympatric (dashed grey line) populations of each subspecies estimated with PopSize ABC. Lines represent the time of onset of hybridization (blue) and time of increase in gene flow (orange) estimated with coalescent modeling. Shading represents the 95% confidence interval of the median.

### Parallel ecological gradients

We found that 15 out of the 19 WorldClim bioclimatic variables related to temperature and precipitation were strongly correlated with elevation in this area of Colombia (S6 Fig). Thus, we used elevation as a proxy for climatic conditions. We also used tree landcover (%) as descriptors of ecological conditions because deforestation seems to be facilitating hybridization (Sibley, 1958). All transects traversed gradients from humid, warm, low elevations and dense vegetation in the Pacific to drier, colder, higher elevations and more disturbed ecosystems in the Cauca valley. Thus, steep environmental and altitudinal gradients were replicated, albeit with some variability across localities (S7 Fig).

Elevation was positively correlated with genomic ancestry in all transects (T1 *r* = 0.78, *p* < 2.3 × 10^−10^, T2 *r* = 0.94, *p* < 1.2 × 10^−35^, T3 *r* = 0.9, *p* < 4.01 × 10^−22^; S3 Fig). Birds at higher elevations and eastern localities along all transects tended to have higher proportions of red *flammigerus* ancestry, and birds at lower elevations and western localities tended to have higher proportions of yellow *icteronotus* ancestry. In contrast, tree landcover was negatively correlated with hybrid index in all transects (T1 *r* = - 0.52 *p* < 0.02 × 10^−2^, T2 *r* = −0.61 *p* < 2.84 × 10^−9^, T3 *r* = −0.59 *p* < 5.84 × 10^−7^; S3 Fig). Birds at western localities in the Pacific rainforest, which has greater tree landcover, had higher proportions of yellow *icteronotus* ancestry, while birds at eastern localities, in the Cauca valley where deforestation is extensive, had higher proportions of red *flammigerus* ancestry.

### Similar demographic history but different time of hybrid zone formation

Our reconstruction of fluctuations in effective population size (*N*_e_) in recent time with PopSizeABC (Boitard et al., 2016) showed that all populations experienced a significant crash in effective population size around 6000 - 10000 years before present (ybp), followed by a rapid population expansion towards the present (Fig 3B). Broader estimations based on the site-frequency spectrum using Stairwayplot2 (Liu & Fu, 2020) also showed a recent crash in effective population size, but revealed a population expansion that happened only in the yellow *icteronotus* subspecies around 250,000 ybp (S8 Fig). The overall trends were similar across populations of the same subspecies and both methods converged in relatively low current effective population sizes (Fig 3B & S8 Fig).

Across transects, the best-fit coalescent model was consistently a model of divergence with gene flow, where subspecies likely diverged in isolation with sporadic gene flow during the Pleistocene. However, the hybrid zones became well-established (*i.e.,* sustained gene flow) more recently in the slopes of the Western Andes (Fig 3A). In T1, coalescent model 7 was the best fit (S5 Table), which indicates that gene flow has remained constant from the moment of split. In T2 and T3, model 8 was the best fit (S6 & S7 Tables), which also suggests there was gene flow from the moment of population split, but rather than remaining constant, gene flow increased at a certain point in time. Parameter estimates for effective population sizes, magnitudes and directionality of gene flow were similar across transects (Table 1). Effective population size was always higher in the yellow *icteronotus* subspecies than in the red *flammigerus* subspecies. Also, gene flow was > 15-fold higher from the yellow *icteronotus* subspecies into the hybrid zone in all transects. However, the time at which the hybrid zones were formed differs. Sustained hybridization began first in T1, at least 6,000 ybp, then in T2 around 5,000 ybp and most recently in T3 around 3,000 ybp. Estimations for the age of the hybrid zones were consistent with a northeast population expansion of the yellow *icteronotus* subspecies starting from T1, which then resulted in a later pulse of admixture in T2 and T3. Importantly, the time of onset of hybridization estimated with fastSIMCOAL2 was coincident with the time of rapid population expansion after a population crash estimated with PopSizeABC.

### Progressive hybrid zone movement relative to environmental transitions

We defined the environmental transitions as the center of the overlap between the elevation and tree landcover ranges of allopatric subspecies (see methods; S7A Fig). The yellow *icteronotus* subspecies is found from 0 – 1400 m elevation and at 70% to 100% tree landcover. The red *flammigerus* subspecies is found from 800 – 2100 m elevation and at 30% to 99% tree landcover. The elevation overlap between subspecies occurred between 800 and 1400m, and the center of the area of overlap (hereafter elevation transition) occurred at 1100m. The tree landcover overlap between subspecies occurred between 70% and 99%, and the center of the area of overlap (hereafter landcover transition) occurred at 85% tree landcover (Fig S7A). Then, we calculated the intersection point between the linear transects and the 1100m and 85% canopy cover isoclines and used an admixture surface to estimate where the genetic center of the hybrid zone occurred relative to these intersection points representing environmental transitions (Fig 4A; S7B Fig). We found that the distance from the environmental transitions to the genetic center differed across transects and was consistent with the age of the hybrid zone estimated from demographic modeling (Fig 4B). In T1—the oldest hybrid zone—the genetic center was displaced 11.8 km east from the elevation transition and only 3.4 km west from the tree land cover transition. In T2—the intermediate hybrid zone—the genetic center was also displaced 12km east from the elevation transition but 25 km west of the tree landcover transition. In T3—the youngest hybrid zone—the genetic center occurred only 1.4 km east from the elevation transition and 33.5 km west from the tree landcover transition (Fig 4B).

**Figure 4.**
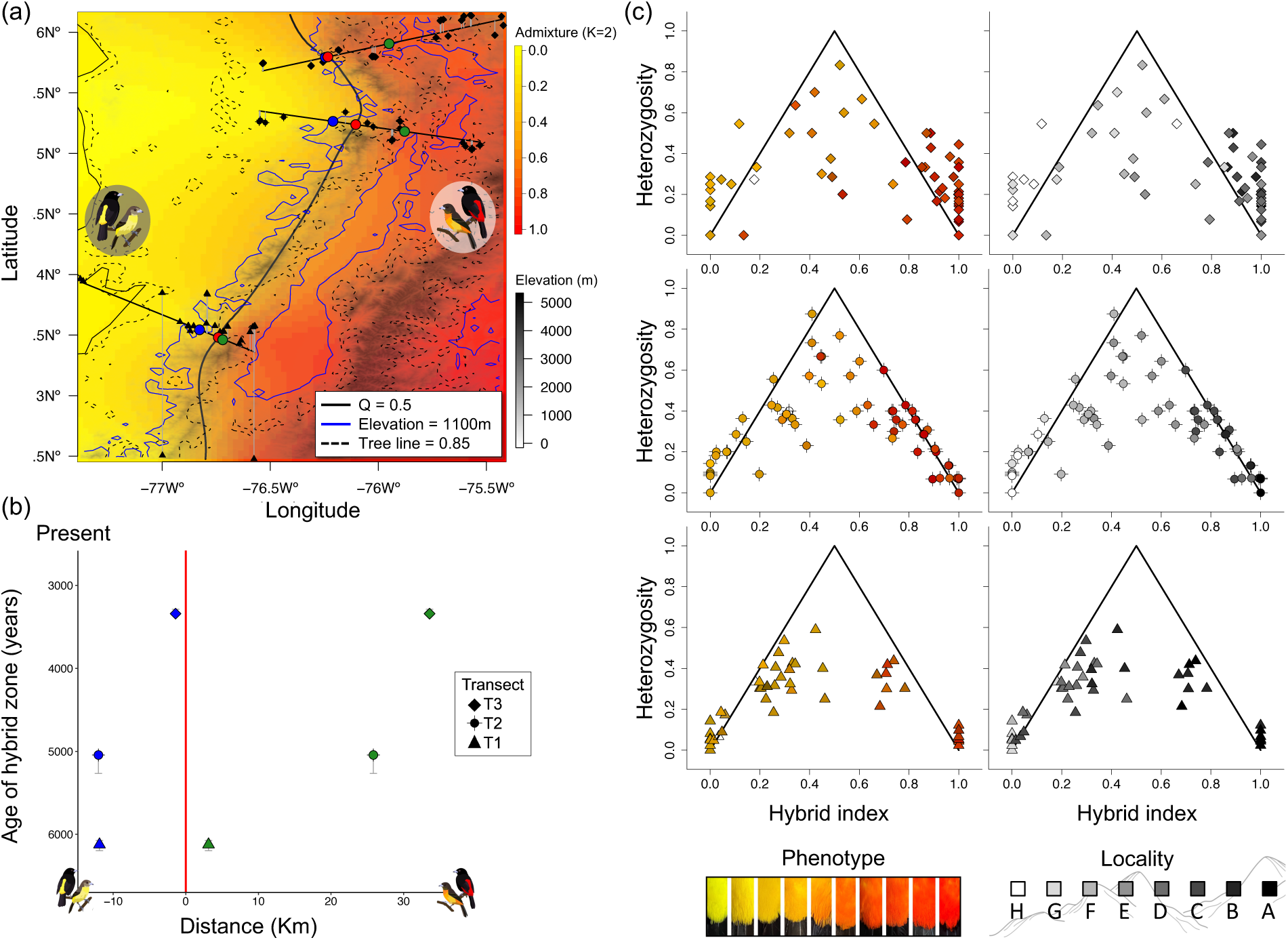
Extent of hybrid zone displacement from environmental transitions and characterization of hybrid classes are consistent with the age of the hybrid zones. **A)** Admixture surface prediction across the geographic landscape. Lines represent the geographic isoclines where the hybrid index = 0.5 (solid-black), elevation is 1100 m (solid-blue) and tree land cover is 85% (dashed-black). Colored points along the linear transects indicate the interception point with the isoclines, representing the genetic center (red) the elevation transition (blue) and the tree landcover transition (green) in each transect. **B)** Distance from environmental transitions show progressive displacement of the hybrid zones through time. The genetic center (red line) is displaced from the elevation transition (blue) in the oldest (T1) and intermediate (T2) hybrid zones but not in the youngest (T3) and is closer to the tree land cover transition (green) in T1 and farther away in T2 and T3. Vertical lines represent the 95% confidence intervals for time of onset of hybridization estimated with FastSIMCOAL2. **C)** Triangle plots of hybrid index vs. interspecific heterozygosity calculated using 15 diagnostic SNPs between subspecies (*F*_ST_ > 0.9). Left panel is color coded according to the rump color phenotype of each bird and right panel is color coded according to sampling locality.

### Classification of hybrid classes and position of geographic clines support hybrid zone age and movement

We found only later generation hybrids and backcrosses in the oldest hybrid zone (T1) while F1s, later generation hybrids and back crosses were found in the younger hybrid zones of T2 & T3 (Fig 4C). In the oldest hybrid zone, late generation backcrosses and pure yellow *icteronotus* birds occurred at the easternmost localities in the Cauca Valley (A & B), while in T2 and T3, F1 hybrids were limited to localities around the genetic center of the hybrid zone and pure yellow *icteronotus* birds only occurred in the westernmost allopatric localities in the Pacific (Fig 4C).

Similar to the location of the overall genetic center of the hybrid zone (Fig 4A), the genomic and plumage cline centers were significantly displaced eastward from the elevation transition for both sexes in the oldest (T1) and intermediate (T2) transects (Fig 5; Table 2). In the youngest transect (T3), the plumage color cline for females was only slightly displaced eastward from the elevation transition, but the genomic clines for both sexes and the plumage color cline for males were not. On the other hand, clines for all traits in both sexes were only coincident with the tree landcover transition for T1, except for the female plumage color cline which was significantly displaced east. In T2 and T3 cline centers for all traits in both sexes were located west of the tree landcover transition.

**Figure 5.**
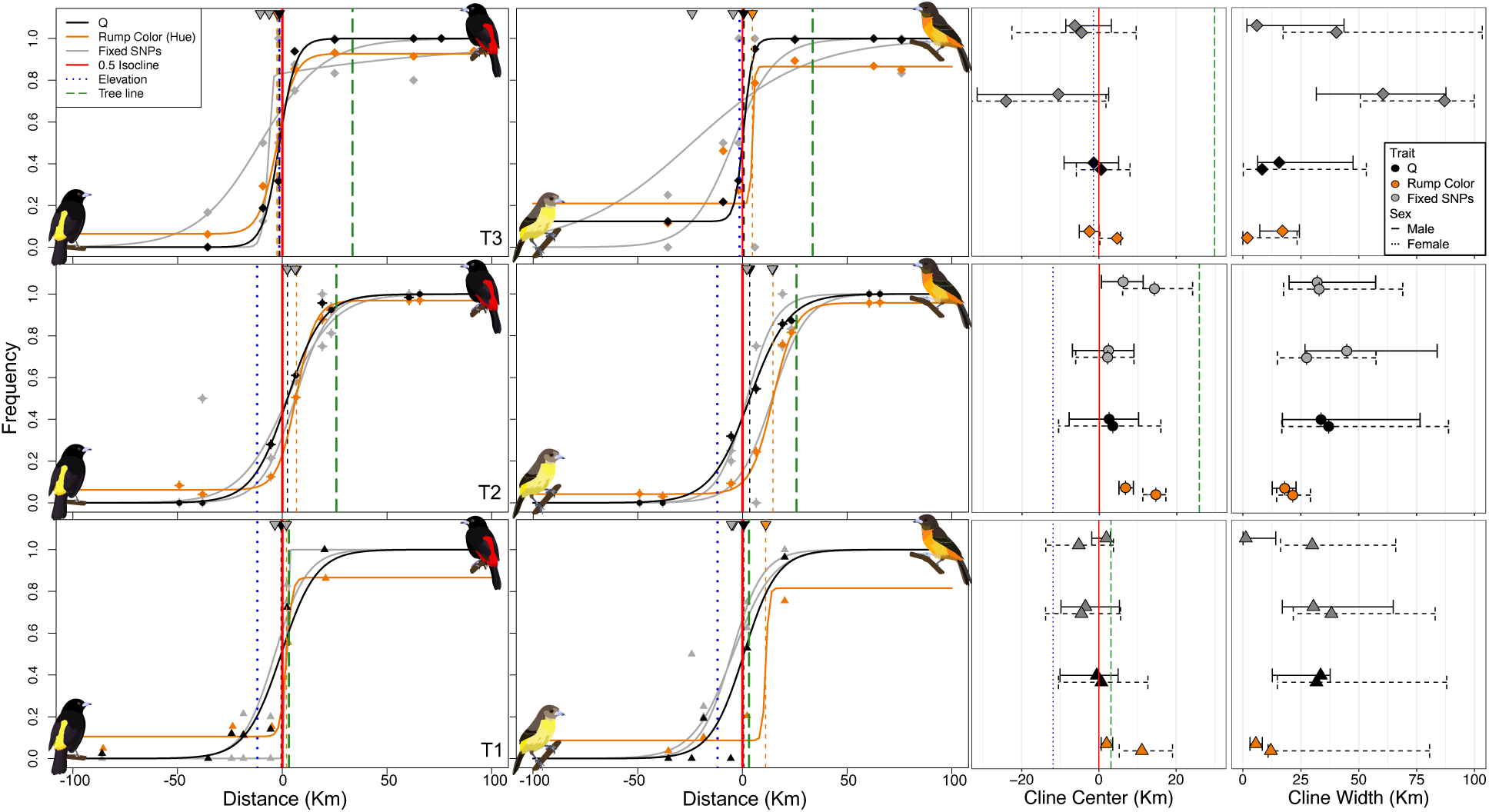
Geographic cline analysis reveals chronological displacement of the hybrid zones away from elevation transition towards the tree landcover transition. A) Best supported cline models for Q (based on ADMIXTURE proportions for K=2; black), rump color hue (orange) and two fixed SNPs (gray) for males (left) and females (right). Distance indicates the position along the transects, which are all centered at 0 km representing the genetic center of the hybrid zone (red; location where the transects intercept the isocline where the hybrid index = 0.5 in the geographic admixture surface). B) Cline center estimates and 95% confidence intervals for each trait in males (solid line) and females (dashed lines). Genomic and color clines are displaced from the elevation transition (blue dotted line) in T1 and T2, but not in T3. In contrast, all clines are coincident with the tree landcover transition (green dashed line) only in T1. C) Cline width estimates and 95 % confidence intervals.

Clines for plumage color in both sexes were consistently steep and narrower than expected under neutral diffusion, even if the dispersal distance was as low as 1km and hybridization began as recently as 1000 ybp (ca. 49 km). However, cline centers for plumage color were coincident with their respective sex-specific genomic cline (Q) in all transects; thus, we did not find evidence of differential introgression of plumage color in either sex (Fig 5).

### Genomic clines for the only shared outlier region also predict plumage color

After filtering, we kept 830 SNPs in T1, 898 in T2 and 558 in T3. The hybrid index calculated with *gghybrid* based on these sites confirmed our assignment of pure yellow *icteronotus* (S0; T1 = 14, T2 = 16, T3 = 8) or red *flammigerus* (S1; T1 = 11, T2 = 18, T3 = 28) populations (S10 Fig). Hybrid indices also revealed that the admixed individuals had a strong skew towards the yellow *icteronotus* subspecies ancestry in T1, likely reflecting later generation backcrosses and recapitulating the findings from our triangle plots (S10A Fig). However, the magnitude and direction of introgression based on genomic clines were highly variable across transects (Table 3; S10-12 Figs). T1 had very similar proportions of *u* (center) outliers displaced into the yellow *icteronotus* (51%) or red *flammigerus* (49%) background. In contrast, T2 and T3 had slightly higher proportions of outliers displaced into the red *flammigerus* background (T2 = 55%, T3 = 52 %; Table 3; S10-12 Figs). Chromosome-specific introgression patterns were also highly variable and neither of the fixed SNPs in chromosomes 6 or Z showed evidence of differential introgression (Fig 5 and 6A; S11-12 Figs).

**Figure 6.**
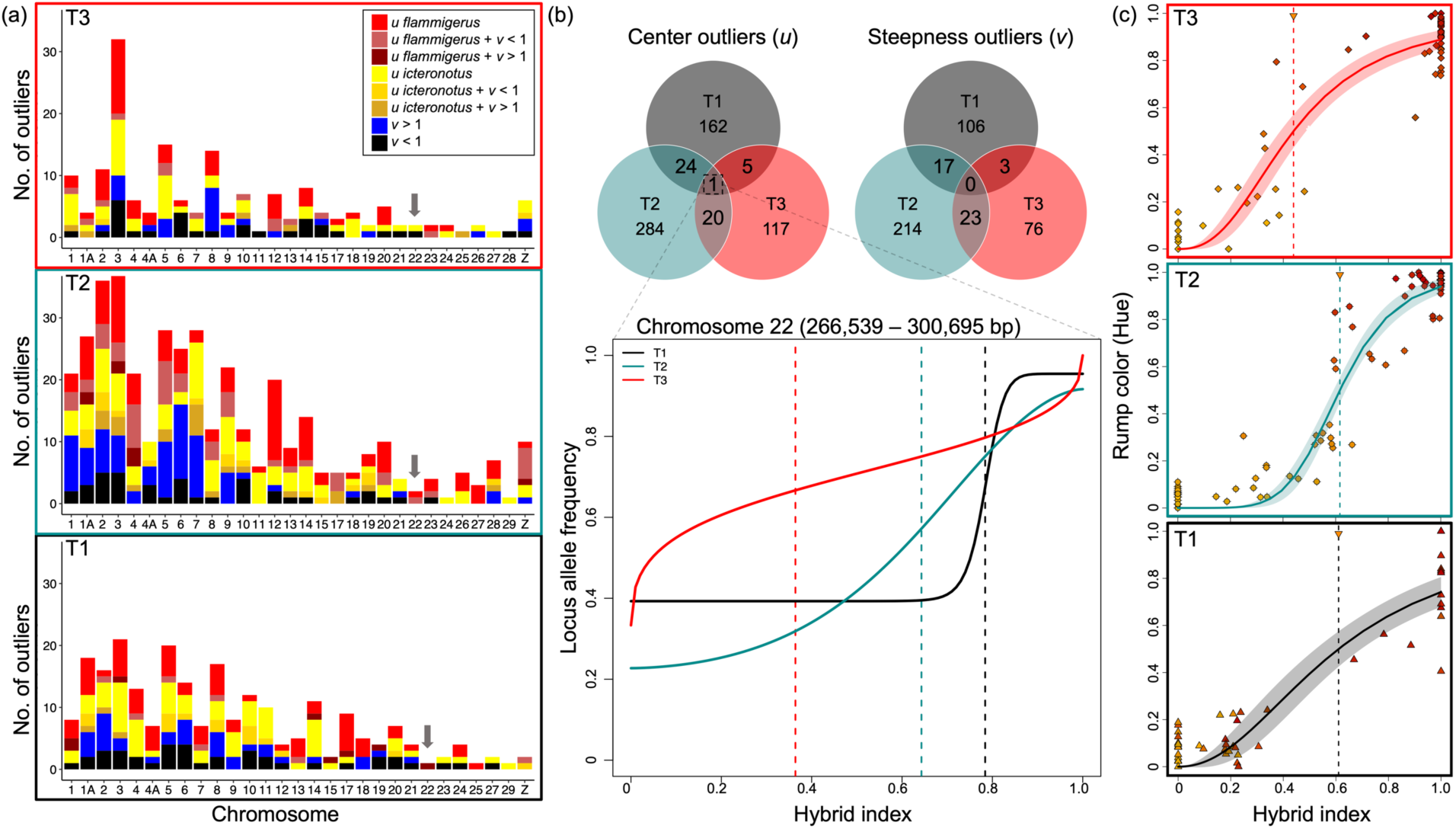
Only shared genomic cline outlier region across transects predicts plumage color. A) Low levels of outlier sharing across transects inferred from the number of loci with clines that deviated from the null expectation in cline center (*u*) or steepness (*v*) along the genome. Number of clines per chromosome that were significantly displaced into the *R. f. flammigerus u* > 0.5 (red) or *R. f. icteronotus u* < 0.5 (yellow) backgrounds. Dark red or yellow indicates that clines for these loci were also significantly steeper than expected *v* > 1 and light red or yellow indicates that they were also significantly wider than expected *v* < 1. Clines that were only significantly steeper than expected are plotted in blue or significantly wider than expected in black. B) Total number of outliers for center and steepness across transects. Only one 100Kb region contains shared center outliers and introgression patterns for these loci are consistent with the time of onset of hybridization: cline for T1 is displaced farther (black), cline for T2 in the middle (cyan) and cline for T3 (red) is not yet displaced into the *R. f. flammigerus* background. C) Center of plumage color genomic clines match the position of the clines for loci in the only shared outlier region.

**Table 3.** Genomic cline outliers for center *(u*) and steepness (*v*).

| Transect | Cline center ($u$) | *Icteronotus*<br>$u < 0.5$ | *flammigerus*<br>$u > 0.5$ | Cline steepness ($v$) | $v > 1$ | $v < 1$ | Both ($u$ & $v$) |
| --- | --- | --- | --- | --- | --- | --- | --- |
| T1 | 162 | 84 | 78 | 106 | 56 | 50 | 33 (18I-15F) |
| T2 | 284 | 125 | 159 | 214 | 145 | 69 | 99 (40I-59F) |
| T3 | 117 | 56 | 61 | 76 | 40 | 36 | 20 (7I-13F) |

Total number of loci with significant deviations (outliers) for genomic cline center and/or steepness in each transect. Outlier loci were further categorized according to whether the cline center occurred in the yellow *icteronotus* ( $u < 0.5$ ) or the red *flammigerus* ( $u > 0.5$ ) background and/or if they had steep ( $v > 1$ ) or narrow clines ( $v < 1$ ). A locus may be an outlier for one or both parameters.

We found only one 100-kb region with genomic cline center outliers across all transects. The region is between 266,539 and 300,695 bp of chromosome 22 and had one SNP in each transect that showed significant *u* (center) displacement, albeit in different directions (S13-15 Tables). Further, the extent to which the genomic clines for the three outlier loci were displaced into the red *flammigerus* background was consistent with the age of the hybrid zones (Fig 6B; Table 4 & S13-15 Tables). In the older hybrid zones (T1 and T2), the cline center of the introgression outlier SNPs occurred at a hybrid index of 0.78 and 0.64 respectively, within the red *flammigerus* genomic background. In contrast, in the youngest hybrid zone (T3), the cline center occurred at a hybrid index of 0.36, within the yellow *icteronotus* genomic background. In addition, the center of the color cline of each transect mirrored the displacement of the genomic clines for loci in this region of chromosome 22, suggesting this region is likely linked to plumage color (Fig 6C; S13 Fig). Finally, our genome-wide association study (GWAS) revealed that the only loci strongly associated with rump plumage color phenotype were within 50Kb of this region of chromosome 22 (S16 Table; S14 Fig).

**Table 4.**
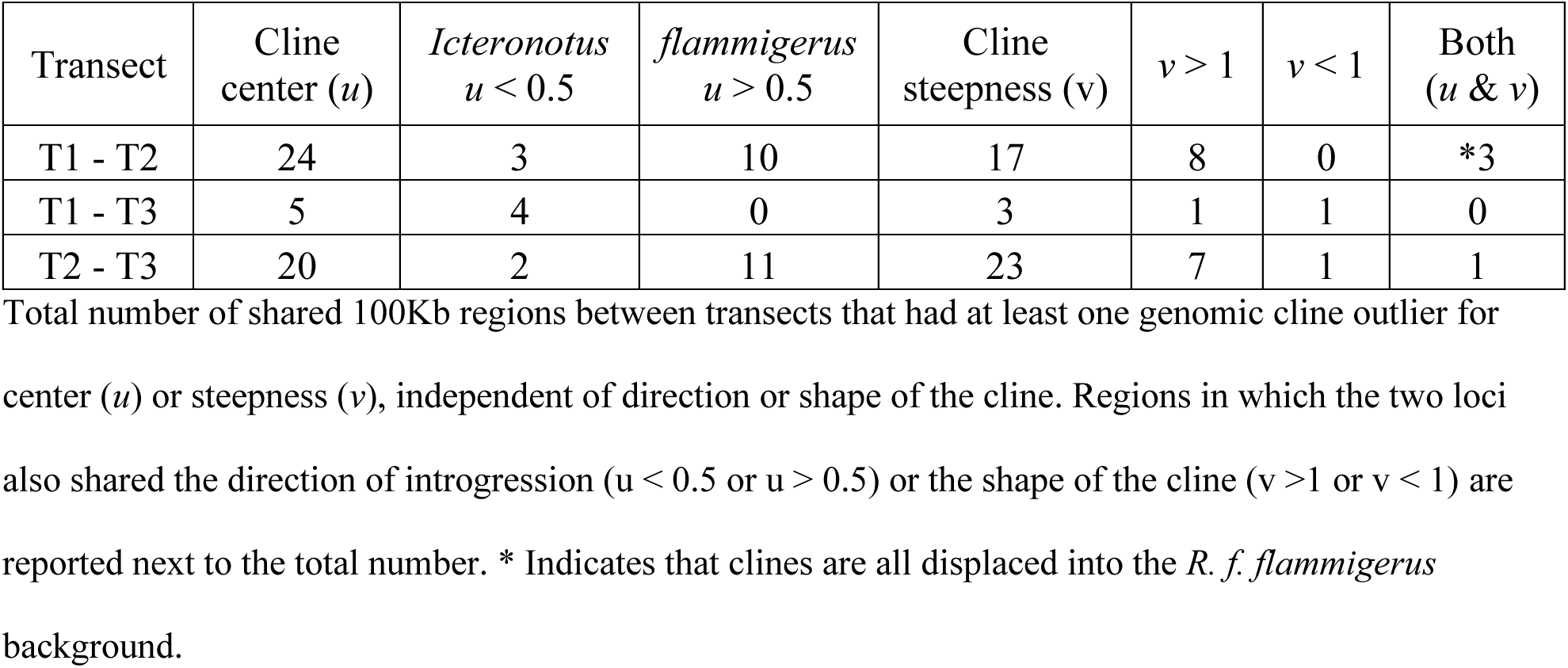
Number of shared regions with genomic cline outliers for center *(u*) and/or steepness (*v*) between transects.

| Transect | Cline center (*u*) | *Icteronotus*<br>*u* < 0.5 | *flammigerus*<br>*u* > 0.5 | Cline steepness (*v*) | *v* > 1 | *v* < 1 | Both<br>(*u* & *v*) |
| --- | --- | --- | --- | --- | --- | --- | --- |
| T1 - T2 | 24 | 3 | 10 | 17 | 8 | 0 | *3 |
| T1 - T3 | 5 | 4 | 0 | 3 | 1 | 1 | 0 |
| T2 - T3 | 20 | 2 | 11 | 23 | 7 | 1 | 1 |

Total number of shared 100Kb regions between transects that had at least one genomic cline outlier for center (*u*) or steepness (*v*), independent of direction or shape of the cline. Regions in which the two loci also shared the direction of introgression (*u* < 0.5 or *u* > 0.5) or the shape of the cline (*v* > 1 or *v* < 1) are reported next to the total number. \* Indicates that clines are all displaced into the *R. f. flammigerus* background.

## DISCUSSION

We sampled transects across three low-elevation passes that cut through the Western Andes creating separate hybrid zones between the yellow *icteronotus* and red *flammigerus* subspecies of the Flame-rumped Tanager. Across transects we observed parallel patterns of phenotypic divergence and ecological gradients, low levels of population structure throughout the ranges of parental populations, as well as confirmation that each hybrid zone was formed independently. Further, changes in effective population size over time and the magnitude and directionality of gene flow were also replicated across transects. Specifically, we found disproportionate gene flow from the yellow *icteronotus* subspecies into the hybrid zones and the red *flammigerus* subspecies’ range. However, despite similar demographic parameters and asymmetric introgression patterns, hybrid zones differed in age, which influenced the extent of geographic and genomic cline displacement. Additionally, the only shared outliers across all transects were in a genomic region that predicts plumage color, and the extent to which this has penetrated the red *flammigerus* background was consistent with the age of the hybrid zones (*i.e*., greater displacement in older zones). Still, clines for plumage traits were consistently narrow regardless of the shift in cline center over time, suggesting rump color plays a role in reproductive isolation. Overall, despite heterogeneous ancestry and introgression patterns across the genome for the three replicate hybrid zones, we found that a few key loci can maintain long-term stability of reproductive isolation, and similar environmental and demographic contexts has produced repeatable patterns of hybrid zone movement.

### Divergence with gene flow in *R. flammigerus*

Hybridization along the slopes of the Western Andes of Colombia has been documented across diverse taxa, including butterflies (Arias et al., 2008, 2012) and poison dart frogs (Márquez et al., 2020; Medina et al., 2013). Clustering of multispecies hybrid zones indicates that this biogeographic region of the Andes is likely a suture zone (Wait & Peñalba, 2024), where hybrid zones are products of secondary contact (Arias et al., 2012) or primary differentiation along ecological gradients (Arias et al., 2008). Previous studies suggested that the hybrid zone between red *flammigerus* and yellow *icteronotus* formed as a result of secondary contact after expansions from populations that diverged in refugia during the Quaternary (Morales-Rozo et al., 2017). The results of our demographic modeling with genomic data, however, provide a more complex scenario. We hypothesize that climatic fluctuations in the Pleistocene connected the eastern and the western slopes of the mountain through low elevation mountain passes, allowing for dispersal events and subsequent differentiation in isolation, as reported in other species (Cadena et al., 2016; Haffer, 1967; Ortego et al., 2023). While populations continued to diverge, intermittent gene flow likely occurred between populations mediated by changes in vegetation and climate (Cadena et al., 2016; Ramírez-Barahona & Eguiarte, 2013; Smith et al., 2014; Willis et al., 2004). Then, the hybrid zones became well-established after dramatic increases in population size and range expansions of both subspecies, resulting in an increase in gene flow along altitudinal gradients on the slopes of the Western Andes. This hypothesis is further supported by our estimations of changes in effective population size over time, which indicated that all populations experienced a crash in size around 6,000 −10,000 ybp, followed by a rapid expansion (Fig 3B). Our population size approximations are coincident with our estimate of the time of onset of hybridization for the oldest transect (ca. 6,123 ybp; Fig 3A) and with niche modeling from previous studies, which indicated that the distributions of the two subspecies were in contact at least 6000 ybp (Morales-Rozo et al., 2017). Studies on the demographic history of human populations in the same region of South America report similar drastic population size crashes in the Holocene (4200 – 10000 ybp), followed by recovery periods and subsequent stabilization (Goldberg et al., 2016; Riris & Arroyo-Kalin, 2019). In addition, dramatic population size changes across different organisms has been attributed to extreme climatic events that drastically changed humidity and temperature conditions and, consequently, the altitudinal ranges of forest ecosystems in the region (Cadena et al., 2016; Haffer, 1967; Ramírez-Barahona & Eguiarte, 2013; Riris & Arroyo-Kalin, 2019).

Time 1 (“Tm1”) in our demographic models may reflect either a more recent split or more recent sustained gene flow between subspecies across transects (Fig 3A; S9 Fig). However, because gene flow can shorten the time to coalescence for neutral loci and our divergence-with-gene-flow models allow gene flow from the moment of population split, we interpret Tm1 as the time of formation of a stable hybrid zone (Fig 3A). That is, rather than suggesting that there were three parallel divergence events (*i.e*., origin of yellow *icteronotus* subspecies) that occurred at different times, variation in the timing of sustained gene flow (*i.*e., hybrid zone formation) can explain why we recovered a more recent Tm1 in the middle and northern hybrid zones. Additional lines of evidence, including hybrid classification from triangle plots, geographic cline displacement from environmental transitions and genomic cline introgression (see discussion below), support our inference that the onset of sustained hybridization varied across the three transects.

### Long-term stability in reproductive isolation amid hybrid zone movement

Because plumage color and genomic ancestry clines are displaced away from the elevation transition in the older T1 and T2 hybrid zones but not in the younger T3 hybrid zone, we attribute this pattern to temporal differences in hybrid zone movement rather than elevation acting as a selective pressure that confines the hybrid zone to an ecotone in T3 (Fig 4B). We find the latter hypothesis to be unlikely because environmental gradients are replicated across transects and there is low population structure between parental populations, which reduces the chance for local adaptation (Bolte et al., 2024; Cutter, 2012). In addition, because dispersal abilities and demographic parameters are similar, the lack of displacement cannot be attributed to differences related to population density across transects. Thus, since hybridization began first in T1 then T2, these hybrid zones show cline displacement away from the elevational transition due to prolonged effects of selection and drift, while in T3 the hybrid zone remains aligned with the elevational transition due to more recent onset of sustained hybridization. Because climate change is expected to drive hybrid zone movement to higher elevations (Freeman et al., 2018), we expect eventual cline displacement for this younger hybrid zone. This inference is further supported by our hybrid classification with triangle plots, where the oldest hybrid zone only had later generation hybrids and backcrosses with the yellow *icteronotus* form (Fig 4C), while T2 and T3 had more F1s found almost exclusively near the center of the hybrid zone. In addition, we recovered the same pattern of cline displacement contingent on the time of onset of hybridization with genomic cline analysis for color and color-linked loci (Fig 6B & C). Despite no evidence of differential introgression of plumage color from geographic clines, clines for this trait in both sexes were consistently steeper and narrower than expected under neutral diffusion, which suggests plumage color is involved in reproductive isolation. Taken together, the replicated transects with progressive cline center displacement away from the elevation transition but closer to the tree landcover transition suggest that hybrid zones are moving upwards and east into the red *flammigerus* background through time.

Repeated sampling over the past century in the oldest transect (T1), revealed a consistent eastward and upslope shift of the hybrid zone over time (Morales-Rozo et al., 2017). Leveraging our replicate hybrid zones, we further explored the factors driving the hybrid zone movement in this system. Our results suggest that the *R. flammigerus* hybrid zone movement is a result of demographic processes amplified by climate change and anthropogenic disturbances. Sibley (1958) proposed that lack of evidence of selection against hybrids, deforestation along the main roads creating ideal corridors to connect the ranges of both subspecies and larger population sizes in the yellow *icteronotus* would result in increased gene flow from coastal to interior populations. Consistent with Sibley’s hypothesis, we found evidence of larger population sizes in the yellow *icteronotus* subspecies and hybrid zone movement from primary to secondary canopy cover. Further, increased temperatures in the last century have driven hybrid zones upslope toward cooler conditions across diverse taxa (Freeman et al., 2018). However, the observations that genomic differentiation is maintained between subspecies, clines are consistently narrow irrespective of time of onset of hybridization, and the color cline was narrower compared to the genomic ancestry cline for males suggest that there is selection against hybrids that is not weakening with time. Hence, we speculate that the hybrid zone will remain stable in shape, but movement will continue as subspecies are asymmetrical in population size and as they adjust to altered climate and habitats. Finally, social selection via male-male dominance could also contribute to hybrid zone movement, because our preliminary behavioral experiments indicate that yellow *icteronotus* birds are more aggressive than their red *flammigerus* counterparts potentially conferring reproductive advantages in the hybrid zone (Castaño et. al Unpublished data).

### Genetic architecture of plumage color

We found that a highly differentiated region of chromosome 22 (S4 Fig) is an outlier in all three transects (Fig 6) and is also the most significant predictor of plumage color in a GWAS (S14 Figs). Two genes are located within 100kb of the outlier SNPs: *OGDH* and *RETSAT*. *OGDH* codes for a 2-oxoglutarate dehydrogenase, which is involved in tricarboxylic acid cycle in the mitochondria, and so it may mediate oxidative stress and energy metabolism related to red ketocarotenoid metabolism (Hill et al., 2019; Sharma & Kumar, 2019; Simons et al., 2012). *RETSAT* codes for a retinol saturase and has been linked to carotenoid-based spectral tuning of the avian retina (Toomey et al., 2016), beta-carotene-mediated coloration in lizards (San-Jose et al., 2013), and shifts in carotenoid-based coloration in fish (Salis et al., 2019, Ahi et al., 2020) and frogs (Linderoth et al., 2023). On-going studies using whole-genome sequence data will provide a robust test of the role of *RETSAT*, or other candidate color genes potentially missed by the reduced-representation approach used in this study, in mediating plumage color in *Ramphocelus*.

### Insights from replicated hybrid zones

Despite similar ecological and demographic conditions, we found high variability in genomic cline parameters across the genome—except for a single region linked to plumage color, which likely contributes to reproductive isolation. Differences in hybrid zone age influenced the degree of outlier sharing across replicate zones, though a few key traits consistently maintained reproductive isolation over time. Overall, while locus-specific patterns were heterogeneous, the magnitude and directionality of neutral introgression was predictable under similar demographic conditions and for traits involved in reproductive isolation. Together, our work highlights the interplay of both stochastic and deterministic processes in mediating the repeatability of patterns of introgression in replicate hybrid zones.

## Acknowledgments

Santiago Monroy and Manuel Sánchez provided invaluable fieldwork and logistic support. We thank Elsie Shogren and Diego Ocampo for advice on data analysis and for helpful comments on earlier versions of this manuscript. We are grateful to SELVA, ProAves Foundation, Corporacion Autónoma Regional de Risaralda (CARDER), Hacienda Venecia, Ecoparques de Manizales, Ecoparque el Salado and Consejo Municipal de Tutunendó among many other Colombian agencies and local businesses that allowed us to collect samples for this project. Daniel Mejía, Blas Cárdenas, Octaviano, Valeriano Mena, Gustavo Echeverry, Ivan Solis, Gabriel Moscoso, Marmato, Daisy Gómez and Paula Saravia assisted with field work and logistic support and were direct links to local communities that allowed us to collect samples in their land. We thank the Collection of Genetic Resources of the Luisiana State University Museum of Natural Science for loaning tissue samples under their care (LSUMNS invoice 12163, 12345). We thank CIRC (Center for Integrated Research Computing) at the University of Rochester for useful advice and resources for data analysis.

## AUTHOR CONTRIBUTIONS

Conceptualization: MIC and JACU. Field work, lab work and data curation: MIC. Formal analysis and methodology: MIC and EC. Funding acquisition: MIC and JACU. Project Administration: MIC, CDC and JACU. Resources: CDC and JACU. Software: MIC and EC. Visualization: MIC. Supervision: JACU. Writing – original draft: MIC and JACU. Writing – review & editing: MIC, CDC and JACU. All authors read, reviewed and approved the final manuscript.

## BENEFIT-SHARING STATEMENT

Benefits Generated: A research collaboration was developed with scientists from the countries providing genetic samples and collaborators are included as co-authors. We supported local communities by consulting with community councils and hiring local and indigenous guides for assistance with field work. The contributions of all individuals and institutions to the research are described in the acknowledgments. Outcomes of this study will be disseminated in ornithological conferences and museum exhibits in Colombia. In addition, benefits from this research accrue from the sharing of our data and results on public databases as described above.

## DATA AVAILABILITY STATEMENT

The reference genome assembly, annotation GFF file, raw RNAseq reads and raw unfiltered vcf file will be available on DRYAD (DOI: 10.5061/dryad.zkh1893n9). The code and custom scripts for all analyses are available on GitHub: https://github.com/micastano10/ReplicateHybridZones

