## Supporting information for "Replicate avian hybrid zones reveal the progression of genetic and trait introgression through time"

**Available in repository** (Dryad DOI: 10.5061/dryad.zkh1893n9)**:**

1. Reference genome assembly
2. Annotation GFF file
3. Raw RNAseq reads
4. Raw unfiltered vcf file
5. S1 & S16 Tables.
6. Code and custom scripts (GitHub: https://github.com/mi-castano10/ReplicateHybridZones)

**S16 Table.** Results from a genome-wide association study (GWAS) with individuals from all transects combined, to identify regions of the genome associated with rump color (hue). **Dryad** 36
Supplementary figures ........ 37–51

### SUPPLEMENTARY MATERIALS AND METHODS

#### Genetic and phenotypic sample collection

We sampled both males and females from transects across three independent contact zones between the two taxa. The three contact zones occur along low mountain passes where roads connect the western and eastern flanks of the Cordillera Occidental in Valle del Cauca, Risaralda and Antioquia (Fig 1B). Sampling for the southern transect (T1) was conducted in 2007-2010 over a 140 km distance by road between the cities of Cali and Buenaventura, Valle del Cauca (Morales-Rozo et al., 2017). For T1, we obtained feather and tissue samples for 58 specimens deposited in the Museo de Historia Natural de la Universidad de Los Andes (ANDES). For the central (T2) and northern (T3) transects, we collected samples between November and February of 2022 and 2023. In T2, we sampled 80 birds over a 199 km distance by road between the cities of Manizales, Caldas and Tadó, Chocó. In T3 we sampled 78 birds over a 236 km distance by road between the towns of La Ceja, Antioquia and Tutunendó, Chocó. We used a combination of incidental and playback-stimulated mist-netting to capture birds. Then, we collected blood and feather samples, and took morphological measurements, including bill width, bill height, bill length, wing length, tail length, tarsus length and weight. We stored blood samples in Longmire's buffer (Longmire et al., 1997) and plucked 5-10 feathers from the rump. In 2023, we opportunistically plucked young feather follicles from molting birds, stored them in 2ml of RNAprotect and kept them at low temperatures for the duration of the field season. To prevent resampling, we banded each bird with a uniquely numbered aluminum band (National Band and Tag Company, USA). All protocols were approved by the University of Rochester Committee on Animal Resources (Protocol: 102459/2021-003). Sample collection and exportation permits in Colombia were issued by the Agencia Nacional de Licencias Ambientales (ANLA) under the Permiso Marco de Recolección No. IDB 0359 to Universidad de los Andes (resolución 1177 del 09 de octubre de 2014).

#### DNA and RNA extraction and sequencing

High molecular weight DNA from a single female *R. f. icteronotus* sampled in a locality distant from any hybrid zone in Playa de Oro, Chocó, Colombia (T2 – locality G) was isolated using the Qiagen MagAttract kit (QIAGEN, shanghai, China) following the manufacturer's instructions. One SMRTbell library of circular consensus sequencing (CCS) was constructed and sequenced on three PacBio Sequel II SMRT cell platforms in CCS mode. This mode calls consensus reads from subreads that are generated after multiple passes of the enzyme around the circularized template (Hon et al., 2020). In addition, we extracted RNA from 5 emerging feather follicles of the same female using the Monarch Total RNA Miniprep Kit (New England Biolabs, USA). RNA samples were received and validated via standard quality control procedures at Novogene Co. Ltd. Sacramento, CA. Sequencing was performed in a NovaSeq PE 150 platform, using an mRNA library preparation with standard poly A enrichment.

We extracted genomic DNA from a total of 252 birds (S1 Table) using Qiagen DNeasy Blood and Tissue Extraction Kit (Qiagen, USA) following the manufacturer's instructions, with the optional addition of 4 µL of RNase to remove any potential RNA contaminants. We sent the samples for genotype-by-sequencing (GBS) (Elshire et al., 2011) to the University of Wisconsin Biotechnology Center. Samples were digested with the *Ape*Ki restriction enzyme, and short fragments from throughout the genome were sequenced on an illumina NovaSeq 6000 2 X 150 Shared Sequencing. We sent one 96-well plate with samples from T1 and other species from the genus *Ramphocelus* for sequencing in 2021, and two plates with the rest of the samples for T2 and T3 in 2023. The 2021 plate yielded 627 million sequences, and the 2023 plates for T2 and T3 yielded 550 and 523 million sequences, respectively.

#### *De novo* chromosome scale genome assembly for *R. f. icteronotus*

Prior to assembly we assessed the estimated genome size, rate of heterozygosity and repetitive element content of our genome. We used Meryl (v1.4.1;(Rhie et al., 2020)) to count and compute a histogram of k-mer frequencies from the raw adapter trimmed PacBio Hifi reads using count k = 21. We uploaded the output of Meryl to the Galaxy web platform and used the public server at usegalaxy.org to run GenomeScope2.0 with the following parameters: --ploidy 2 –kmer_length 21 (Ranallo-Benavidez et al., 2020; The Galaxy Community, 2024). The model indicated that *R. f. icteronotus* has an estimated genome size of 1,164,347 bp, with 0.88% heterozygosity levels and around 20% of the genome composed of repetitive elements (S1 Fig). We generated a phased genome assembly using the -primary flag and default parameters of hifiasm (v0.16.1;(Cheng et al., 2021)). The primary assembly was 1,394,387,738bp long, assembled in 347 contigs with an N50 of 71.1 Mb and a GC content of 44.97%. We used Blobtools2 from the BlobToolKit suite (Challis et al., 2020) to screen the assembly for contamination. First, we performed a BLASTn (Camacho et al., 2009) search of our assembly against the general RefSeq blast -nt database using the following parameters: -outfmt '6 qseqid staxids bitscore std' -max_target_seqs 1 -max_hsps 1 -num_threads 12 -evalue 1e-25. Then we used minimap2 (v2.17;(Li, 2018)) to map the raw reads back to the assembly. We used the function blobtools –add to create a BlobDir database that included the bast output (hits file) and the read coverage (bam file). Lastly, we employed the online viewer (blobtools host 'pwd') to make a Blobplot with the hits and respective coverage of our contamination scan and removed 11 contigs that didn't match the order Passeriformes. After removing spurious contigs, we assembled the mitochondrial genome de novo using MitoHiFi (v3.2; (Uliano-Silva et al., 2023)) with default parameters and the Black-throated Flowerpiercer (*Diglossa brunneiventris*; NCBI Accession: NC_062466.1) mitochondrial genome as a reference. The final mitochondrial genome for *R. f. icteronotus* was 16,784 bp long, with 37 genes. We used BLASTn (Camacho et al., 2009) to identify and remove other mitochondrial contaminant sequences present in the nuclear genome and added the contig with the MitoHiFi mitochondrial genome at the end of the *R. f. icteronotus* assembly. We evaluated assembly quality and completeness using Benchmarking Universal Single-copy Orthologs (BUSCO v5.8; (Manni et al., 2021)) with the passeriformes_odb10 database, available in the public server at usegalaxy.org (The Galaxy Community, 2024). Finally, to achieve a chromosome-level genome assembly, we scaffolded our assembly using the chromosome-level genomes of the Black-throated Flowerpiercer (*Diglossa brunneiventris*; NCBI assembly ID: GCA_019023105.1) and the Zebra finch (*Taeniopygia guttata;* NCBI assembly ID: GCA_003957565.4). We mapped our assembly to the reference genomes using minimap2 (Li, 2018), and ordered and numbered our contigs first according to identity with *D. brunneiventris* and then *T. guttata* for the micro chromosomes that were not present in the *D. brunneiventris* assembly. We used custom python scripts to assemble the contigs that mapped to the same reference chromosome into scaffolds by adding 100 Ns in between

**Read processing, variant calling and filtering**

We used Tassel (v5.2.65; (Glaubitz et al., 2014)) to process the raw reads from the three plates combined, which included samples from our three transects, yellow *icteronotus* samples from Ecuador and Panama, and samples from other species in the genus *Ramphocelus* (S1 Table). First, we demultiplexed the reads and called variants following the GBSv2 SNP discovery and production pipeline. We aligned the short reads to our reference genome assembly for a female *R. f. icteronotus* with Bowtie2 (Langmead & Salzberg, 2013) and we used the default parameter settings for the rest of the pipeline, except for the minimum quality score (default mnQS = 0), which we increased to 20. After read demultiplexing, mapping and variant calling with tassel, we obtained 2,598,209 genome-wide SNPs for 254 individuals. We then used VCFtools (v0.1.16; (Danecek et al., 2011)) to filter our main dataset based on the following criteria: remove samples with mean depth < 1, include only biallelic SNPs with no indels and site depth ≥ 4, and exclude SNPs with > 0.25 missing data. The first round of filtering kept 229 individuals and 53,559 SNPs (dataset A). Then we proceeded to subset individuals only from our species *R. flammigerus* (Ecuador and Panama and the three sampling transects) and applied a minor allele count = 1 filter to keep sites that were polymorphisms between *R. flammigerus* subspecies. The second round of filtering kept 215 individuals (Ecuador, Panama and other locations in Colombia = 19, T1 = 47, T2 = 79, T3 = 70) and 50,868 SNPs (dataset B), which we used for $F_{ST}$ calculations, classification of hybrid classes, genomic clines and a GWAS for which we imputed missing data with BEAGLE (v5.5; (Browning et al., 2018)). Starting from dataset B, we conducted a third round of filtering to include only sites with minor allele frequency > 0.05 and thinned so that SNPs were ≥100 bp apart (one SNP per read) to reduce effects of linkage. This third round of filtering retained 14,123 SNPs (dataset C), which we used as input for ADMIXTURE (Alexander et al., 2009), estimating effective migration surface (EEMS;

(Petkova et al., 2015)) and principal component analysis. Finally, we implemented a last round of stringent filtering for analysis related to demographic history. Starting from dataset B (no minor allele filtering and no thinning), we further excluded all individuals with > 30% missing data, SNPs that were missing in > 10% of the individuals, and all non-autosomal SNPs because sex chromosomes can have different demographic histories than the rest of the genome (Wilson Sayres, 2018). We calculated the KING relatedness coefficient using the -*relatedness2* function of VCFtools (v0.1.16; (Danecek et al., 2011)) and, for all combinations in which the kinship coefficient was > 0.25 (first degree relatives), we removed the individual in each pair that had a greater proportion of missing data. This last round of filtering retained 12,103 SNPs and 173 Individuals (dataset D) which we further split by transect and used for estimations of effective population size over time with popsizeABC ((Boitard et al., 2016)). We used a thinned version of dataset D to obtain a complete site-frequency spectrum (SFS) that was unaffected by linkage and not truncated by minor allele frequency filtering. The thinned version retained 5006 SNPs that were used as input for demographic modeling with fastsimcoal2 (v27.09; (Excoffier et al., 2021)) and SFS based effective population size estimations with stairwayplot2 (Liu & Fu, 2020). A summary of the four datasets and the analysis they were used for are available in S3 Table. Overall, having a centralized dataset of SNPs for all transects allowed us to make accurate inferences about replicated patterns.

**Phenotypic and genomic ancestry assignment**

We used 14,123 unlinked SNPs (dataset C; Table S3) as input for ADMIXTURE v.1.3.0 (Alexander et al., 2009) to identify the most likely number of distinct population clusters (k = 1–5) and assign genomic ancestry. We executed four runs; one for all *R. flammigerus* individuals (including samples from the three transects, Panama, Ecuador and other regions of Colombia) to obtain a range-wide ancestry assignment, and one for each transect separately. We converted VCFs to BED format with PLINK v1.9 (Chang et al., 2015), ran ADMIXTURE with -C 100 -B 200 and used 10-fold cross-validation to determine the optimal number of populations (k = 2 in all runs; S4 Table).We visualized genomic ancestry as pie charts over elevation maps using the *sf* (Pebesma et al., 2024), *elevatr* (Hollister et al., 2023) and *rnaturalearth* (Massicotte et al., 2023) packages in *R* studio.

For color analysis, we measured the spectral properties of the rump, which is the most distinctive color patch between taxa, from 300 to 700nm. To standardize the background, we mounted 5 feathers of the rump of each bird in a microscope slide painted with matte black paint. We used an Ocean Optics USB2000 spectrophotometer and a PX-2 pulsed xenon light source (Ocean Optics) with the average reading set at 50, integrating time at 16ms, boxcar smoothing at 10 and correction for electric dark noise activated. The tip of the fiber-optic probe was covered by a black anodized aluminum cap angled at 45° to reduce glare (as in (Ocampo et al., 2023; Uy & Stein, 2007)). To maintain consistency, we calibrated the spectrophotometer every 3 measurements with a Spectralon white diffuse standard and a dark standard (matte black painted slide). Then we used the R package *pavo2* (Maia et al., 2019) to smooth the data and remove spectral noise while preserving the original spectral shape. We applied a Loess smoothing factor of 0.2 and corrected for any negative values in our measurements. After smoothing, we calculated 23 colorimetric variables and used hue (wavelength of maximum reflectance - H4; (Montgomerie, 2006)) as our quantitative value for color. We decided to use hue because in *R. flammigerus* male feathers have specialized microstructures that can influence color variables related to saturation (e.g., chroma) (McCoy et al., 2021). However, in pigment based coloration microstructures don´t usually change the pigment´s wavelength absorption, so they have limited effect on hue (Shawkey et al., 2006). We used the *spec2rgb* function to calculate RGB values from spectral data and used the hexadecimal code to plot the rump color phenotype in all figures.

To characterize morphological variation, we ran a principal components analysis using with the 7 morphometric characters measured in the field and used PC1 as a proxy for body and bill size. All juvenile birds were excluded from downstream phenotypic analysis (S1, 8-10 Tables). To standardize across transects, we converted values for phenotypic traits into scores that ranged from 0 to 1 for each transect independently, with the following equation: $(X_i - X_{min})/(X_{max} - X_{min})$. Here, $X_i$ corresponds to the individual value for the trait, and $X_{max}$ and $X_{min}$ correspond to the maximum and minimum trait values for each transect. Finally, we used the R package *corrplot* (Wei et al., 2024) to calculate one of Pearson's product moment correlation coefficient between genomic ancestry (Q) and rump plumage color (hue), body size (PC1) and distance along transect to determine the extent to which a given variable predicts genomic ancestry.

**Demographic history**

PopSizeABC uses Approximate Bayesian Computation (ABC) to model a single population experiencing changes in $N_e$ over time by comparing observed genetic data (*i.e.,* summary statistics related to allele frequencies and linkage disequilibrium) to simulated data under a specified demographic model (Boitard et al., 2016). In contrast, Stairwayplot2 (Liu & Fu, 2020) infers $N_e$ by comparing the observed site frequency spectrum (SFS) to expectations under a theoretical model derived from coalescent theory, that uses assumptions of neutrality, random mating, and the infinite-sites mutation model to estimate how the distribution of allele frequencies (SFS) reflects historical changes in effective population size. Stairwayplo2 was designed for GBS data and performs better when estimating population size history in the more distant past (Nadachowska-Brzyska et al., 2022). We used the results from both $N_e$ estimation methods to guide our model-based coalescent simulations with fastSIMCOAL2 (v27.09; (Excoffier et al., 2021)).

For each transect, we compared 15 models with different demographic scenarios, all with three populations for which we had previously calculated the SFS: AB, CDEF, GH (pure allopatric red *flammigerus* from localities A-B, hybrids in localities C,D,E,F and allopatric yellow *icteronotus* from localities G-H; S9 Fig). Models 1-3 were null models with no gene flow between subspecies but different modes of population divergence. Model 1 tested an allopatric model of divergence and was the only model with two populations. Model 2 tested a parapatric model of divergence, where a hybrid population was formed in the Western Andes at time of divergence. Model 3 tested an allopatric model of divergence where subspecies first diverged in isolation and then the population in the slopes of the Western Andes split from the pure allopatric red *flammigerus*. Splitting time between the allopatric red *flammigerus* population and the hybrid population would approximate the time at which the hybrid zone was formed. Under models 4-6 we tested an allopatric model of divergence, but with constant (model 4), increasing (model 5; homogenization) or decreasing (model 6; reinforcement) gene flow after secondary contact. Under models 7-9, we tested divergence with geneflow. In model 7, gene flow was constant from the moment of split, indicating plumage differentiation occurred without strong reproductive isolation. In model 8, gene flow increases after a point in time, indicating either strong pulses of admixture from allopatric populations or eroding reproductive barriers. In model 9, gene flow decreases after a point in time, which would be consistent with increasing reproductive isolation through time (E. Luzuriaga-Aveiga et al., 2021). Models 10 to 12 tested the same divergence with geneflow demographic scenarios as models 7-9, but with population expansions in the yellow *icteronotus* (model 10), the red *flammigerus* (model 11) or in both (model 12). Lastly, models 13-15 tested the same allopatric model of divergence scenarios as models 4-6, but with population expansions in the yellow *icteronotus* (model 13), the red *flammigerus* (model 14) or in both (model 15).

After finding the best model, we performed a bootstrap following the fastSIMCOAL2 manual to calculate confidence intervals and thus evaluate the certainty of our parameter estimates (Table 1). In brief, we simulated 100 SFS under the best fit model, and for each bootstrap replicate we re-estimated the demographic parameters with 30 runs of FastSIMCOAL2. Then we used the bootstrap estimates to calculate 95% confidence intervals based on the $2.5^{th}$ and $97.5^{th}$ percentiles of the parameter distributions.

**Genomic Clines**

The package *gghybrid* estimates Buerkle's genome-wide hybrid index (Buerkle, 2005) and fits Fitzpatrick's logit-logistic genomic cline models (Fitzpatrick, 2013) to identify loci with restricted or biased introgression. The null model assumes that the probability of ancestry at a specific locus is directly predicted by the hybrid index of that individual. Any locus in which the probability of ancestry deviates from the null model will be considered an outlier,  exhibiting excess cline parameter values of *u* (center), *v* (steepness) or both (Bailey, 2024).

### TABLES

**S1 Table. Sample information.**

Metadata for all samples included in the study. (Available in dryad repository - DOI: 10.5061/dryad.zkh1893n9).

**S2 Table. Reference genome assembly statistics.**

| Assembly | Raw assembly | Final scaffolded assembly |
|---|---|---|
| # contigs (>= 0 bp) | 347 | 109 |
| # contigs (>= 1000 bp) | 347 | 109 |
| # contigs (>= 5000 bp) | 347 | 109 |
| # contigs (>= 10000 bp) | 347 | 109 |
| # contigs (>= 25000 bp) | 311 | 100 |
| # contigs (>= 50000 bp) | 213 | 77 |
| Total length (>= 0 bp) | 1394387738 | 1393652517 |
| Total length (>= 1000 bp) | 1394387738 | 1393652517 |
| Total length (>= 5000 bp) | 1394387738 | 1393652517 |
| Total length (>= 10000 bp) | 1394387738 | 1393652517 |
| Total length (>= 25000 bp) | 1393643733 | 1393484582 |
| Total length (>= 50000 bp) | 1390139663 | 1392694278 |
| # contigs | 347 | 109 |
| Largest contig | 163891001 | 166932742 |
| Total length | 1394387738 | 1393652517 |
| GC (%) | 44.97 | 44.97 |
| N50 | 71109032 | 75408491 |
| N75 | 18099118 | 22266477 |
| L50 | 7 | 7 |
| L75 | 19 | 17 |
| # N's per 100 kbp | 0 | 1.63 |

Summary of the *Ramphocelus flammigerus icteronotus* reference genome statistics, including both the raw and final scaffolded assembly.

**S3 Table. Datasets and analysis summary.**

| Dataset | MAF | Thinning | #SNPs | Individuals included | Analysis |
|---|---|---|---|---|---|
| Dataset A | No | No | 53,559 | 229 - All *Ramphocelus* | SNP calling |
| Dataset B | No | No | 50,868 | 215 - All *R. flammigerus* | Overall Fst |
| | No | No | 50,868 | 196 - Transects only | Fst, Introgress, genomic clines, Gwas |
| Dataset C | 0.05 | Yes --100 | 14,123 | 215 - All *R. flammigerus* | Overall ADMIXTURE |
| | 0.05 | Yes --100 | 14,123 | 196 - Transects only | Admixture, PCA, EEMS |
| Dataset D | No | No | 12,103 | 45 - Only pure parentals | PopsizeABC |
| | No | Yes --100 | 5,006 | 54 - Transects only | easySFS, Stairwayplot2, FastSIMCOAL2 |

Dataset name, filtering parameters, number of SNPs and individuals kept and the analysis they were used for.

**S4 Table. ADMIXTURE cross-validation error.**

| k | All | T1 | T2 | T3 |
|---|---|---|---|---|
| 1 | 0.51966 | 0.54851 | 0.51548 | 0.51104 |
| **2** | **0.47791** | **0.53215** | **0.49059** | **0.50889** |
| 3 | 0.47866 | 0.57377 | 0.51557 | 0.54473 |
| 4 | 0.47993 | 0.62933 | 0.54906 | 0.60778 |
| 5 | 0.49005 | 0.69611 | 0.56893 | 0.63185 |

Cross validation error for ADMIXTURE of K = 1-5 ran using all available *R. flammigerus* samples (All) or with samples from each transect independently.

**S5 Table. Model comparison using Akaike information Criteria (AIC) for alternative demographic scenarios tested with FastSIMCOAL2 in T1.**

| Model | Scenario | K | AIC | ΔAIC | wAIC |
|---|---|---|---|---|---|
| **Model7** | **Divergence with geneflow** | **9** | **16062.07** | **0** | **0.992741** |
| Model9 | Divergence with geneflow (reinforcement) | 14 | 16073.56 | 11.494 | 0.003169 |
| Model10 | Divergence with geneflow (*ict* expansion) | 13 | 16073.71 | 11.638 | 0.002949 |
| Model8 | Divergence with geneflow (homogenization) | 14 | 16075.61 | 13.538 | 0.001141 |
| Model11 | Divergence with geneflow (*flam* expansion) | 13 | 16997.74 | 935.674 | 6.57E-204 |
| Model4 | Allopatric (constant geneflow) | 12 | 17543.62 | 1481.55 | 1.93e-322 |
| Model12 | Divergence with geneflow (both expansions) | 16 | 17828.82 | 1766.754 | 0 |
| Model13 | Allopatric (*ict* expansion) | 16 | 17568.95 | 1506.878 | 0 |
| Model14 | Allopatric (*flam* expansion) | 16 | 17555.18 | 1493.108 | 0 |
| Model15 | Allopatric (both expansions) | 19 | 17570.55 | 1508.482 | 0 |
| Model1 | Allopatric (null 2 populations) | 4 | 17850.03 | 1787.958 | 0 |
| Model2 | Divergence with geneflow (null) | 5 | 17834.73 | 1772.666 | 0 |
| Model3 | Allopatric (null 3 populations) | 7 | 17829.3 | 1767.232 | 0 |
| Model5 | Allopatric (homogenization) | 17 | 17565.29 | 1503.218 | 0 |
| Model6 | Allopatric (reinforcement) | 17 | 17557.91 | 1495.844 | 0 |

Table includes the number of parameters (k), AIC values, ΔAIC (difference from the best model), and Akaike weights (wAIC) for 15 demographic models tested using FastSIMCOAL2. Input SFS was constructed using only samples only from T1.

**S6 Table. Model comparison using Akaike information Criteria (AIC) for alternative demographic scenarios tested with FastSIMCOAL2 in T2.**

| Model | Scenario | k | AIC | ΔAIC | wAIC |
|---|---|---|---|---|---|
| **Model8** | **Divergence with geneflow (homogenization)** | **14** | **14769.03** | **0** | **1.00** |
| Model9 | Divergence with geneflow (reinforcement) | 14 | 14815.58 | 46.558 | 7.76E-11 |
| Model11 | Divergence with geneflow (*flam* expansion) | 13 | 14816.28 | 47.258 | 5.47E-11 |
| Model7 | Divergence with geneflow (constant) | 9 | 14816.86 | 47.836 | 4.10E-11 |
| Model10 | Divergence with geneflow (*ict* expansion) | 13 | 14817.34 | 48.31 | 3.23E-11 |
| Model14 | Allopatric (*flam* expansion) | 16 | 16044.85 | 1275.828 | 9.07E-278 |
| Model5 | Allopatric (homogenization) | 17 | 16051.54 | 1282.518 | 3.20E-279 |
| Model4 | Allopatric (constant geneflow) | 12 | 16054.91 | 1285.884 | 5.94E-280 |
| Model15 | Allopatric (both expansions) | 19 | 16057.52 | 1288.494 | 1.61E-280 |
| Model13 | Allopatric (*ict* expansion) | 16 | 16064.8 | 1295.772 | 4.23E-282 |
| Model6 | Allopatric (reinforcement) | 17 | 16070.64 | 1301.61 | 2.29E-283 |
| Model3 | Allopatric (null 3 populations) | 7 | 16318.39 | 1549.36 | 0.00 |
| Model12 | Divergence with geneflow (both expansions) | 16 | 16336.89 | 1567.862 | 0.00 |
| Model2 | Divergence with geneflow (null) | 5 | 16368.3 | 1599.276 | 0.00 |
| Model1 | Allopatric (null 2 populations) | 4 | 17537.71 | 2768.686 | 0.00 |

Table includes the number of parameters (k), AIC values, ΔAIC (difference from the best model), and Akaike weights (wAIC) for 15 demographic models tested using FastSIMCOAL2. Input SFS was constructed using only samples only from T2.

**S7 Table. Model comparison using Akaike information Criteria (AIC) for alternative demographic scenarios tested with FastSIMCOAL2 in T3.**

| Model | Scenario | k | AIC | ΔAIC | wAIC |
|---|---|---|---|---|---|
| **Model8** | **Divergence with geneflow (homogenization)** | **14** | **13458.524** | **0** | **0.926881** |
| Model7 | Divergence with geneflow (constant) | 9 | 13463.948 | 5.424 | 0.061548 |
| Model10 | Divergence with geneflow (*ict* expansion) | 13 | 13467.348 | 8.824 | 0.011244 |
| Model9 | Divergence with geneflow (reinforcement) | 14 | 13474.426 | 15.902 | 0.000327 |
| Model11 | Divergence with geneflow (*flam* expansion) | 13 | 13516.816 | 58.292 | 2.04E-13 |
| Model1 | Allopatric (null 2 populations) | 4 | 14331.308 | 872.784 | 2.78E-190 |
| Model14 | Allopatric (*flam* expansion) | 16 | 14718.742 | 1260.218 | 2.06E-274 |
| Model4 | Allopatric (constant geneflow) | 12 | 14722.958 | 1264.434 | 2.50E-275 |
| Model13 | Allopatric (*ict* expansion) | 16 | 14723.866 | 1265.342 | 1.59E-275 |
| Model15 | Allopatric (both expansions) | 19 | 14725.652 | 1267.128 | 6.51E-276 |
| Model5 | Allopatric (homogenization) | 17 | 14726.902 | 1268.378 | 3.49E-276 |
| Model3 | Allopatric (null 3 populations) | 7 | 14944.126 | 1485.602 | 2.47e-323 |
| Model2 | Divergence with geneflow (null) | 5 | 14955.774 | 1497.25 | 0 |
| Model6 | Allopatric (reinforcement) | 17 | 14971.182 | 1512.658 | 0 |
| Model12 | Divergence with geneflow (both expansions) | 16 | 14979.582 | 1521.058 | 0 |

Table includes the number of parameters (k), AIC values, ΔAIC (difference from the best model), and Akaike weights (wAIC) for 15 demographic models tested using FastSIMCOAL2. Input SFS was constructed using only samples only from T3.

**S8 Table. Genetic and morphological sampling scheme for T1.**

| Sample ID | Latitude | Longitude | Q | Locality | Sex | Elevation | Fitted Latitude (Linear Transect) | Distance (Km) | Rump color (Hue) |
|---|---|---|---|---|---|---|---|---|---|
| RFI1199COL_T1_LG_M | 3.9635 | -77.37908 | 0.00001 | G | Male | 5 | 3.94004 | -87.13954 | 0.07558 |
| RFI1200COL_T1_LG_M | 3.9635 | -77.37908 | 0.12164 | G | Male | 7 | 3.94004 | -87.13954 | 0.00000 |
| RFI604COL_T1_LG_M | 3.943694444 | -77.36608333 | 0.00001 | G | Male | 12 | 3.93070 | -85.36651 | 0.10059 |
| RFI602COL_T1_LG_M | 3.945888889 | -77.36597222 | 0.00001 | G | Male | 24 | 3.93062 | -85.35135 | 0.02508 |
| RFI603COL_T1_LG_M | 3.945888889 | -77.36597222 | 0.00001 | G | Male | 28 | 3.93062 | -85.35135 | 0.03804 |
| RFI1211COL_T1_LF_M | 3.84847 | -77 | 0.00001 | F | Male | 13 | 3.66755 | -35.41960 | 0.08580 |
| RFI1210COL_T1_LF_F | 3.84847 | -77 | 0.00001 | F | Female | 20 | 3.66755 | -35.41960 | 0.02255 |
| RFI1195COL_T1_LF_F | 2.506472222 | -76.99925 | 0.00001 | F | Female | 1464 | 3.66701 | -35.31727 | 0.05168 |
| RFI1223COL_T1_LE_F | 3.608666667 | -76.918 | 0.00001 | E | Female | 155 | 3.60860 | -24.23052 | 0.00000 |
| RFI1186COL_T1_LE_M | 3.608666667 | -76.918 | 0.08032 | E | Male | 155 | 3.60860 | -24.23052 | 0.08840 |
| RFI1187COL_T1_LE_M | 3.60867 | -76.918 | 0.15921 | E | Male | 163 | 3.60860 | -24.23052 | 0.21886 |
| RFI1176COL_T1_LD_F | 3.576166667 | -76.88008333 | 0.09634 | D | Female | 637 | 3.58134 | -19.05655 | 0.07528 |
| RFI1216COL_T1_LD_F | 3.576166667 | -76.88008333 | 0.17852 | D | Female | 637 | 3.58134 | -19.05655 | 0.07333 |
| RFI1177COL_T1_LD_M | 3.576166667 | -76.88008333 | 0.20912 | D | Male | 637 | 3.58134 | -19.05655 | 0.08571 |
| RFI1185COL_T1_LD_F | 3.576138889 | -76.87947222 | 0.23774 | D | Female | 608 | 3.58090 | -18.97316 | 0.23045 |
| RFI609COL_T1_LD_M | 3.575033333 | -76.87931667 | 0.00001 | D | Male | 632 | 3.58079 | -18.95193 | 0.08232 |
| RFI611COL_T1_LD_M | 3.575033333 | -76.87931667 | 0.00001 | D | Male | 632 | 3.58079 | -18.95193 | 0.14758 |
| RFI1179COL_T1_LD_M | 3.572916667 | -76.87866667 | 0.18186 | D | Male | 595 | 3.58033 | -18.86324 | 0.05362 |
| RFI1222COL_T1_LD_M | 3.572916667 | -76.87866667 | 0.19292 | D | Male | 593 | 3.58033 | -18.86324 | 0.06524 |
| RFI612COL_T1_LD_M | 3.57427 | -76.87782 | 0.00001 | D | Male | 607 | 3.57972 | -18.74770 | 0.09487 |
| RFI608COL_T1_LD_M | 3.574266667 | -76.87781667 | 0.20936 | D | Male | 601 | 3.57971 | -18.74725 | 0.22381 |
| RFI1178COL_T1_LD_F | 3.534611111 | -76.87091667 | 0.22772 | D | Female | 666 | 3.57475 | -17.80569 | 0.04944 |
| RFI610COL_T1_LD_F | 3.574583333 | -76.85469444 | 0.21722 | D | Female | 605 | 3.56309 | -15.59202 | 0.08042 |
| Elevation | NA | -76.82756043 | NA | Elevation | NA | NA | 3.54359 | -11.88930 | NA |

| | | | | | | | | | |
|---|---|---|---|---|---|---|---|---|---|
| RFI1206COL_T1_LC_F | 3.59608 | -76.79569 | 0.00001 | C | Female | 1100 | 3.52068 | -7.54018 | 0.17641 |
| RFI1208COL_T1_LC_M | 3.59528 | -76.79556 | 0.00001 | C | Male | 1100 | 3.52059 | -7.52244 | 0.18825 |
| RFI606COL_T1_LC_M | 3.84458897 | -76.7931059 | 0.17972 | C | Male | 245 | 3.51882 | -7.18754 | 0.11557 |
| RFI605COL_T1_LC_M | 3.83566358 | -76.79076542 | 0.33948 | C | Male | 289 | 3.51714 | -6.86815 | 0.24019 |
| RFI1204COL_T1_LC_M | 3.57614 | -76.75619 | 0.00001 | C | Male | 1198 | 3.49228 | -2.14982 | 0.12696 |
| RFI1202COL_T1_LC_M | 3.57614 | -76.75619 | 0.18200 | C | Male | 1198 | 3.49228 | -2.14982 | 0.09348 |
| Qisocline | NA | -76.7404366 | NA | Qisocline | NA | NA | 3.48096 | 0.00000 | NA |
| RFI1189COL_T1_LB_F | 3.525805556 | -76.73261111 | 0.22453 | B | Female | 1393 | 3.47533 | 1.06793 | 0.01524 |
| RFI1192COL_T1_LB_F | 3.525805556 | -76.73261111 | 0.22942 | B | Female | 1393 | 3.47533 | 1.06793 | 0.00000 |
| RFI1188COL_T1_LB_F | 3.525805556 | -76.73261111 | 0.30590 | B | Female | 1393 | 3.47533 | 1.06793 | 0.08524 |
| RFH1191COL_T1_LB_M | 3.525805556 | -76.73261111 | 0.66877 | B | Male | 1393 | 3.47533 | 1.06793 | 0.45454 |
| RFH1190COL_T1_LB_M | 3.525805556 | -76.73261111 | 0.99999 | B | Male | 1393 | 3.47533 | 1.06793 | 0.72937 |
| RFH1182COL_T1_LB_F | 3.526333333 | -76.71952778 | 0.99999 | B | Female | 1439 | 3.46593 | 2.85339 | 0.40531 |
| RFI1180COL_T1_LB_M | 3.527833333 | -76.71916667 | 0.22445 | B | Male | 1465 | 3.46567 | 2.90267 | 0.19529 |
| TreeLine | NA | -76.7204366 | NA | TreeLine | NA | NA | 3.46096 | 3.13174 | NA |
| RFH1181COL_T1_LB_M | 3.528138889 | -76.71488889 | 0.99999 | B | Male | 1470 | 3.46260 | 3.48645 | 0.84017 |
| RFH1183COL_T1_LB_F | 3.5705 | -76.69719444 | 0.88526 | B | Female | 1222 | 3.44988 | 5.90120 | 0.51549 |
| RFF614COL_T1_LA_F | 3.4275 | -76.64338889 | 0.99999 | A | Female | 2063 | 3.41120 | 13.24416 | 0.67535 |
| RFH1198COL_T1_LA_M | 3.455 | -76.63505556 | 0.99999 | A | Male | 1742 | 3.40521 | 14.38144 | 0.69132 |
| RFF616COL_T1_LA_F | 3.525861111 | -76.59361111 | 0.99999 | A | Female | 1847 | 3.37542 | 20.03761 | 0.83157 |
| RFF334ACOL_T1_LA_M | 3.566194444 | -76.58133333 | 0.99999 | A | Male | 2010 | 3.36659 | 21.71325 | 0.89618 |
| RFF335ACOL_T1_LA_F | 3.566194444 | -76.58133333 | 0.99999 | A | Female | 2010 | 3.36659 | 21.71325 | 0.63835 |
| RFF1197COL_T1_LA_F | 2.477305556 | -76.57544444 | 0.99999 | A | Female | 1840 | 3.36236 | 22.51695 | 1.00000 |
| RFF1196COL_T1_LA_M | 2.477305556 | -76.57544444 | 0.99999 | A | Male | 1840 | 3.36236 | 22.51695 | 1.00000 |
| RFF601COL_T1_LA_F | 3.571388889 | -76.57475 | 0.99999 | A | Female | 1825 | 3.36186 | 22.61173 | 0.82400 |
| RFF599COL_T1_LA_F | 3.575444444 | -76.57158333 | 0.78227 | A | Female | 1762 | 3.35958 | 23.04391 | 0.56236 |

Elevation, tree landcover and Q isocline indicate the intersection point of the contour lines for 1100m of elevation, 85% tree landcover and Q = 0.5 with the linear transect respectively.

**S9 Table. Genetic and morphological sampling scheme for T2.**

| Sample ID | Latitude | Longitude | Q | Locality | Sex | Elevation | Fitted Latitude (Linear Transect) | Distance (Km) | Rump color (Hue) |
|---|---|---|---|---|---|---|---|---|---|
| RFI108COL_T2_LH_F | 5.2613 | -76.5512 | 0.00001 | H | Female | 75 | 5.35041 | -50.98822 | 0.04624 |
| RFI308COL_T2_LH_M | 5.274115 | -76.5461978 | 0.00001 | H | Male | 88 | 5.34915 | -50.41717 | 0.09101 |
| RFI311COL_T2_LH_U | 5.274115 | -76.5461978 | 0.00001 | H | U | 88 | 5.34915 | -50.41717 | NA |
| RFI310COL_T2_LH_F | 5.274115 | -76.5461978 | 0.00001 | H | Female | 88 | 5.34915 | -50.41717 | 0.05560 |
| RFI309COL_T2_LH_M | 5.274115 | -76.5461978 | 0.00001 | H | Male | 88 | 5.34915 | -50.41717 | 0.06812 |
| RFI303COL_T2_LH_F | 5.25386 | -76.52311 | 0.00001 | H | Female | 88 | 5.34333 | -47.78145 | 0.00000 |
| RFI305COL_T2_LH_F | 5.25386 | -76.52311 | 0.00001 | H | Female | 88 | 5.34333 | -47.78145 | 0.08274 |
| RFI304COL_T2_LH_F | 5.25386 | -76.52311 | 0.00001 | H | Female | 88 | 5.34333 | -47.78145 | 0.03225 |
| RFI307COL_T2_LH_M | 5.25386 | -76.52311 | 0.00001 | H | Male | 88 | 5.34333 | -47.78145 | 0.06405 |
| RFI306COL_T2_LH_M | 5.25386 | -76.52311 | 0.00001 | H | Male | 88 | 5.34333 | -47.78145 | 0.11027 |
| RFI302COL_T2_LG_M | 5.29916 | -76.43832 | 0.00001 | G | Male | 91 | 5.32196 | -38.10159 | 0.07471 |
| RFI200COL_T2_LG_M | 5.29916 | -76.43832 | 0.00001 | G | Male | 91 | 5.32196 | -38.10159 | 0.00000 |
| RFI301COL_T2_LG_F | 5.29916 | -76.43832 | 0.00001 | G | Female | 91 | 5.32196 | -38.10159 | 0.01496 |
| RFI198COL_T2_LG_M | 5.29916 | -76.43832 | 0.00001 | G | Male | 91 | 5.32196 | -38.10159 | 0.04723 |
| RFI196COL_T2_LG_F | 5.29916 | -76.43832 | 0.00001 | G | Female | 91 | 5.32196 | -38.10159 | 0.05982 |
| RFI199COL_T2_LG_F | 5.29916 | -76.43832 | 0.00001 | G | Female | 91 | 5.32196 | -38.10159 | 0.01693 |
| Elevation | NA | -76.2099871 | NA | Elevation | NA | NA | 5.26442 | -12.03291 | NA |
| RFI111COL_T2_LF_M | 5.3405496 | -76.1523781 | 0.14560 | F | Male | 335 | 5.24990 | -5.45535 | 0.04820 |
| RFI192COL_T2_LF_F | 5.3405496 | -76.1523781 | 0.18675 | F | Female | 341 | 5.24990 | -5.45535 | 0.02747 |
| RFI194COL_T2_LF_M | 5.3405496 | -76.1523781 | 0.22051 | F | Male | 341 | 5.24990 | -5.45535 | 0.08073 |
| RFI190COL_T2_LF_U | 5.3405496 | -76.1523781 | 0.23779 | F | U | 341 | 5.24990 | -5.45535 | NA |
| RFI191COL_T2_LF_M | 5.3405496 | -76.1523781 | 0.24944 | F | Male | 341 | 5.24990 | -5.45535 | 0.30580 |
| RFI195COL_T2_LF_F | 5.3405496 | -76.1523781 | 0.28758 | F | Female | 341 | 5.24990 | -5.45535 | 0.08509 |
| RFI193COL_T2_LF_M | 5.3405496 | -76.1523781 | 0.29590 | F | Male | 341 | 5.24990 | -5.45535 | 0.12574 |

| | | | | | | | | | |
|---|---|---|---|---|---|---|---|---|---|
| RFI114COL_T2_LF_M | 5.3405496 | -76.1523781 | 0.31440 | F | Male | 335 | 5.24990 | -5.45535 | 0.04548 |
| RFI189COL_T2_LF_M | 5.3405496 | -76.1523781 | 0.33659 | F | Male | 341 | 5.24990 | -5.45535 | 0.18037 |
| RFI113COL_T2_LF_M | 5.3405496 | -76.1523781 | 0.33662 | F | Male | 335 | 5.24990 | -5.45535 | 0.17148 |
| RFI112COL_T2_LF_M | 5.3405496 | -76.1523781 | 0.33931 | F | Male | 335 | 5.24990 | -5.45535 | 0.04719 |
| RFI109COL_T2_LF_F | 5.3405496 | -76.1523781 | 0.42542 | F | Female | 335 | 5.24990 | -5.45535 | 0.13502 |
| RFI110COL_T2_LF_F | 5.3405496 | -76.1523781 | 0.45818 | F | Female | 335 | 5.24990 | -5.45535 | 0.12387 |
| Qisocline | NA | -76.1045988 | NA | Qisocline | NA | NA | 5.23786 | 0.00000 | NA |
| RFI178COL_T2_LE_F | 5.2499516 | -76.0509846 | 0.54345 | E | Female | 1216 | 5.22435 | 6.12168 | 0.28555 |
| RFH176COL_T2_LE_M | 5.2499516 | -76.0509846 | 0.57692 | E | Male | 1216 | 5.22435 | 6.12168 | 0.35292 |
| RFI175COL_T2_LE_M | 5.2499516 | -76.0509846 | 0.58945 | E | Male | 1216 | 5.22435 | 6.12168 | 0.25513 |
| RFH177COL_T2_LE_M | 5.2499516 | -76.0509846 | 0.60032 | E | Male | 1216 | 5.22435 | 6.12168 | 0.59056 |
| RFI184COL_T2_LE_M | 5.2519906 | -76.0507449 | 0.52106 | E | Male | 1222 | 5.22429 | 6.14905 | 0.26932 |
| RFI185COL_T2_LE_F | 5.2519906 | -76.0507449 | 0.52626 | E | Female | 1222 | 5.22429 | 6.14905 | 0.30748 |
| RFI186COL_T2_LE_M | 5.2519906 | -76.0507449 | 0.55130 | E | Male | 1222 | 5.22429 | 6.14905 | 0.31761 |
| RFI187COL_T2_LE_M | 5.2519906 | -76.0507449 | 0.58140 | E | Male | 1222 | 5.22429 | 6.14905 | 0.29742 |
| RFI188COL_T2_LE_F | 5.2519906 | -76.0507449 | 0.58504 | E | Female | 1222 | 5.22429 | 6.14905 | 0.27174 |
| RFH182COL_T2_LE_M | 5.2519906 | -76.0507449 | 0.59418 | E | Male | 1222 | 5.22429 | 6.14905 | 0.62647 |
| RFH181COL_T2_LE_M | 5.2519906 | -76.0507449 | 0.59785 | E | Male | 1222 | 5.22429 | 6.14905 | 0.82997 |
| RFH179COL_T2_LE_M | 5.2519906 | -76.0507449 | 0.65281 | E | Male | 1222 | 5.22429 | 6.14905 | 0.85490 |
| RFI183COL_T2_LE_M | 5.2519906 | -76.0507449 | 0.66276 | E | Male | 1222 | 5.22429 | 6.14905 | 0.26829 |
| RFH180COL_T2_LE_M | 5.2519906 | -76.0507449 | 0.66418 | E | Male | 1222 | 5.22429 | 6.14905 | 0.76834 |
| RFF115COL_T2_LE_F | 5.221031 | -76.024403 | 0.52839 | E | Female | 1450 | 5.21765 | 9.15682 | 0.11149 |
| RFF116COL_T2_LE_M | 5.221031 | -76.024403 | 0.73779 | E | Male | 1450 | 5.21765 | 9.15682 | 0.63236 |
| RFF173COL_T2_LD_F | 5.1057108 | -75.9390822 | 0.79063 | D | Female | 1676 | 5.19615 | 18.89913 | 0.60695 |
| RFF172COL_T2_LD_F | 5.1057108 | -75.9390822 | 0.91621 | D | Female | 1676 | 5.19615 | 18.89913 | 1.00000 |
| RFF174COL_T2_LD_M | 5.1057108 | -75.9390822 | 0.97080 | D | Male | 1676 | 5.19615 | 18.89913 | 0.80145 |
| RFF170COL_T2_LD_M | 5.1118423 | -75.9360677 | 0.94312 | D | Male | 1792 | 5.19539 | 19.24334 | 0.95625 |
| RFF171COL_T2_LD_F | 5.1111554 | -75.9327149 | 0.86181 | D | Female | 1902 | 5.19454 | 19.62619 | 0.66700 |

| | | | | | | | | | |
|---|---|---|---|---|---|---|---|---|---|
| RFF117COL_T2_LC_M | 5.268865 | -75.9009971 | 0.86375 | C | Male | 1981 | 5.18655 | 23.24795 | 0.89887 |
| RFF160COL_T2_LC_M | 5.268865 | -75.9009971 | 0.88986 | C | Male | 1981 | 5.18655 | 23.24795 | 0.93985 |
| RFH169COL_T2_LC_F | 5.2064 | -75.90034 | 0.72865 | C | Female | 1704 | 5.18638 | 23.32298 | 0.65351 |
| RFF166COL_T2_LC_M | 5.2064 | -75.90034 | 0.82911 | C | Male | 1704 | 5.18638 | 23.32298 | 0.97348 |
| RFF167COL_T2_LC_F | 5.2064 | -75.90034 | 0.89067 | C | Female | 1704 | 5.18638 | 23.32298 | 0.98902 |
| RFF168COL_T2_LC_M | 5.2064 | -75.90034 | 0.93811 | C | Male | 1704 | 5.18638 | 23.32298 | 0.94196 |
| RFF165COL_T2_LC_M | 5.2620272 | -75.8986003 | 0.93235 | C | Male | 1868 | 5.18594 | 23.52164 | 0.97095 |
| RFF162COL_T2_LC_M | 5.2620272 | -75.8986003 | 0.96947 | C | Male | 1868 | 5.18594 | 23.52164 | 0.81445 |
| RFF164COL_T2_LC_M | 5.2620272 | -75.8986003 | 0.96978 | C | Male | 1868 | 5.18594 | 23.52164 | 0.94425 |
| RFF163COL_T2_LC_F | 5.2620272 | -75.8986003 | 0.99549 | C | Female | 1868 | 5.18594 | 23.52164 | 0.80562 |
| RFF161COL_T2_LC_M | 5.2620272 | -75.8986003 | 0.99999 | C | Male | 1868 | 5.18594 | 23.52164 | 0.92601 |
| TreeLine | NA | -75.87828478 | NA | TreeLine | NA | NA | 5.18082 | 25.84143 | NA |
| RFF118COL_T2_LB_M | 5.0871 | -75.6088 | 0.99999 | B | Male | 1295 | 5.11291 | 56.61511 | 0.90978 |
| RFF149COL_T2_LB_M | 5.0572 | -75.5965 | 0.99999 | B | Male | 1225 | 5.10981 | 58.01978 | 1.00000 |
| RFF150COL_T2_LB_M | 5.0337 | -75.57085 | 0.99999 | B | Male | 1292 | 5.10335 | 60.94904 | 0.95229 |
| RFF1762COL_T2_LB_F | 5.0337 | -75.57085 | 0.99999 | B | Female | 1292 | 5.10335 | 60.94904 | 0.97078 |
| RFF146COL_T2_LB_F | 5.03726 | -75.5657 | 0.99999 | B | Female | 1304 | 5.10205 | 61.53718 | 0.95010 |
| RFF144COL_T2_LB_F | 5.03726 | -75.5657 | 0.99999 | B | Female | 1304 | 5.10205 | 61.53718 | 0.94207 |
| RFF145COL_T2_LB_M | 5.034288 | -75.56409 | 0.91972 | B | Male | 1341 | 5.10164 | 61.72104 | 0.98664 |
| RFF147COL_T2_LB_M | 5.0379 | -75.564 | 0.99999 | B | Male | 1326 | 5.10162 | 61.73132 | 0.99746 |
| RFF148COL_T2_LB_F | 5.0379 | -75.564 | 0.99999 | B | Female | 1326 | 5.10162 | 61.73132 | 0.96481 |
| RFF158COL_T2_LA_M | 5.06605 | -75.53032 | 0.99999 | A | Male | 1941 | 5.09313 | 65.57767 | 0.99430 |
| RFF159COL_T2_LA_F | 5.06605 | -75.53032 | 0.99999 | A | Female | 1941 | 5.09313 | 65.57767 | 0.96881 |
| RFF155COL_T2_LA_M | 5.06605 | -75.53032 | 0.99999 | A | Male | 1941 | 5.09313 | 65.57767 | 0.98358 |
| RFF156COL_T2_LA_F | 5.06605 | -75.53032 | 0.99999 | A | Female | 1941 | 5.09313 | 65.57767 | 0.94308 |
| RFF157COL_T2_LA_F | 5.06605 | -75.53032 | 0.99999 | A | Female | 1941 | 5.09313 | 65.57767 | 0.96607 |
| RFF154COL_T2_LA_M | 5.0666 | -75.52963 | 0.99999 | A | Male | 1957 | 5.09296 | 65.65647 | 0.95024 |
| RFF152COL_T2_LA_F | 5.0679 | -75.5285 | 0.99999 | A | Female | 1987 | 5.09268 | 65.78552 | 0.98407 |

| | | | | | | | | | |
|---|---|---|---|---|---|---|---|---|---|
| RFF153COL_T2_LA_F | 5.0679 | -75.5285 | 0.99999 | A | Female | 1987 | 5.09268 | 65.78552 | 0.93663 |
| RFF151COL_T2_LA_M | 5.0679 | -75.5285 | 0.99999 | A | Male | 1987 | 5.09268 | 65.78552 | 0.95543 |

Elevation, tree landcover and Q isocline indicate the intersection point of the contour lines for 1100m of elevation, 85% tree landcover and Q = 0.5 with the linear transect respectively.

**S10 Table. Genetic and morphological sampling scheme for T3.**

| Sample ID | Latitude | Longitude | Q | Locality | Sex | Elevation | Fitted Latitude (Linear Transect) | Distance (Km) | Rump color (Hue) |
|---|---|---|---|---|---|---|---|---|---|
| RFI107COL_T3_LH_M | 5.7425991 | -76.5355638 | 0.00001 | H | Male | 46 | 5.68098 | -35.7808 | 0.0291 |
| RFI106COL_T3_LH_M | 5.7425991 | -76.5355638 | 0.00001 | H | Male | 46 | 5.68098 | -35.7808 | 0 |
| RFI318COL_T3_LH_U | 5.7425991 | -76.5355638 | 0.00001 | H | U | 45.7178 | 5.68098 | -35.7808 | 0.0509 |
| RFI314COL_T3_LH_M | 5.7455185 | -76.5339415 | 0.00001 | H | Male | 91.4355 | 5.6816 | -35.5886 | 0.1138 |
| RFI316COL_T3_LH_F | 5.7455185 | -76.5339415 | 0.00001 | H | Female | 91.4355 | 5.6816 | -35.5886 | 0.0414 |
| RFI315COL_T3_LH_F | 5.7455185 | -76.5339415 | 0.37069 | H | Female | 91.4355 | 5.6816 | -35.5886 | 0.2538 |
| RFI312COL_T3_LH_M | 5.7459795 | -76.5320093 | 0.00001 | H | Male | 76.1963 | 5.68235 | -35.3596 | 0.1067 |
| RFH324COL_T3_LG_M | 5.7209532 | -76.3115756 | 0.00001 | G | Male | 451.082 | 5.76688 | -9.24336 | 0.1568 |
| RFH325COL_T3_LG_F | 5.7209532 | -76.3115756 | 0.00001 | G | Female | 451.082 | 5.76688 | -9.24336 | 0.0768 |
| RFH321COL_T3_LG_U | 5.7209532 | -76.3115756 | 0.00296 | G | U | 451.082 | 5.76688 | -9.24336 | NA |
| RFH323COL_T3_LG_M | 5.7209532 | -76.3115756 | 0.32502 | G | Male | 451.082 | 5.76688 | -9.24336 | 0.4885 |
| RFH320COL_T3_LG_M | 5.7209532 | -76.3115756 | 0.32986 | G | Male | 451.082 | 5.76688 | -9.24336 | 0.4272 |
| RFI352COL_T3_LG_M | 5.725073 | -76.3111552 | 0.09431 | G | Male | 448.034 | 5.76704 | -9.19356 | 0.0987 |
| RFH351COL_T3_LG_F | 5.725073 | -76.3111552 | 0.64764 | G | Female | 448.034 | 5.76704 | -9.19356 | 0.8459 |
| RFI355COL_T3_LF_M | 5.75282 | -76.25601 | 0.15261 | F | Male | 716.245 | 5.78819 | -2.66066 | 0.2546 |
| RFI354COL_T3_LF_F | 5.75282 | -76.25601 | 0.26163 | F | Female | 716.245 | 5.78819 | -2.66066 | 0.195 |
| RFI353COL_T3_LF_M | 5.75282 | -76.25601 | 0.30168 | F | Male | 716.245 | 5.78819 | -2.66066 | 0.2225 |
| RFH356COL_T3_LF_U | 5.75282 | -76.25601 | 0.38419 | F | U | 716.245 | 5.78819 | -2.66066 | 0.1429 |
| Elevation | NA | -76.24543163 | NA | Elevation | NA | NA | 5.79225 | -1.40749 | NA |
| RFI348COL_T3_LF_F | 5.774774 | -76.2442827 | 0.19193 | F | Female | 761.963 | 5.79269 | -1.27139 | 0 |
| RFH350COL_T3_LF_F | 5.774774 | -76.2442827 | 0.22735 | F | Female | 761.963 | 5.79269 | -1.27139 | 0.2607 |
| RFI359COL_T3_LF_M | 5.774774 | -76.2442827 | 0.33650 | F | Male | 761.963 | 5.79269 | -1.27139 | 0.1106 |
| RFH349COL_T3_LF_F | 5.774774 | -76.2442827 | 0.37365 | F | Female | 761.963 | 5.79269 | -1.27139 | 0.794 |
| RFH357COL_T3_LF_M | 5.774774 | -76.2442827 | 0.47364 | F | Male | 761.963 | 5.79269 | -1.27139 | 0.6887 |

| | | | | | | | | | |
|---|---|---|---|---|---|---|---|---|---|
| RFH358COL_T3_LF_F | 5.774774 | -76.2442827 | 0.48026 | F | Female | 761.963 | 5.79269 | -1.27139 | 0.244 |
| Qisocline | NA | -76.2335504 | NA | Qisocline | NA | NA | 5.79681 | 0 | NA |
| RFH104COL_T3_LE_M | 5.8544 | -76.1855 | 0.71466 | E | Male | 1466.02 | 5.81523 | 5.692155 | 0.903 |
| RFH105COL_T3_LE_F | 5.8544 | -76.1855 | 0.90234 | E | Female | 1466.02 | 5.81523 | 5.692155 | 0.5579 |
| RFH331COL_T3_LE_U/F | 5.8544 | -76.1855 | 0.93799 | E | U/Female | 1466.02 | 5.81523 | 5.692155 | 0.8299 |
| RFH332COL_T3_LE_U/F | 5.8544 | -76.1855 | 0.95566 | E | U/Female | 1466.02 | 5.81523 | 5.692155 | 0.8383 |
| RFH327COL_T3_LE_M | 5.8520953 | -76.1824061 | 0.97940 | E | Male | 1462.97 | 5.81642 | 6.05866 | 0.7414 |
| RFH333COL_T3_LE_F | 5.8520953 | -76.1824061 | 0.99999 | E | Female | 1462.97 | 5.81642 | 6.05866 | 0.9198 |
| RFH334COL_T3_LE_M | 5.8520953 | -76.1824061 | 0.99999 | E | Male | 1462.97 | 5.81642 | 6.05866 | 0.7525 |
| RFH328COL_T3_LE_M | 5.8520953 | -76.1824061 | 0.99999 | E | Male | 1462.97 | 5.81642 | 6.05866 | 0.9473 |
| RFH329COL_T3_LE_M | 5.8520953 | -76.1824061 | 0.99999 | E | Male | 1462.97 | 5.81642 | 6.05866 | 0.9262 |
| RFF343COL_T3_LD_F | 5.80205 | -76.02899 | 0.99999 | D | Female | 1523.93 | 5.87526 | 24.23153 | 0.8502 |
| RFF126COL_T3_LD_M | 5.7995 | -76.0288 | 0.96221 | D | Male | 1438.59 | 5.87533 | 24.25404 | 0.9869 |
| RFF125COL_T3_LD_F | 5.7995 | -76.0288 | 0.99999 | D | Female | 1438.59 | 5.87533 | 24.25404 | 0.8106 |
| RFF342COL_T3_LD_M | 5.7973209 | -76.0283833 | 0.99999 | D | Male | 1426.39 | 5.87549 | 24.3034 | 0.9046 |
| RFF338COL_T3_LD_M | 5.7998596 | -76.0279994 | 0.99999 | D | Male | 1371.53 | 5.87564 | 24.34887 | 0.9637 |
| RFF340COL_T3_LD_F | 5.7997512 | -76.021309 | 0.97804 | D | Female | 1350.2 | 5.8782 | 25.14134 | 1 |
| RFF339COL_T3_LD_F | 5.7997512 | -76.021309 | 0.99999 | D | Female | 1350.2 | 5.8782 | 25.14134 | 0.9127 |
| RFF341COL_T3_LD_M | 5.7997512 | -76.021309 | 0.99999 | D | Male | 1350.2 | 5.8782 | 25.14134 | 0.9041 |
| RFF335COL_T3_LD_M | 5.79653 | -76.013549 | 0.99999 | D | Male | 1386.77 | 5.88118 | 26.0605 | 0.8469 |
| RFF336COL_T3_LD_M | 5.79653 | -76.013549 | 0.99999 | D | Male | 1386.77 | 5.88118 | 26.0605 | 0.9432 |
| RFF337COL_T3_LD_M | 5.79653 | -76.013549 | 0.99999 | D | Male | 1386.77 | 5.88118 | 26.0605 | 0.9765 |
| TreeLine | NA | -75.95007345 | NA | TreeLine | NA | NA | 5.90552 | 33.57893 | NA |
| RFF143COL_T3_LC_M | 5.9554 | -75.733 | 0.99999 | C | Male | 1408.11 | 5.98877 | 59.28827 | 1 |
| RFF137COL_T3_LC_F | 6.09554 | -75.71358 | 0.99999 | C | Female | 1789.09 | 5.99622 | 61.58813 | 0.9683 |
| RFF136COL_T3_LC_M | 6.09554 | -75.71358 | 0.99999 | C | Male | 1789.09 | 5.99622 | 61.58813 | 0.904 |
| RFF139COL_T3_LC_M | 6.1006 | -75.7101 | 0.99999 | C | Male | 1779.95 | 5.99755 | 62.00026 | 0.8728 |
| RFF138COL_T3_LC_M | 6.1006 | -75.7101 | 0.99999 | C | Male | 1779.95 | 5.99755 | 62.00026 | 0.896 |

| | | | | | | | | | |
|---|---|---|---|---|---|---|---|---|---|
| RFF140COL_T3_LC_F | 6.1006 | -75.7101 | 0.99999 | C | Female | 1779.95 | 5.99755 | 62.00026 | 0.8342 |
| RFF141COL_T3_LC_U | 6.1051 | -75.6994 | 0.99999 | C | U | 1895.76 | 6.00166 | 63.26742 | 0.9013 |
| RFF142COL_T3_LC_F | 6.1051 | -75.6994 | 0.99999 | C | Female | 1895.76 | 6.00166 | 63.26742 | 0.7692 |
| RFF124COL_T3_LC_M | 5.968364 | -75.677051 | 0.99999 | C | Male | 1237 | 6.01023 | 65.9141 | 0.9001 |
| RFFA42COL_T3_LC_U | 5.968364 | -75.677051 | 0.99999 | C | U | 1237 | 6.01023 | 65.9141 | NA |
| RFF134COL_T3_LB_M | 6.09898 | -75.61516 | 0.99999 | B | Male | 1816.52 | 6.03396 | 73.24336 | 0.9584 |
| RFF135COL_T3_LB_F | 6.09898 | -75.61516 | 0.99999 | B | Female | 1816.52 | 6.03396 | 73.24336 | 0.9525 |
| RFF131COL_T3_LB.1_F | 6.133493 | -75.604142 | 0.67267 | B.1 | Female | 1749.47 | 6.03819 | 74.54811 | 0.7711 |
| RFF132COL_T3_LB.1_M | 6.142165 | -75.57372 | 0.97464 | B.1 | Male | 1831.45 | 6.04986 | 78.15062 | 0.9676 |
| RFF127COL_T3_LB.1_M | 6.136891 | -75.569427 | 0.86426 | B.1 | Male | 1871.38 | 6.0515 | 78.65898 | 0.5038 |
| RFF129COL_T3_LB.1_M | 6.136891 | -75.569427 | 0.94159 | B.1 | Male | 1871.38 | 6.0515 | 78.65898 | 0.8548 |
| RFF130COL_T3_LB_F | 6.136891 | -75.569427 | 0.98209 | B | Female | 1871.38 | 6.0515 | 78.65898 | 0.8638 |
| RFF128COL_T3_LB_F | 6.136891 | -75.569427 | 0.99872 | B | Female | 1871.38 | 6.0515 | 78.65898 | 0.7365 |
| RFF347COL_T3_LA.1_F | 5.9750942 | -75.4607568 | 0.76209 | A.1 | Female | 2371.23 | 6.09318 | 91.5269 | 0.778 |
| RFF346COL_T3_LA_M | 5.9750942 | -75.4607568 | 0.99999 | A | Male | 2371.23 | 6.09318 | 91.5269 | 0.9589 |
| RFF345COL_T3_LA_F | 5.9750942 | -75.4607568 | 0.99999 | A | Female | 2371.23 | 6.09318 | 91.5269 | 0.9841 |
| RFF344COL_T3_LA_M | 5.9750942 | -75.4607568 | 0.99999 | A | Male | 2371.23 | 6.09318 | 91.5269 | 0.9213 |
| RFF133COL_T3_LA.1_F | 6.130826 | -75.4300994 | 0.84979 | A.1 | Female | 2109.11 | 6.10494 | 95.15696 | 0.6291 |
| RFF101COL_T3_LA.1_M | 6.056436 | -75.421677 | 0.96080 | A.1 | Male | 2148 | 6.10817 | 96.15423 | 0.5739 |
| RFF119COL_T3_LA.1_F | 6.05675 | -75.42129 | 0.80703 | A.1 | Female | 2173 | 6.10831 | 96.20005 | 0.6577 |
| RFF120COL_T3_LA.1_M | 6.05675 | -75.42129 | 0.89373 | A.1 | Male | 2173 | 6.10831 | 96.20005 | 0.3633 |

Elevation, tree landcover and Q isocline indicate the intersection point of the contour lines for 1100m of elevation, 85% tree landcover and Q = 0.5 with the linear transect respectively.

**S11 Table. Data used for geographic cline analysis of genetic markers.**

| Transect | Sex | Locality | Distance | Q | SNP 1 | SNP 2 |
|---|---|---|---|---|---|---|
| 1 | Male | A | 20.19769289 | 0.99999 (N=3) | 1 (N=6) | 1 (N=6) |
| | | B | 2.275925871 | 0.7233 (N=4) | 0.83333 (N=6) | 0.75 (N=8) |
| | | C | -5.569658769 | 0.14024 (N=5) | 0.2 (N=10) | 0 (N=10) |
| | | D | -18.5554833 | 0.11333 (N=7) | 0.21429 (N=14) | 0 (N=14) |
| | | E | -24.23052295 | 0.11977 (N=2) | 0 (N=2) | 0 (N=2) |
| | | F | -35.38549021 | 0.00001 (N=1) | 0 (N=2) | 0 (N=2) |
| | | G | -86.06966022 | 0.02434 (N=5) | 0 (N=10) | 0 (N=10) |
| | Female | A | 20.19769289 | 0.9637 (N=6) | 1 (N=8) | 1 (N=6) |
| | | B | 2.275925871 | 0.52902 (N=5) | 0.625 (N=8) | 0.75 (N=8) |
| | | C | -5.569658769 | 0.00001 (N=2) | 0 (N=2) | 0 (N=2) |
| | | D | -18.5554833 | 0.19151 (N=5) | 0.2 (N=10) | 0.25 (N=8) |
| | | E | -24.23052295 | 0.00001 (N=1) | 0.5 (N=2) | 0 (N=2) |
| | | F | -35.38549021 | 0.00001 (N=2) | 0 (N=4) | 0 (N=4) |
| 2 | Male | A | 65.65570702 | 0.99999 (N=4) | 1 (N=8) | 1 (N=8) |
| | | B | 60.5323325 | 0.98394 (N=5) | 1 (N=10) | 1 (N=10) |
| | | C | 23.39963877 | 0.92405 (N=8) | 0.9375 (N=16) | 0.8125 (N=16) |
| | | D | 19.11338225 | 0.95696 (N=2) | 0.75 (N=4) | 1 (N=4) |
| | | E | 6.518182551 | 0.61083 (N=12) | 0.58333 (N=24) | 0.58333 (N=24) |
| | | F | -5.455349585 | 0.2798 (N=8) | 0.21429 (N=14) | 0.125 (N=16) |
| | | G | -38.10158934 | 0.00001 (N=3) | 0.5 (N=2) | 0 (N=2) |
| | | H | -49.15641635 | 0.00001 (N=4) | 0 (N=4) | 0 (N=6) |
| | Female | A | 65.65570702 | 0.99999 (N=5) | 1 (N=10) | 1 (N=8) |
| | | B | 60.5323325 | 0.99999 (N=4) | 1 (N=8) | 1 (N=6) |
| | | C | 23.39963877 | 0.8716 (N=3) | 0.83333 (N=6) | 0.83333 (N=6) |
| | | D | 19.11338225 | 0.85621 (N=3) | 1 (N=6) | 0.75 (N=4) |
| | | E | 6.518182551 | 0.54579 (N=4) | 0.75 (N=8) | 0 (N=8) |
| | | F | -5.455349585 | 0.31914 (N=5) | 0.2 (N=10) | 0.25 (N=8) |
| | | G | -38.10158934 | 0.00001 (N=3) | 0 (N=2) | 0 (N=4) |
| | | H | -49.15641635 | 0.00001 (N=6) | 0 (N=10) | 0 (N=4) |
| 3 | Male | A | 91.52690053 | 0.99999 (N=2) | 1 (N=4) | 1 (N=4) |
| | | B | 75.95117391 | 0.99999 (N=1) | 1 (N=4) | 1 (N=4) |
| | | C | 62.68283611 | 0.99999 (N=5) | 1 (N=8) | 0.8 (N=10) |
| | | D | 24.999765 | 0.99459 (N=7) | 0.83333 (N=6) | 1 (N=6) |
| | | E | 5.895768738 | 0.9388 (N=5) | 0.875 (N=8) | 0.75 (N=8) |
| | | F | -1.827096218 | 0.31611 (N=4) | 0.5 (N=4) | 1 (N=2) |
| | | G | -9.229132937 | 0.1873 (N=4) | 0.5 (N=6) | 0.125 (N=8) |
| | | H | -35.63823745 | 0.00001 (N=4) | 0.16667 (N=6) | 0 (N=6) |

| | | | | | | |
|---|---|---|---|---|---|---|
| | Female | A | 91.52690053 | 0.99999 (N=1) | 1 (N=2) | 1 (N=2) |
| | | B | 75.95117391 | 0.9936 (N=3) | 0.83333 (N=6) | 1 (N=6) |
| | | C | 62.68283611 | 0.99999 (N=5) | 1 (N=4) | 1 (N=6) |
| | | D | 24.999765 | 0.9945 (N=4) | 1 (N=4) | 1 (N=6) |
| | | E | 5.895768738 | 0.94899 (N=4) | 1 (N=6) | 0 (N=2) |
| | | F | -1.827096218 | 0.31984 (N=6) | 0.5 (N=2) | 1 (N=2) |
| | | G | -9.229132937 | 0.21687 (N=3) | 0.5 (N=4) | 0.5 (N=4) |
| | | H | -35.63823745 | 0.12357 (N=3) | 0.25 (N=4) | 0 (N=4) |

Average distance from the center of the hybrid zone, hybrid index (Q) and allele frequencies of the only two fixed SNPs for males and females in each locality/transect. N = Sample size.

**S12 Table. Parameters for geographic cline models tested with HZAR.**

| Model # | Scaling | Tails | Traits |
|---|---|---|---|
| Null | NA | NA | Q, Fixed SNPs, Rump Color |
| 1 | none | none | Q, Fixed SNPs |
| 2 | none | right | Q, Fixed SNPs |
| 3 | none | left | Q, Fixed SNPs |
| 4 | none | mirror | Q, Fixed SNPs |
| 5 | none | both | Q, Fixed SNPs |
| 6 | fixed | none | Q, Fixed SNPs |
| 7 | fixed | right | Q, Fixed SNPs |
| 8 | fixed | left | Q, Fixed SNPs |
| 9 | fixed | mirror | Q, Fixed SNPs |
| 10 | fixed | both | Q, Fixed SNPs |
| 11 | free | none | Q, Fixed SNPs, Rump Color |
| 12 | free | right | Q, Fixed SNPs, Rump Color |
| 13 | free | left | Q, Fixed SNPs, Rump Color |
| 14 | free | mirror | Q, Fixed SNPs, Rump Color |
| 15 | free | both | Q, Fixed SNPs, Rump Color |

**S13 Table. Center (*u*) and steepness (*v*) genomic cline values for outlier loci in T1 and T2 located within 100kb of each other.**

| Outlier | Chr | Locus T1 | Prop. *ict* T1 | Prop. *fla* T1 | Steepness (v) T1 | Center (u) T1 | AIC diff T1 | Locus T2 | Prop. *ict* T2 | Prop. *fla* T2 | Steepness (v) T2 | Center (u) T2 | AIC diff T2 |
|---|---|---|---|---|---|---|---|---|---|---|---|---|---|
| Center | 11 | 19187872 | 0.08 | 0.63 | 1.12 | 0.01 | -2.46 | 19195397 | 0.12 | 0.91 | 0.38 | 0.04 | -6.25 |
| Center | 13 | 14405123 | 0.06 | 0.64 | 10.20 | 0.69 | -1.29 | 14440397 | 0.05 | 0.56 | 0.63 | 0.68 | -2.69 |
| Center | 14 | 4762891 | 0.10 | 0.67 | 5.89 | 0.98 | -0.16 | 4762891 | 0.00 | 0.53 | 0.64 | 0.66 | -0.32 |
| Center | 14 | 16209346 | 0.05 | 0.63 | 0.37 | 1.00 | -2.24 | 16209346 | 0.07 | 0.61 | 2.80 | 0.61 | -1.73 |
| Center | 17 | 9952546 | 0.04 | 0.83 | 2.10 | 0.88 | -3.80 | 9952546 | 0.10 | 0.62 | 0.15 | 0.01 | -0.59 |
| Center | 17 | 13047816 | 0.14 | 0.92 | 0.11 | 0.99 | -0.81 | 13035911 | 0.00 | 0.56 | 1.73 | 0.80 | -15.32 |
| Center | 1A | 70343379 | 0.04 | 0.89 | 1.15 | 0.90 | -2.38 | 70343379 | 0.08 | 0.79 | 13.88 | 0.38 | -2.88 |
| Center | 1A | 70343393 | 0.04 | 0.89 | 1.26 | 0.79 | -1.14 | 70343379 | 0.08 | 0.79 | 13.88 | 0.38 | -2.88 |
| Center | 1A | 70343395 | 0.04 | 0.89 | 1.38 | 0.80 | -1.01 | 70343379 | 0.08 | 0.79 | 13.88 | 0.38 | -2.88 |
| Center | 1A | 40963616 | 0.05 | 0.57 | 0.33 | 0.01 | -0.39 | 40963618 | 0.00 | 0.50 | 0.32 | 0.95 | -4.29 |
| Center | 2 | 2659193 | 0.21 | 1.00 | 0.14 | 0.03 | -0.78 | 2659193 | 0.33 | 0.94 | 0.73 | 0.21 | -0.34 |
| Center | 2 | 130102130 | 0.00 | 0.75 | 1.05 | 0.87 | -1.22 | 130102130 | 0.03 | 0.56 | 2.85 | 0.63 | -2.40 |
| Center | 20 | 7643333 | 0.38 | 0.94 | 1.05 | 0.18 | -1.31 | 7678500 | 0.19 | 0.97 | 0.27 | 0.86 | -0.65 |
| Center | 20 | 7643333 | 0.38 | 0.94 | 1.05 | 0.18 | -1.31 | 7678502 | 0.42 | 0.97 | 0.23 | 0.95 | -1.11 |
| **Center** | **22** | **300695** | **0.39** | **0.95** | **7.92** | **0.78** | **-1.96** | **271593** | **0.23** | **0.92** | **1.64** | **0.64** | **-3.24** |
| Center | 3 | 90288754 | 0.41 | 0.94 | 4.86 | 0.34 | -1.16 | 90288754 | 0.15 | 0.96 | 0.29 | 0.03 | -1.55 |
| Center | 3 | 90288754 | 0.41 | 0.94 | 4.86 | 0.34 | -1.16 | 90358686 | 0.00 | 0.69 | 0.24 | 0.86 | -0.23 |
| Center | 4 | 11749308 | 0.05 | 0.89 | 0.69 | 0.97 | -0.07 | 11817000 | 0.06 | 0.69 | 1.69 | 0.75 | -11.50 |
| Center | 7 | 16053933 | 0.50 | 1.00 | 0.21 | 0.01 | -2.51 | 16041256 | 0.00 | 0.56 | 0.65 | 0.75 | -2.20 |
| Center | 7 | 35089368 | 0.36 | 0.95 | 7.21 | 0.92 | -7.35 | 35089368 | 0.20 | 0.91 | 0.32 | 0.00 | -7.19 |
| Center | 7 | 35089368 | 0.36 | 0.95 | 7.21 | 0.92 | -7.35 | 35089383 | 0.10 | 0.69 | 0.53 | 0.02 | -0.94 |
| Center | 8 | 2416298 | 0.44 | 1.00 | 0.26 | 0.01 | -3.93 | 2416299 | 0.21 | 0.93 | 12.51 | 0.57 | -3.67 |
| Center | 9 | 10116669 | 0.38 | 0.94 | 0.39 | 0.01 | -2.32 | 10049831 | 0.33 | 0.92 | 1.06 | 0.08 | -6.97 |
| Center | Z | 62897794 | 0.00 | 0.89 | 98.02 | 0.29 | -16.70 | 62897794 | 0.00 | 1.00 | 1.24 | 0.62 | -3.98 |
| Steepness | 1 | 107039279 | 0.07 | 0.60 | 0.32 | 1.00 | -3.14 | 107102503 | 0.07 | 0.89 | 2.11 | 0.44 | -3.54 |

| | | | | | | | | | | | | | |
|---|---|---|---|---|---|---|---|---|---|---|---|---|---|
| Steepness | 10 | 311437 | 0.00 | 0.50 | 0.12 | 0.07 | -0.13 | 254924 | 0.00 | 0.58 | 6.08 | 0.30 | -3.41 |
| Steepness | 10 | 254924 | 0.10 | 0.67 | 0.08 | 0.08 | -1.01 | 254924 | 0.00 | 0.58 | 6.08 | 0.30 | -3.41 |
| Steepness | 14 | 16209346 | 0.05 | 0.63 | 0.37 | 1.00 | -0.62 | 16209346 | 0.07 | 0.61 | 2.80 | 0.61 | -0.46 |
| Steepness | 17 | 12841286 | 0.00 | 0.64 | 0.03 | 0.68 | -0.71 | 12841286 | 0.10 | 0.74 | 3.80 | 0.69 | -0.94 |
| Steepness | 17 | 13047816 | 0.14 | 0.92 | 0.11 | 0.99 | -3.01 | 13035911 | 0.00 | 0.56 | 1.73 | 0.80 | -0.10 |
| Steepness | 1A | 17741736 | 0.29 | 0.90 | 13.92 | 0.51 | -2.51 | 17710316 | 0.07 | 0.64 | 0.05 | 0.34 | -1.03 |
| Steepness | 1A | 54230444 | 0.00 | 0.56 | 3.08 | 0.37 | -1.95 | 54152411 | 0.08 | 0.69 | 2.92 | 0.52 | -2.42 |
| Steepness | 1A | 17710326 | 0.42 | 0.94 | 8.72 | 0.54 | -1.12 | 17710316 | 0.07 | 0.64 | 0.05 | 0.34 | -1.03 |
| Steepness | 2 | 123362204 | 0.00 | 0.72 | 1.94 | 0.41 | -0.99 | 123362204 | 0.00 | 0.72 | 1.65 | 0.42 | -0.36 |
| Steepness | 2 | 88132886 | 0.08 | 0.90 | 7.20 | 0.71 | -1.84 | 88132886 | 0.07 | 0.72 | 11.05 | 0.50 | -5.99 |
| Steepness | 2 | 162910653 | 0.09 | 0.78 | 9.45 | 0.59 | -0.18 | 162816357 | 0.37 | 0.96 | 2.38 | 0.66 | -0.83 |
| **Steepness** | **22** | **300695** | **0.39** | **0.95** | **7.92** | **0.78** | **-0.11** | **271593** | **0.23** | **0.92** | **1.64** | **0.64** | **-0.39** |
| Steepness | 3 | 121591284 | 0.00 | 0.83 | 0.06 | 0.94 | -0.49 | 121551164 | 0.08 | 0.81 | 1.70 | 0.52 | -0.10 |
| Steepness | 5 | 27116078 | 0.08 | 0.75 | 14.18 | 0.59 | -3.20 | 27116078 | 0.00 | 0.58 | 1.60 | 0.43 | -0.08 |
| Steepness | 7 | 35745376 | 0.33 | 1.00 | 14.53 | 0.71 | -5.65 | 35745373 | 0.04 | 0.78 | 6.85 | 0.50 | -4.64 |
| Steepness | 7 | 35745376 | 0.33 | 1.00 | 14.53 | 0.71 | -5.65 | 35745376 | 0.08 | 0.92 | 2.05 | 0.49 | -3.13 |

Proportions were calculated based on the number of individuals that had the reference or alternative allele at each loci (only loci with allele frequency difference > 0.5 between pure parental individuals were included). AIC difference represent the extent to which a steeper-than-expected or displaced cline provided a better fit than the null model with values fixed at $u = 0.5$ and $v = 1$.

**S14 Table. Center (*u*) and steepness (*v*) genomic cline values for outlier loci in T1 and T3 located within 100kb of each other.**

| **Outlier** | **Chr** | **Locus T1** | **Prop. *ict* T1** | **Prop. *fla* T1** | **Steepness (v) T1** | **Center (u) T1** | **AIC diff T1** | **Locus T3** | **Prop. *ict* T3** | **Prop. *fla* T3** | **Steepness (v) T3** | **Center (u) T3** | **AIC diff T3** |
|---|---|---|---|---|---|---|---|---|---|---|---|---|---|
| Center | 10 | 16796254 | 0.00 | 0.50 | 7.26 | 0.31 | -4.03 | 16763558 | 0.10 | 0.70 | 5.15 | 0.02 | -7.16 |
| Center | 10 | 16796270 | 0.00 | 0.50 | 6.23 | 0.34 | -1.92 | 16763558 | 0.10 | 0.70 | 5.15 | 0.02 | -7.16 |
| Center | 17 | 13857156 | 0.50 | 1.00 | 1.20 | 0.18 | -4.08 | 13857151 | 0.33 | 0.96 | 2.03 | 0.03 | -1.37 |
| **Center** | **22** | **300695** | **0.39** | **0.95** | **7.92** | **0.78** | **-1.96** | **266539** | **0.33** | **1.00** | **0.45** | **0.36** | **-8.81** |
| Center | 3 | 39270509 | 0.08 | 0.63 | 1.13 | 0.32 | -0.26 | 39270509 | 0.00 | 0.55 | 3.92 | 0.01 | -3.56 |
| Steepness | 11 | 19195397 | 0.08 | 0.89 | 0.08 | 0.96 | -3.25 | 19195397 | 0.19 | 0.98 | 0.08 | 0.09 | -1.82 |
| Steepness | 21 | 11302387 | 0.08 | 0.94 | 3.04 | 0.52 | -0.58 | 11261387 | 0.36 | 0.95 | 0.07 | 0.49 | -0.73 |
| Steepness | 8 | 8568476 | 0.38 | 1.00 | 2.63 | 0.26 | -1.46 | 8649595 | 0.17 | 0.91 | 6.28 | 0.51 | -3.92 |

Proportions were calculated based on the number of individuals that had the reference or alternative allele at each loci (only loci with allele frequency difference > 0.5 between pure parental individuals were included). AIC difference represent the extent to which a steeper-than-expected or displaced cline provided a better fit than the null model with values fixed at $u = 0.5$ and $v = 1$.

**S15 Table. Center (*u*) and steepness (*v*) genomic cline values for outlier loci in T2 and T3 located within 100kb of each other.**

| Outlier | Chr | Locus T2 | Prop. *ict* T2 | Prop. *fla* T2 | Steepness (v) T2 | Center (u) T2 | AIC diff T2 | Locus T3 | Prop. *ict* T3 | Prop. *fla* T3 | Steepness (v) T3 | Center (u) T3 | AIC diff T3 |
|---|---|---|---|---|---|---|---|---|---|---|---|---|---|
| Center | 1 | 27925132 | 0.04 | 0.97 | 1.49 | 0.59 | -1.64 | 27925132 | 0.00 | 0.89 | 0.54 | 0.69 | -0.23 |
| Center | 10 | 18436596 | 0.30 | 0.93 | 1.06 | 0.81 | -7.93 | 18376007 | 0.30 | 1.00 | 0.79 | 0.02 | -8.19 |
| Center | 10 | 18436579 | 0.30 | 0.93 | 0.99 | 0.81 | -7.79 | 18376007 | 0.30 | 1.00 | 0.79 | 0.02 | -8.19 |
| Center | 13 | 18132472 | 0.05 | 0.56 | 1.08 | 0.83 | -5.34 | 18132472 | 0.00 | 0.53 | 17.76 | 0.63 | -4.63 |
| Center | 14 | 3018141 | 0.05 | 0.75 | 0.34 | 0.97 | -10.21 | 3018141 | 0.00 | 0.57 | 0.82 | 0.92 | -7.33 |
| Center | 14 | 3018141 | 0.05 | 0.75 | 0.34 | 0.97 | -10.21 | 3018148 | 0.00 | 0.57 | 0.94 | 0.91 | -7.72 |
| Center | 14 | 3018148 | 0.05 | 0.75 | 0.35 | 0.97 | -10.24 | 3018141 | 0.00 | 0.57 | 0.82 | 0.92 | -7.33 |
| Center | 14 | 3018148 | 0.05 | 0.75 | 0.35 | 0.97 | -10.24 | 3018148 | 0.00 | 0.57 | 0.94 | 0.91 | -7.72 |
| **Center** | **22** | **271593** | **0.23** | **0.92** | **1.64** | **0.64** | **-3.24** | **266539** | **0.33** | **1.00** | **0.45** | **0.36** | **-8.81** |
| Center | 23 | 6628152 | 0.21 | 1.00 | 0.75 | 0.71 | -3.20 | 6607300 | 0.20 | 1.00 | 1.97 | 0.79 | -4.81 |
| Center | 24 | 17870019 | 0.42 | 0.97 | 0.31 | 0.00 | -8.10 | 17964817 | 0.08 | 0.68 | 0.46 | 0.81 | -0.49 |
| Center | 3 | 6878516 | 0.21 | 0.97 | 0.83 | 0.26 | -1.33 | 6829351 | 0.25 | 0.98 | 1.23 | 0.01 | -2.73 |
| Center | 3 | 77926646 | 0.46 | 0.97 | 2.86 | 0.77 | -13.46 | 77926646 | 0.42 | 0.95 | 2.01 | 0.92 | -5.70 |
| Center | 5 | 18490823 | 0.04 | 0.65 | 0.51 | 0.06 | -0.62 | 18490823 | 0.07 | 0.61 | 12.89 | 0.58 | -0.85 |
| Center | 5 | 52987028 | 0.00 | 0.70 | 0.56 | 0.92 | -10.16 | 52987029 | 0.38 | 0.90 | 0.30 | 0.97 | -1.18 |
| Center | 5 | 64366662 | 0.07 | 0.59 | 1.88 | 0.60 | -0.32 | 64366662 | 0.00 | 0.63 | 0.50 | 0.92 | -6.22 |
| Center | 5 | 65509819 | 0.04 | 0.74 | 1.14 | 0.61 | -0.61 | 65509819 | 0.07 | 0.63 | 0.50 | 0.76 | -0.02 |
| Center | 9 | 12542763 | 0.08 | 0.78 | 3.51 | 0.41 | -0.27 | 12636779 | 0.08 | 0.73 | 1.92 | 0.74 | -0.25 |
| Center | Z | 20430134 | 0.25 | 1.00 | 3.76 | 0.67 | -16.04 | 20430134 | 0.21 | 1.00 | 0.72 | 0.02 | -3.05 |
| Center | Z | 32957938 | 0.25 | 1.00 | 0.90 | 0.12 | -10.44 | 32957938 | 0.20 | 1.00 | 0.43 | 0.03 | -4.57 |
| Steepness | 20 | 462336 | 0.08 | 0.67 | 20.53 | 0.62 | -11.82 | 538629 | 0.07 | 0.69 | 0.04 | 0.37 | -2.57 |
| Steepness | 23 | 3128165 | 0.10 | 0.72 | 0.04 | 0.06 | -7.38 | 3128176 | 0.36 | 0.98 | 6.42 | 0.77 | -1.13 |
| Steepness | 3 | 96313248 | 0.04 | 0.78 | 1.71 | 0.32 | -0.10 | 96313248 | 0.08 | 0.75 | 4.16 | 0.50 | -2.49 |
| Steepness | 4A | 10609251 | 0.00 | 0.81 | 0.13 | 0.06 | -2.81 | 10609251 | 0.00 | 0.59 | 0.18 | 0.03 | -2.63 |
| Steepness | 4A | 23031383 | 0.10 | 0.85 | 2.54 | 0.46 | -3.29 | 23031383 | 0.00 | 0.80 | 1.89 | 0.56 | -2.30 |

| | | | | | | | | | | | | | |
|---|---|---|---|---|---|---|---|---|---|---|---|---|---|
| Steepness | 6 | 23280789 | 0.00 | 0.50 | 6.89 | 0.40 | -4.13 | 23280789 | 0.00 | 0.57 | 0.06 | 0.42 | -0.16 |
| Steepness | 6 | 23280789 | 0.00 | 0.50 | 6.89 | 0.40 | -4.13 | 23280791 | 0.00 | 0.57 | 0.07 | 0.34 | -0.19 |
| Steepness | 6 | 23280789 | 0.00 | 0.50 | 6.89 | 0.40 | -4.13 | 23280792 | 0.00 | 0.57 | 0.05 | 0.34 | -0.17 |
| Steepness | 6 | 23280790 | 0.00 | 0.50 | 6.95 | 0.41 | -4.15 | 23280789 | 0.00 | 0.57 | 0.06 | 0.42 | -0.16 |
| Steepness | 6 | 23280790 | 0.00 | 0.50 | 6.95 | 0.41 | -4.15 | 23280791 | 0.00 | 0.57 | 0.07 | 0.34 | -0.19 |
| Steepness | 6 | 23280790 | 0.00 | 0.50 | 6.95 | 0.41 | -4.15 | 23280792 | 0.00 | 0.57 | 0.05 | 0.34 | -0.17 |
| Steepness | 6 | 23280791 | 0.00 | 0.50 | 5.86 | 0.41 | -4.51 | 23280789 | 0.00 | 0.57 | 0.06 | 0.42 | -0.16 |
| Steepness | 6 | 23280791 | 0.00 | 0.50 | 5.86 | 0.41 | -4.51 | 23280791 | 0.00 | 0.57 | 0.07 | 0.34 | -0.19 |
| Steepness | 6 | 23280791 | 0.00 | 0.50 | 5.86 | 0.41 | -4.51 | 23280792 | 0.00 | 0.57 | 0.05 | 0.34 | -0.17 |
| Steepness | 6 | 23280792 | 0.00 | 0.50 | 5.61 | 0.41 | -4.58 | 23280789 | 0.00 | 0.57 | 0.06 | 0.42 | -0.16 |
| Steepness | 6 | 23280792 | 0.00 | 0.50 | 5.61 | 0.41 | -4.58 | 23280791 | 0.00 | 0.57 | 0.07 | 0.34 | -0.19 |
| Steepness | 6 | 23280792 | 0.00 | 0.50 | 5.61 | 0.41 | -4.58 | 23280792 | 0.00 | 0.57 | 0.05 | 0.34 | -0.17 |
| Steepness | 7 | 18124058 | 0.04 | 0.58 | 2.46 | 0.52 | -1.35 | 18144487 | 0.00 | 0.64 | 18.90 | 0.65 | -6.18 |
| Steepness | 7 | 18144487 | 0.00 | 0.58 | 3.40 | 0.46 | -2.83 | 18144487 | 0.00 | 0.64 | 18.90 | 0.65 | -6.18 |
| Steepness | 9 | 12542763 | 0.08 | 0.78 | 3.51 | 0.41 | -4.54 | 12533051 | 0.44 | 0.96 | 9.97 | 0.45 | -4.61 |
| Steepness | Z | 18479119 | 0.04 | 0.78 | 2.10 | 0.70 | -0.80 | 18479119 | 0.00 | 0.74 | 4.63 | 0.51 | -1.86 |
| Steepness | Z | 26653391 | 0.22 | 1.00 | 1.82 | 0.60 | -1.90 | 26653391 | 0.33 | 1.00 | 11.62 | 0.58 | -3.38 |
| Steepness | Z | 47249501 | 0.00 | 0.66 | 1.70 | 0.54 | -1.45 | 47249501 | 0.00 | 0.77 | 0.05 | 0.92 | -0.81 |

Proportions were calculated based on the number of individuals that had the reference or alternative allele at each loci (only loci with allele frequency difference > 0.5 between pure parental individuals were included). AIC difference represent the extent to which a steeper-than-expected or displaced cline provided a better fit than the null model with values fixed at $u = 0.5$ and $v = 1$.

**S16 Table. Results from a genome-wide association study (GWAS) with individuals from all transects combined, to identify regions of the genome associated with rump color (hue).** Allele frequencies, effect sizes, standard errors and P-values for multiple tests (Wald, LTR and Score) are reported for each marker. (Available in dryad repository - DOI: 10.5061/dryad.zkh1893n9).

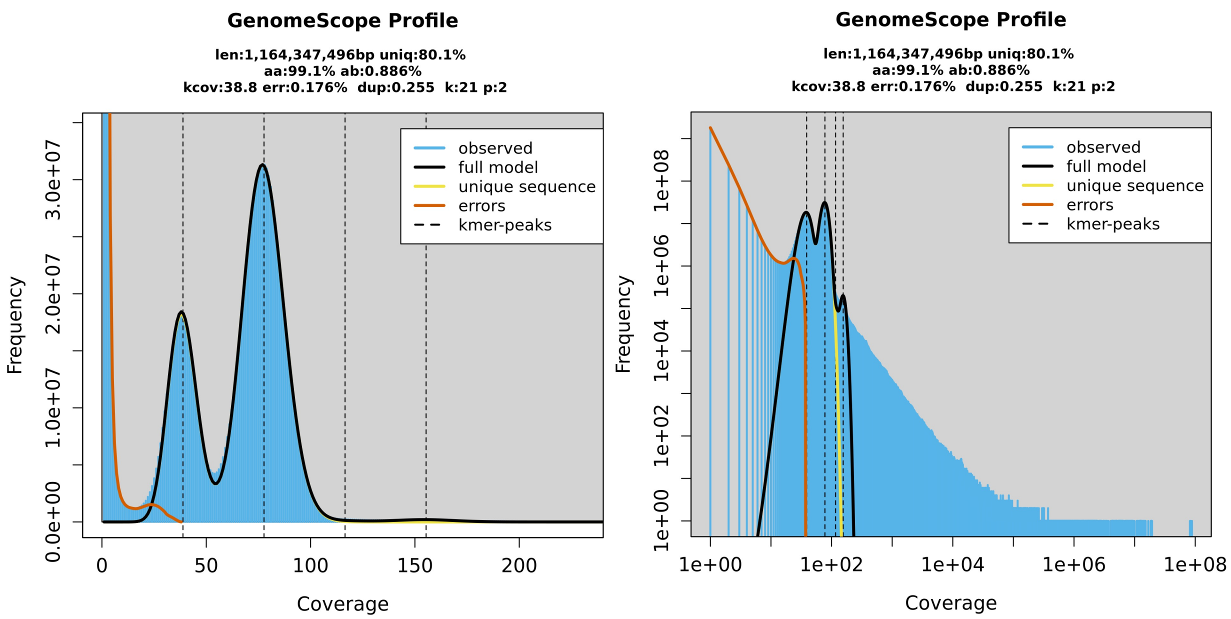

**S1 Fig. Genome size and repeat content estimation using GenomeScope 2.0.** Genomescope2 k-mer (21) distribution from the raw PacBio HiFi reads. K-mer depth is plotted against k-mer frequency for a given depth. The plot displays estimation of genome size (len), percentage of the genome that is not in repetitive elements (uniq), homozygous rate (aa), heterozygous rate (ab) mean k-mer coverage for heterozygous bases (kcov), read error rate (err), average rate of read duplications (dup), k-mer size used on the run (k) and ploidy (p). (a) Expected distribution according to the model. (b) Obtained distribution according to the model. The four peaks correspond to the mean coverage levels of the unique heterozygous, unique homozygous, repetitive heterozygous and repetitive homozygous sequences, respectively.

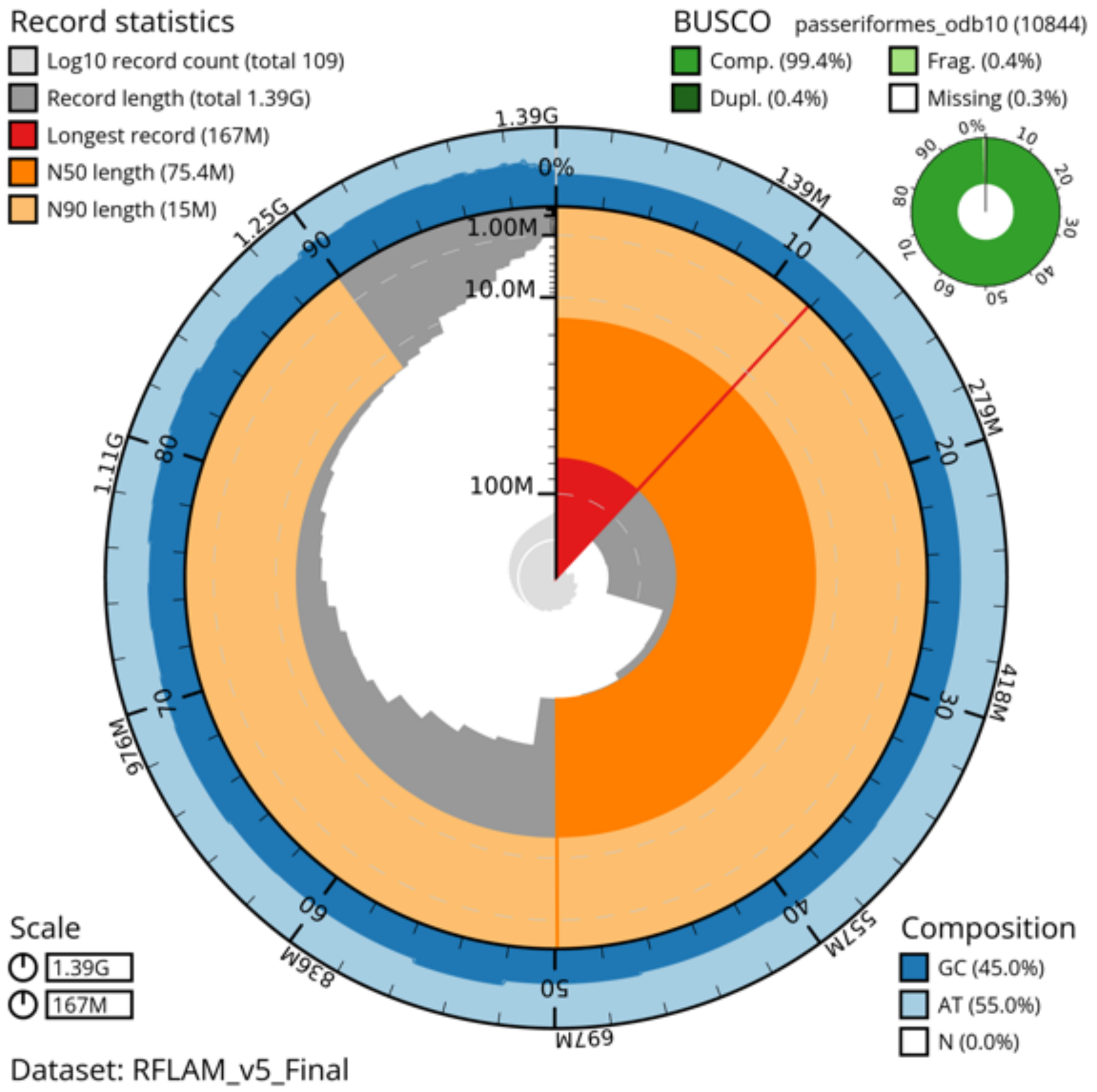

**S2 Fig. Snail plot of the final reference genome assembly.** The contiguity and completeness of the *R. f. icteronotus* genome assembly is plotted as a circle that represents the full length of the assembly (~1.3Gb), assembled in 109 contigs. The N50 (75.4 Mb) is highlighted in dark orange and the N90 (15Mb) in light orange. The longest contig (167 Mb) is highlighted in red. The assembly has uniform GC content of 45.0% except for the last 150 Mb which are unassigned contigs. The BUSCO scores with 99% completeness are shown in the top right corner in green.

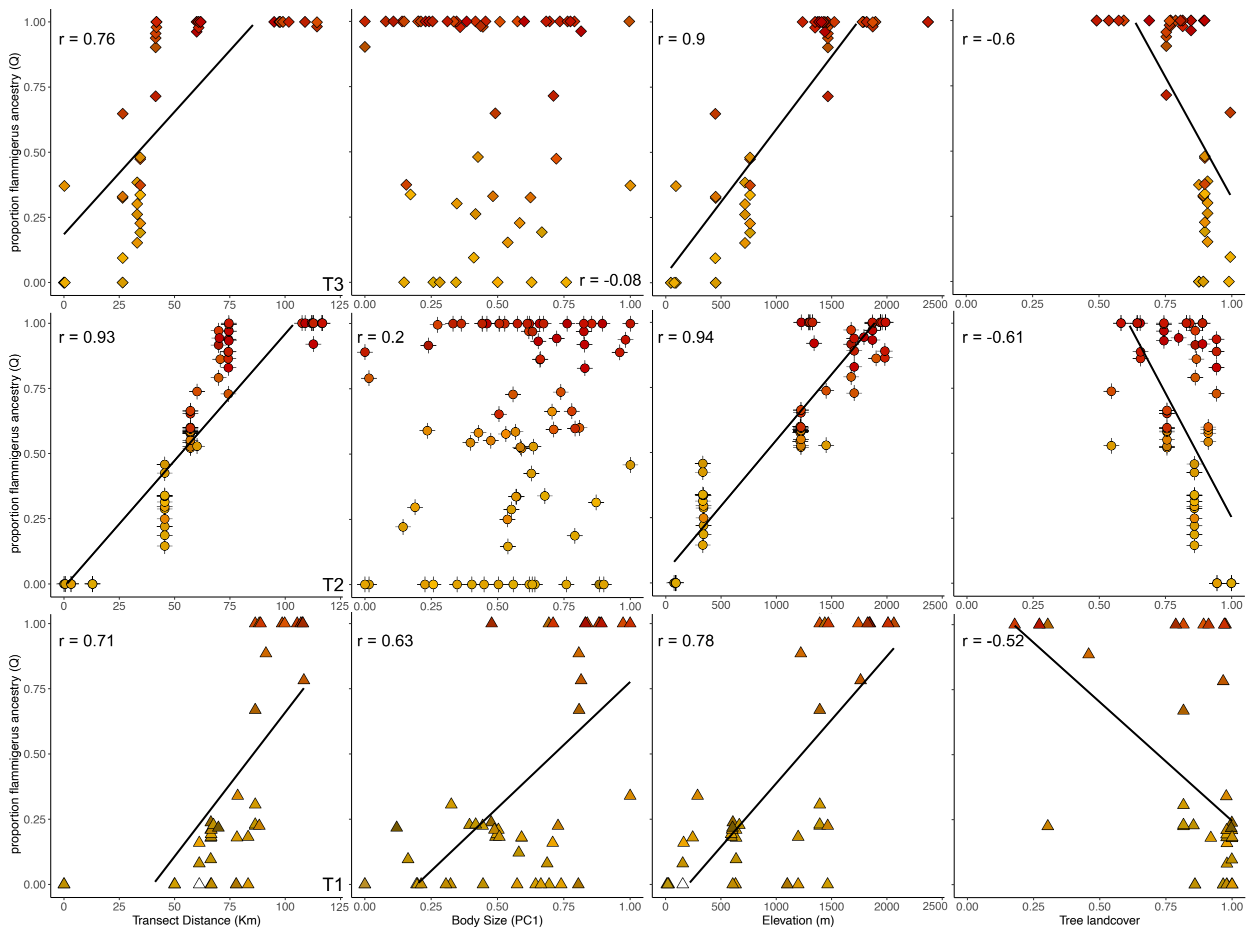

**S3 Fig. Correlations of geographic, environmental and morphological variables with ancestry proportions.** Correlation between hybrid index and A) distance along transect from the westernmost point in km. B) Morphological PC1 as a proxy for body size. C) Elevation in meters. D) tree landcover (%). Points are colored according to their rump plumage color phenotype.

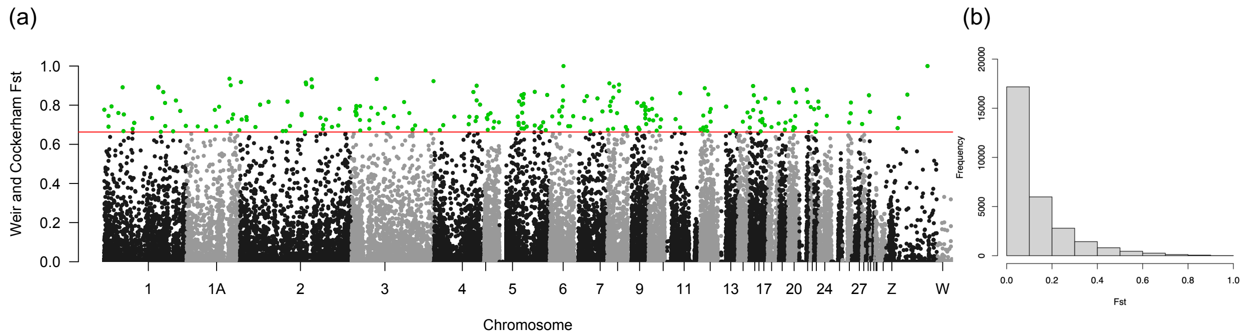

**S4 Fig. $F_{ST}$ manhattan plot and frequency histogram.** A) Weir and Cockerham's $F_{ST}$ between pure allopatric subspecies for 50,868 SNPs (dataset B). We used a 0.995 quantile of $F_{ST}$ (red line) to detect 201 outlier SNPs with high $F_{ST}$ (green). Only two SNPs were fixed between subspecies in chromosome 6 and Z. B) $F_{ST}$ frequency histogram.

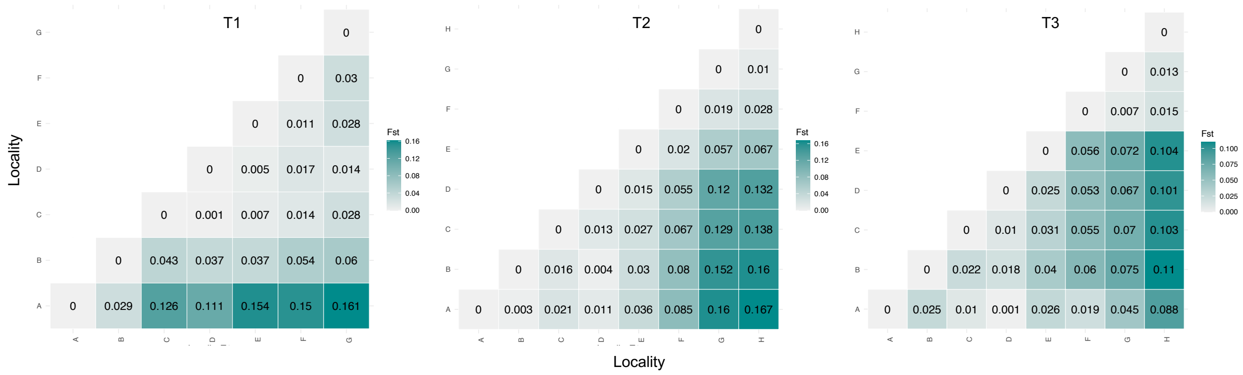

**S5 Fig. Heatmap of pairwise $F_{ST}$ between sampling localities in each transect.** Darker colors represent higher differentiation.

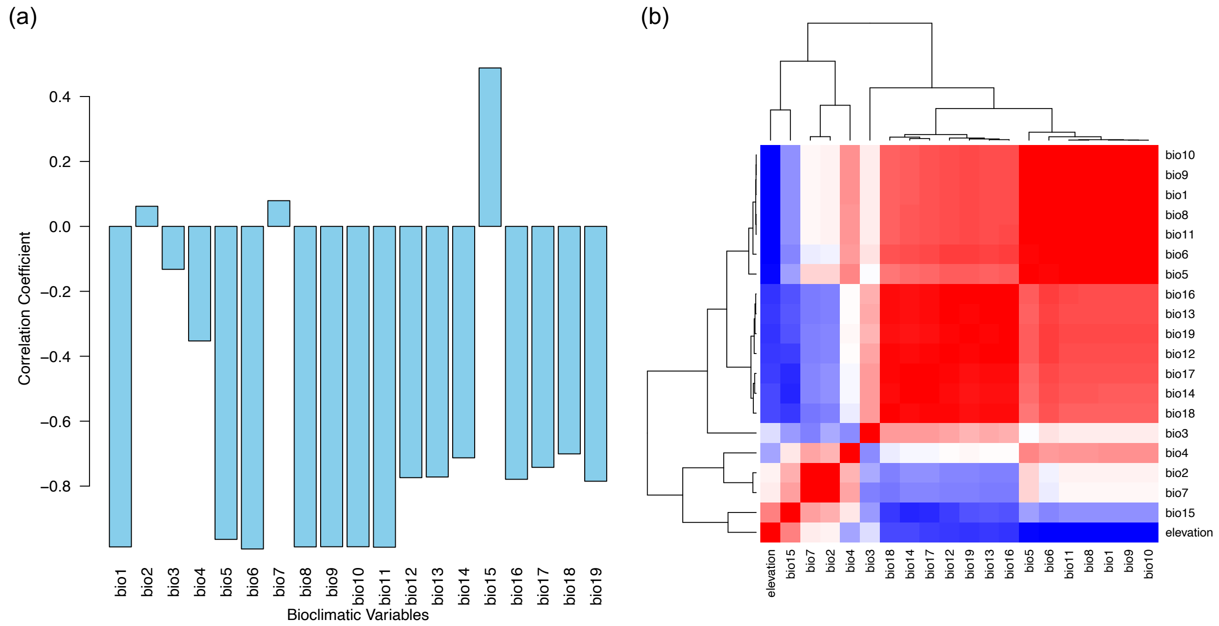

**S6 Fig. Correlation between WorldClim bioclimatic variables and elevation in the sampling area.** A) Correlation coefficient of 19 bioclimatic variables (related to temperature and precipitation) with elevation in our sampling area. B) Correlation heatmap illustrating grouping of variables related to temperature or precipitation and their strong negative correlation with elevation. Color saturation represents strength of the correlation, positive correlations are in red and negative correlations in blue. All correlations were statistically significant.

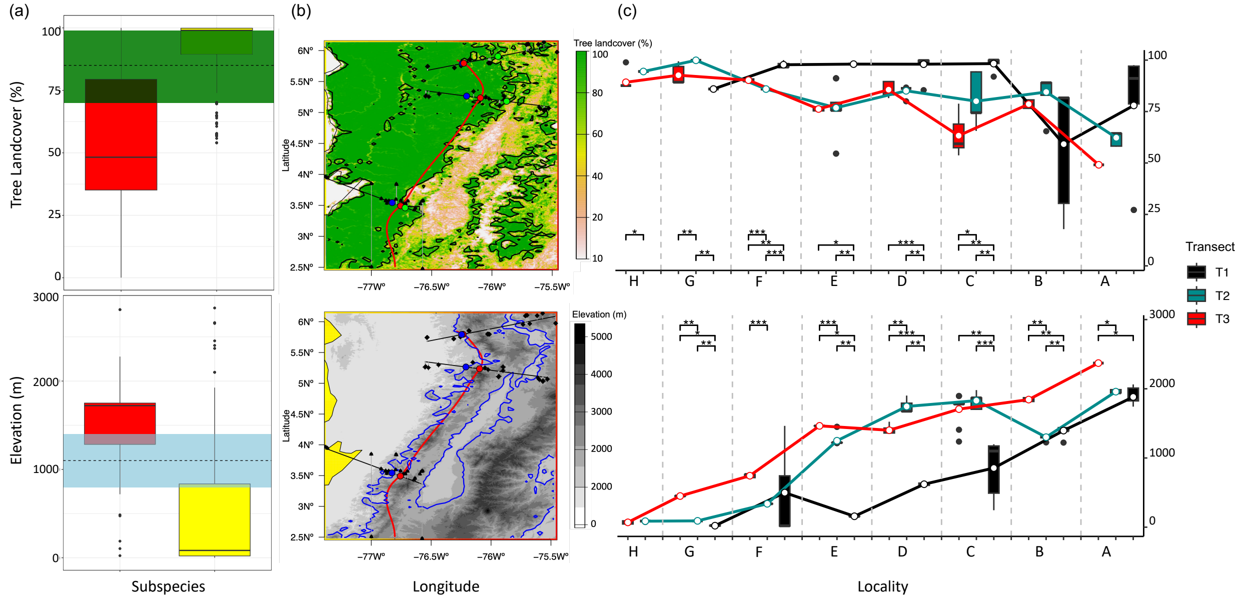

**S7 Fig. Isoclines for environmental transitions and ecological gradients across localities.**
A) Center of the 95% confidence interval of tree landcover (top) and elevation (bottom) overlaps between subspecies. Center of tree landcover overlap (green) occurs at 85%, while center for elevation overlap (light blue) occurs at 1100m (dashed line). B) Red line represents the genetic center of the hybrid zone where the hybrid index = 0.5. Black (top) and blue (bottom) solid lines represent the raster contour lines (isoclines) for environmental transition at 0.85 canopy cover and blue line 1100m elevation. C) Tree landcover (top) and elevation (bottom) across localities in each transect. Lines connect the mean value for each locality along the transects. Degree of statistical significance is plotted above the localities with significantly different mean values between transects.

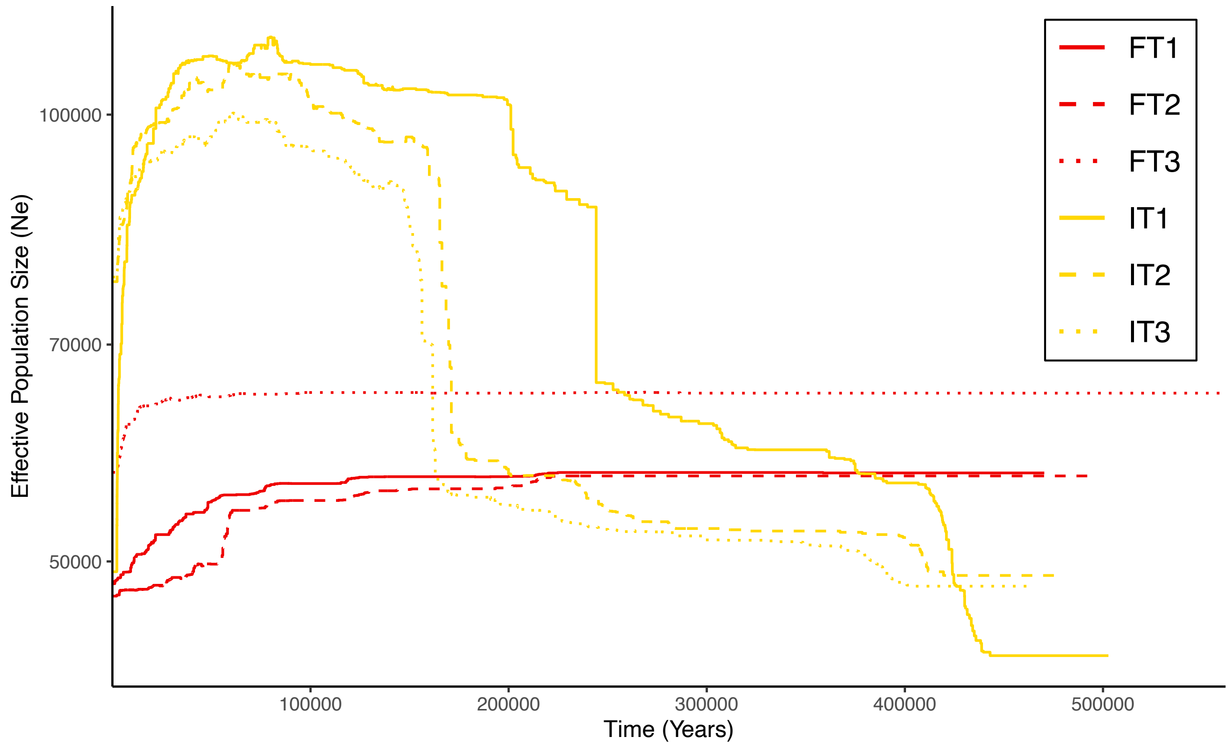

**S8 Fig. Demographic histories of allopatric populations of both subspecies in each transect inferred by Stairway Plot 2.** Simulated assuming a generation length of 2.6 years. Effective population size is in haploid number of individuals.

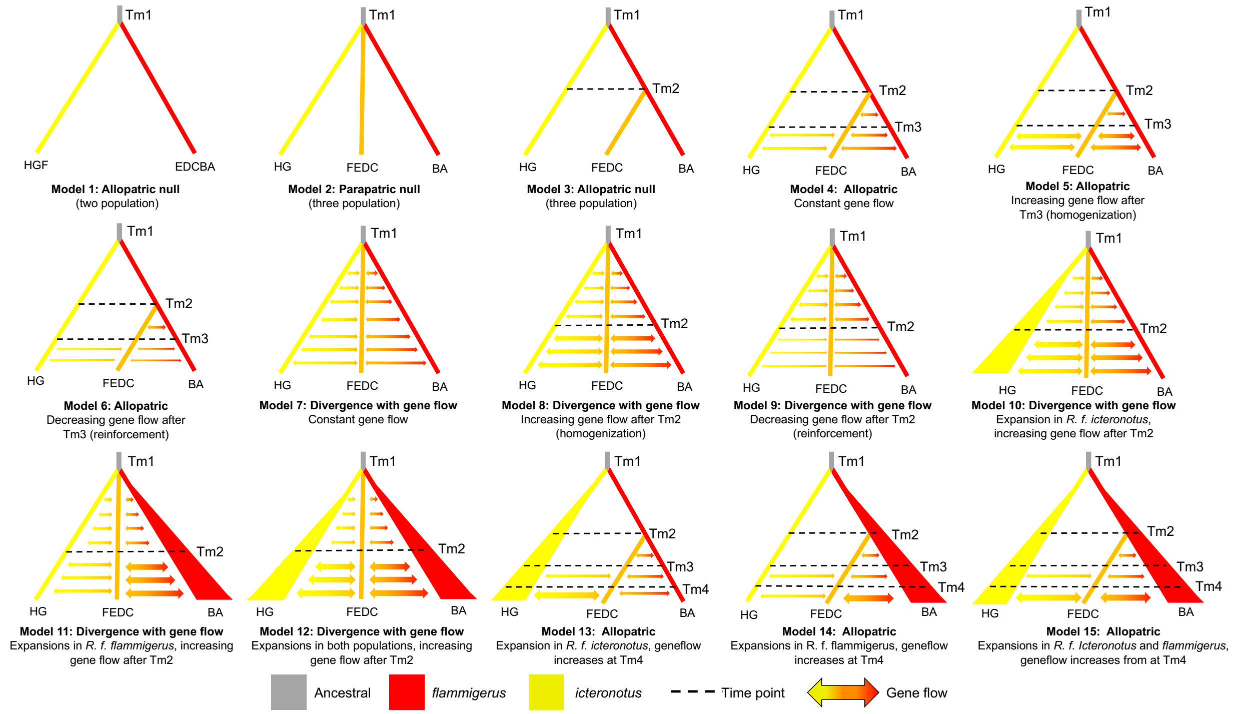

**S9 Fig. Demographic scenarios tested in FastSIMCOAL2.** Red lines represent pure individuals from the allopatric ranges of the *flammigerus* subspecies (localities A, B). Yellow lines represent pure individuals from the allopatric ranges of the *icteronotus* subspecies (localities H, F). Orange lines represent admixed individuals from the hybrid zone (localities C, D, E, F). Grey lines on top of each model represent the ancestral population. In all models time flows from the top to the bottom and arrows represent migration, which was allowed to be asymmetric. The size of the arrows is proportional to the amount of gene-flow in each time period. In all allopatric models Time 1 (Tm1) represents the time of divergence between subspecies. In contrast, because divergence with gene flow models consider gene flow from the moment of population divergence, which affects the coalescent signature (e.g., more recent geneflow pushes forward the time of population split), in all parapatric models Time 1 represents the time of hybrid zone formation. Model 1: Allopatric model of divergence with only two populations and no gene flow after population split. Model 2: Parapatric model of divergence, with a third hybrid population forming at the time of population split but no gene flow thereafter. Model 3: Allopatric model of divergence with no gene flow after population split or after secondary contact. Model 4: Allopatric model of divergence with constant gene flow after secondary contact. Model 5: Allopatric model of divergence with increasing gene flow after secondary contact. Model 6: Allopatric model of divergence with decreasing gene flow after secondary contact. Model 7: Divergence with gene flow, with constant gene flow after population split. Model 8: Divergence with gene flow with increasing gene flow after population split. Model 9: Divergence with gene flow with decreasing gene flow after population split. Model 10: Divergence with gene flow with increasing gene flow after a population expansion in *icteronotus*. Model 11: Divergence with gene flow with increasing gene flow after a population expansion in *flammigerus*. Model 12: Divergence with gene flow with increasing gene flow and population expansion in both subspecies. Model 13: Allopatric model of divergence with increasing gene flow after secondary contact and a population expansion in *icteronotus*. Model 14: Allopatric model of divergence with increasing gene flow after secondary contact and a population expansion in *flammigerus*. Model 15: Allopatric model of divergence with increasing gene flow after secondary contact and population expansion in both subspecies.

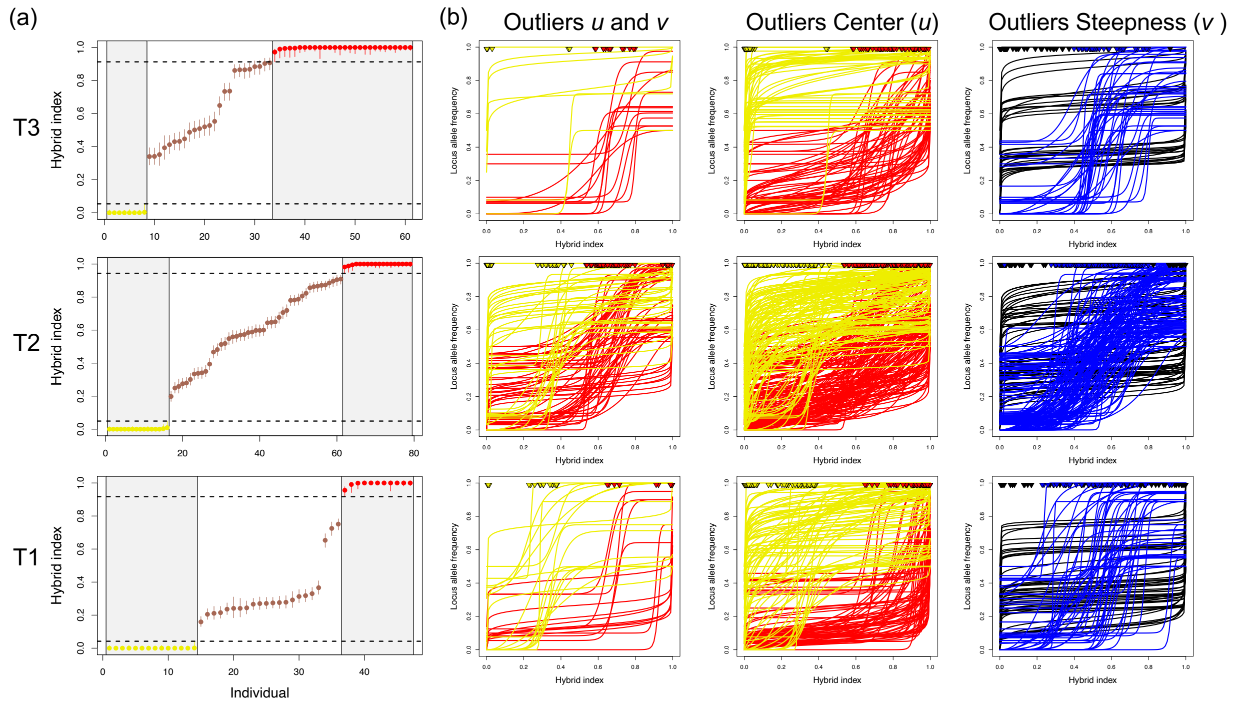

**S10 Fig. Hybrid index and genomic cline outliers estimated with *gghybrid*.** A) Hybrid index calculated for individuals in each transect. Pure *icteronotus* individuals are in yellow and pure *flammigerus* individuals in red and hybrids in dark orange. B) Genomic cline outliers for center (*u*), steepness (*v*) or both (*u* and *v*). Clines in red were significantly displaced into the *flammigerus* background and clines in yellow were significantly displaced into the *icteronotus* background. Blue clines were significantly steeper than the null expectation and black clines were significantly wider than the null expectation. Triangles on top of each plot show the center of each cline.

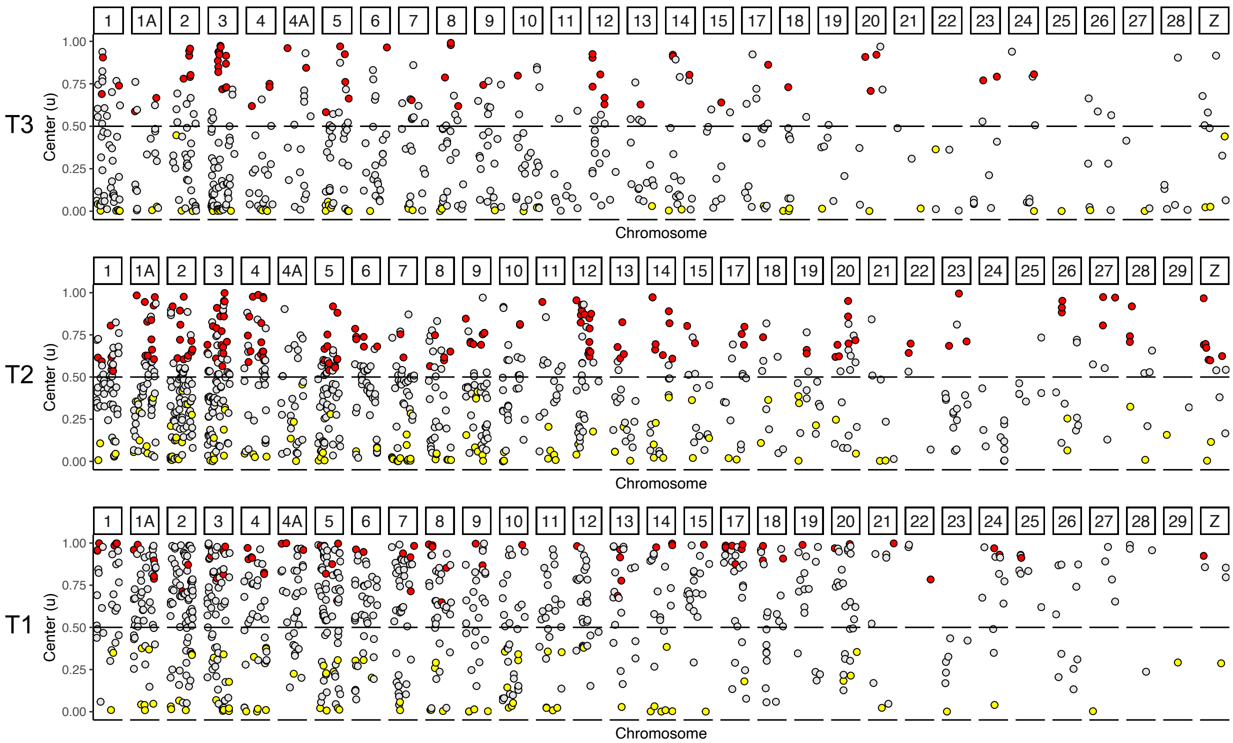

**S11 Fig. Genomic clines center (*u*) across chromosomes**. Cline center outliers that were significantly displaced into the *flammigerus* background ($u > 0.5$) are in red and center outliers significantly displaced into the *icteronotus* background are in yellow ($u < 0.5$). The dashed line represents where the hybrid index = 0.5.

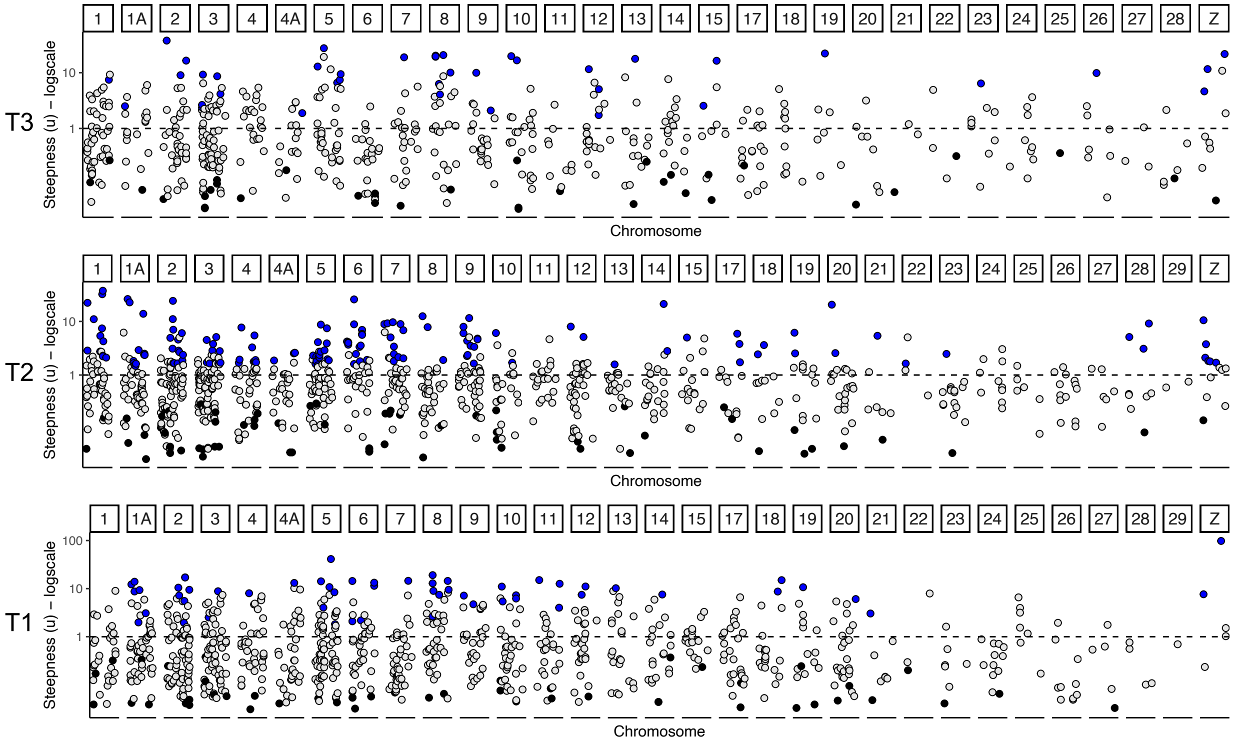

**S12 Fig. Genomic cline steepness (*v*) across chromosomes.** Cline steepness outliers that were significantly narrower than expected are in blue ($v > 1$) and outliers that were significantly wider than expected are in black ($v < 1$). The dashed line represents where $v = 1$ and the y axis is in logscale.

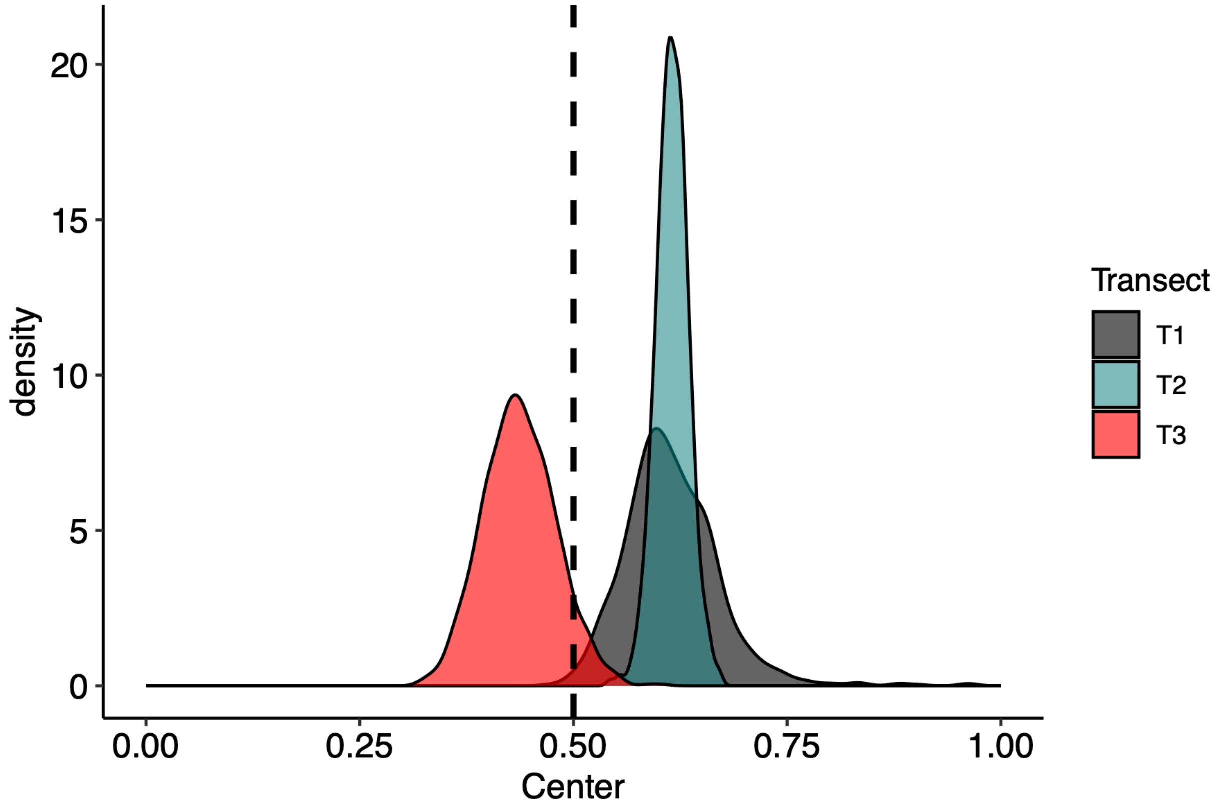

**S13 Fig. Plumage color genomic cline confidence intervals**. Density of center estimates of the plumage color genomic cline estimated with hill functions after bootstrapping. Dashed line represents where the hybrid index = 0.5.

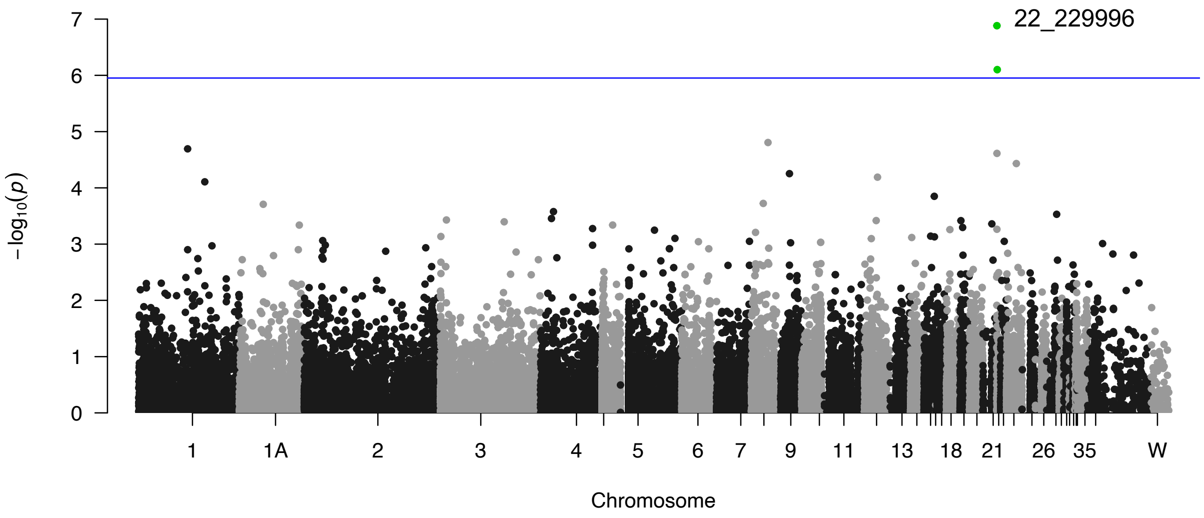

**S14 Fig. GWAS for rump plumage color (hue).** Wald test implemented in GEMMA revealed significant loci (green) in chromosome 22 associated with plumage color. Blue line represents significance threshold calculated with a Bonferroni correction for multiple comparisons accounting for the total number of sites (alpha < $1.115 \times 10^{-6}$).

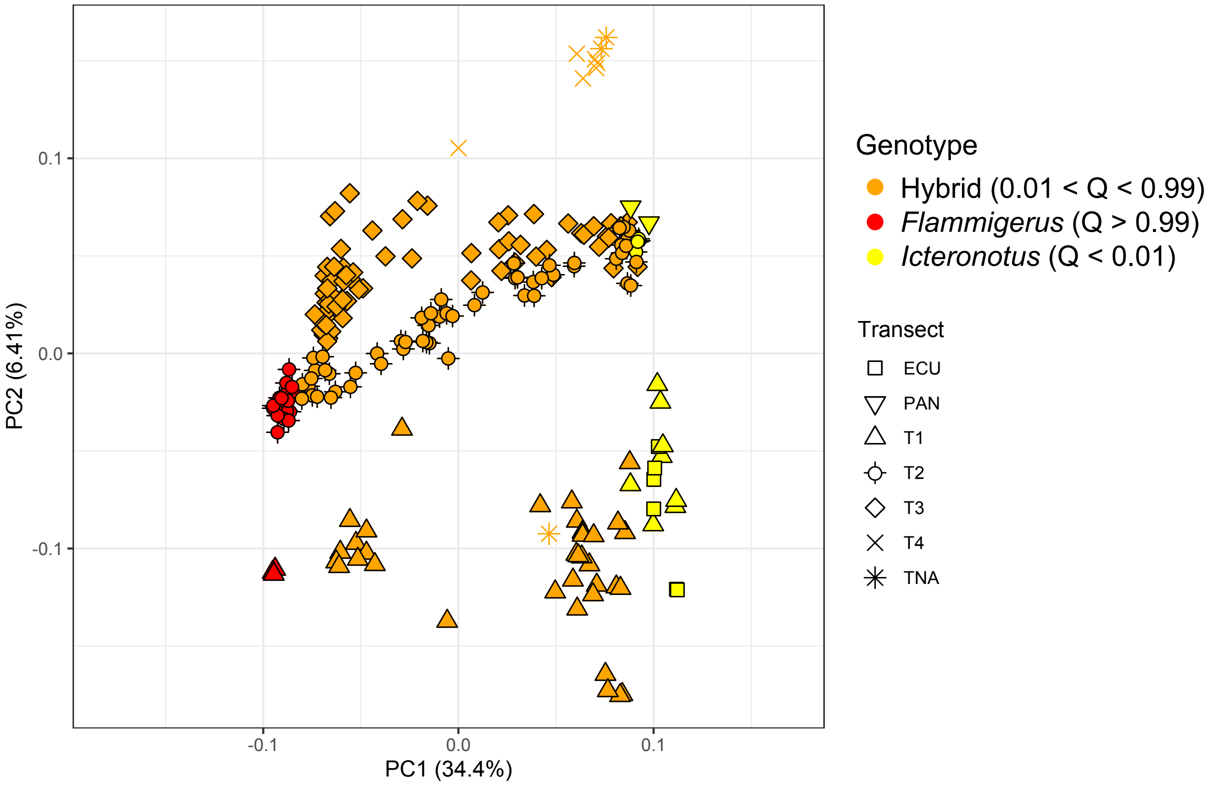

**S15 Fig. Principal components analysis of all available samples.** Genetic PC1 and PC2 of 14,123 genome-wide SNPs including *R.f. icteronotus* samples from Ecuador and Panama. Points are colored according to the ancestry assignment (genotype) for each bird based on ADMIXTURE ran with all samples. TNA indicates samples from individuals in Colombia but far from the sampling transects. PC1 indicates significant genetic differentiation between subspecies and PC2 indicates low population structure consistent with the geographic arrangement of the three transects even when including samples from the full range of *R.f. icteronotus*.
